# The 3D modules of enzyme catalysis: deconstructing active sites into distinct functional entities

**DOI:** 10.1101/2023.06.01.543252

**Authors:** Ioannis G. Riziotis, António J. M. Ribeiro, Neera Borkakoti, Janet M. Thornton

## Abstract

Enzyme catalysis is governed by a limited toolkit of residues and organic or inorganic co-factors. Therefore, it is expected that recurring residue arrangements will be found across the enzyme space, which perform a defined catalytic function, are structurally similar and occur in unrelated enzymes. Leveraging the integrated information in the Mechanism and Catalytic Site Atlas (M-CSA) (enzyme structure, sequence, catalytic residue annotations, catalysed reaction, detailed mechanism description), 3D templates were derived to represent compact groups of catalytic residues. A fuzzy template-template search, allowed us to identify those recurring motifs, which are conserved or convergent, that we define as the “modules of enzyme catalysis”. We show that a large fraction of these modules facilitate binding of metal ions, co-factors and substrates, and are frequently the result of convergent evolution. A smaller number of convergent modules perform a well-defined catalytic role, such as the variants of the catalytic triad (i.e. Ser-His-Asp/Cys-His-Asp) and the saccharide-cleaving Asp/Glu triad. It is also shown that enzymes whose functions have diverged during evolution preserve regions of their active site unaltered, as shown by modules performing similar or identical steps of the catalytic mechanism. We have compiled a comprehensive library of catalytic modules, that characterise a broad spectrum of enzymes. These modules can be used as templates in enzyme design and for better understanding catalysis in 3D.

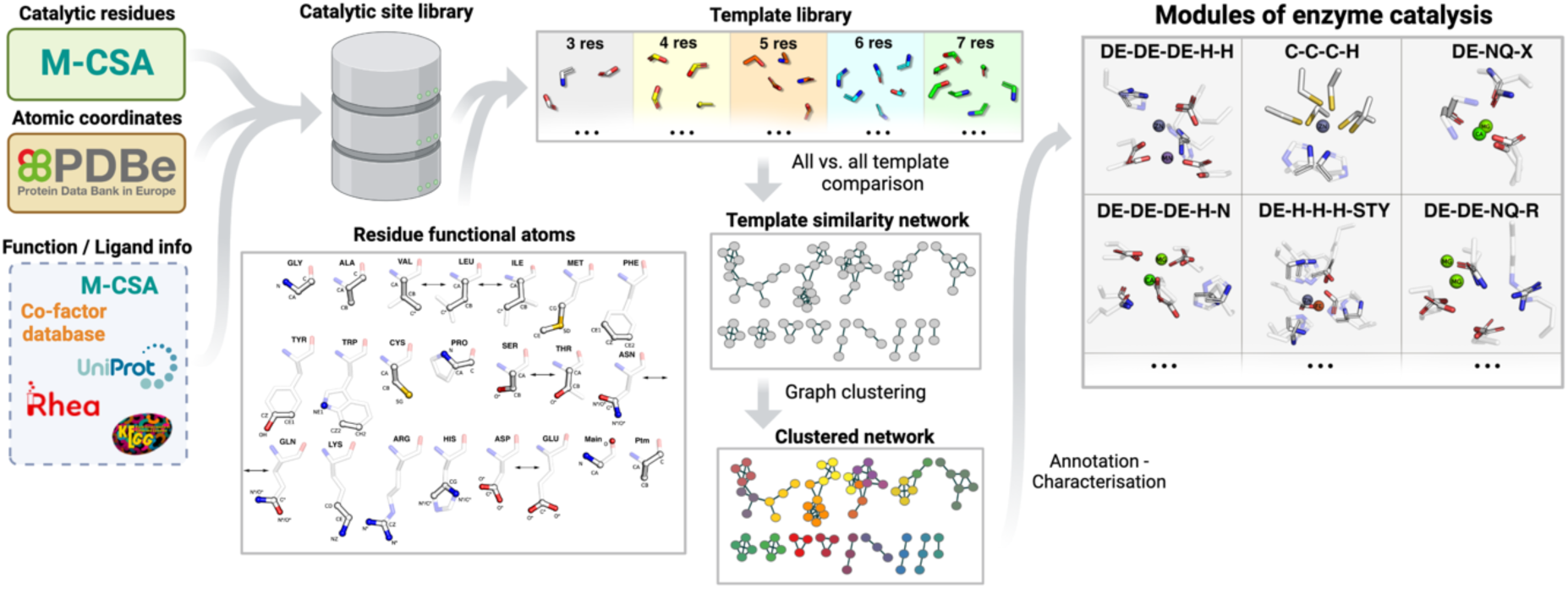

## Introduction

Complex catalytic processes in biological systems are often performed by one or more discrete functional components, comprised of a small number of chemical groups in a defined conformation. Those groups are contributed by basic units, the catalytic residues, that have evolved to be arranged in a defined geometry and are brought into close proximity by the folding of the polypeptide chain. Over the years, our group has been collecting and curating information on the structure and function of enzyme catalytic sites, cataloguing over 1000 enzyme families in the Mechanism and Catalytic Site Atlas (M-CSA)[1,2]. In this database, we have empirically observed that several enzymes in nature adhere to the “discrete functional entities” paradigm, with a simple example shown in Fig. 1. Quinoprotein Glucose Dehydrogenase (M-CSA 104, E.C. 1.1.5.2) is a bacterial oxidoreductase catalysing the oxidation of glucose to gluconolactone with the concurrent reduction of ubiquinone to ubiquinol[3,4]. The enzyme requires pyrroloquinoline quinone (PQQ) and a Ca^2+^ ion as co-factors participating in the oxidation-reduction reaction as well as seven catalytic residues. Even a brief inspection of the catalytic site 3D structure reveals two discrete, compact residue clusters, of three and four residues. The four-residue cluster (Ala293, Tyr295, Asp297, Glu333) binds a Ca^2+^ ion, with the conformation of the residues being driven by organometallic interactions, while the three residue cluster is responsible for the core catalytic process, involving the glucopyranose substrate and the PQQ co-factor. Specifically, a catalytic His168 has the role of the zwitterionic proton donor and acceptor (general acid/base), providing the starting point of a hydride transfer from the C1-OH of the glucose by deprotonation, to the C5 of the PQQ co-factor. The adjacent Asp187 and Arg252 act as activator of His444 and electrostatic stabiliser of negative charge on the PQQ cofactor, respectively. These two entities are simultaneously independent in the 3D space and co-dependent through the course of the reaction (the presence of Ca^2+^ is critical for the activity of the enzyme). Such metal binding sites, which are present in proteins with different folds and functions, are often a result of convergent evolution, with the residue geometry being driven by the coordination sphere of the bound metal. Interestingly in this example, two residues of the metal binding site (Ala293 and Tyr295) ligate the metal through the oxygen atoms of the backbone, and those are the most variable residue positions in the multiple sequence alignment of active sites of the homologous family sequences. This clearly explains the reduced evolutionary pressure in those positions, and the “generic” nature of such motifs within varying structural contexts. Similarly, oxyanion holes in serine proteinases (and other enzymes involving deprotonation of oxygen moieties, usually polymer-cleaving enzymes[5–7]) are formed by two backbone amide groups creating a positively charged pocket that stabilises a negative charge. This is another example of an evolutionarily “generic” motif that has evolved independently in multiple different enzymes. On the other hand, there are residue arrangements that define the specific function of an enzyme and are better conserved and less frequently a result of convergent evolution. Those are usually catalytic sites of enzymes performing complex reactions with several mechanistic steps, involving multiple different functional groups, like in the case of Class A Beta-lactamases (EC 3.5.2.6). Other Beta-lactamases have also converged at the reaction level, with some of which to share similarities in mechanism. A few, for example, use Ser as a nucleophile although surrounding residues are different.

**Fig. 1:**
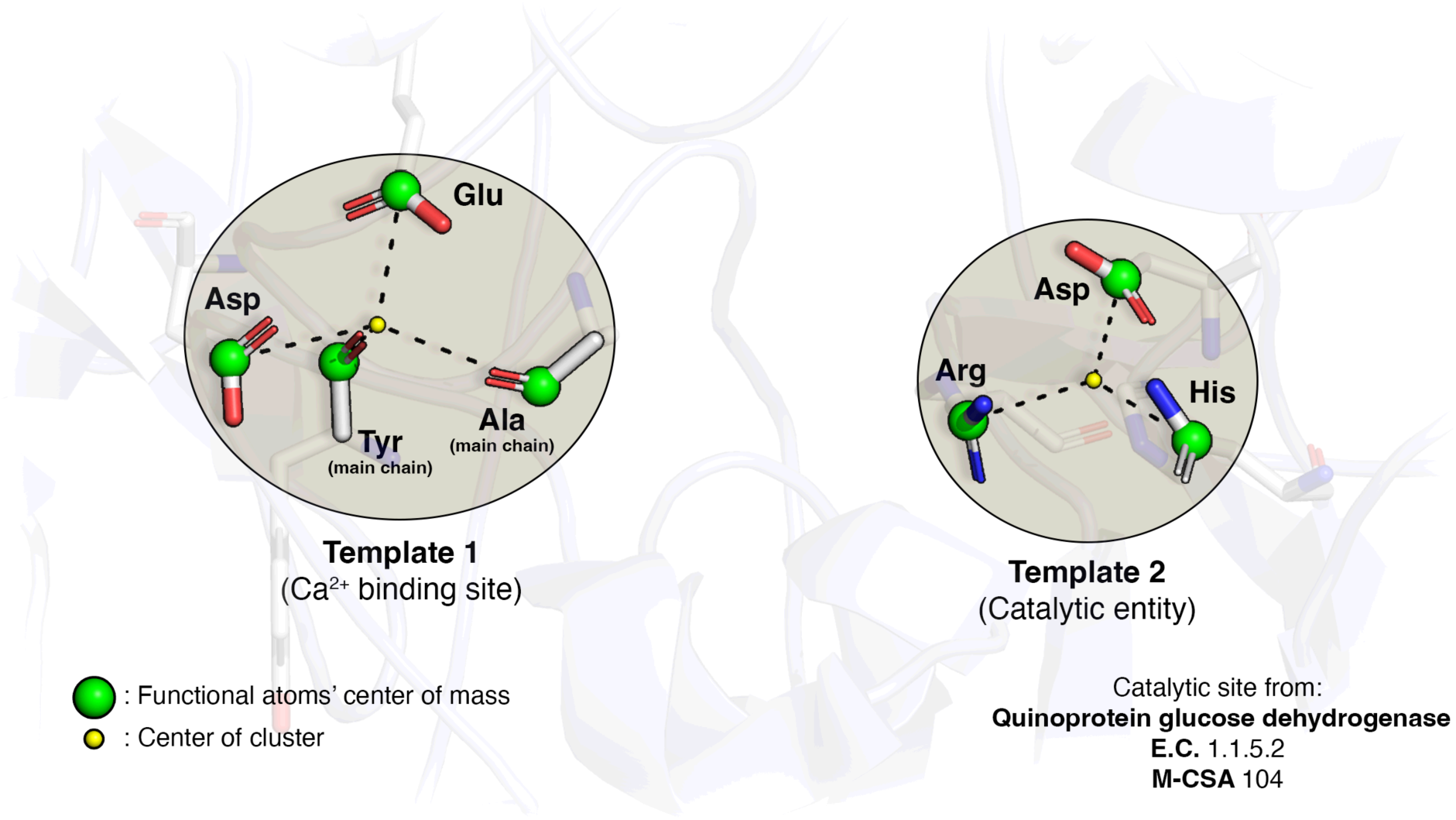
The concept of active site modularity and template fragmentation method. Functional atoms of catalytic residues are shown in sticks (white: C, red: O, blue: N) with their centres of geometry shown as green spheres. Residue clusters (“partial” templates), identified by the k-means algorithm in 3D space, are shaded in grey circles and centres of each cluster are shown as yellow spheres. A partial template of size s consists of the first s residues closest to the cluster centre (dashed lines indicate distances).

Such observations led us to define the concept of the catalytic “module”: a self-contained, compact arrangement of residues, performing one or more tasks in the enzyme catalysis landscape. Each module is named after its residue composition (one letter-code, also accounting for fuzziness – see Methods). Modules can be catalytically active, such as the Arg-His-Asp triad in Quinoprotein Glucose Dehydrogenase, or might be more generic, such as the metal binding site in the same enzyme. Catalytically active modules, however, are not necessarily invariable in terms of 3D conformation and function. For instance, the Arg-His-Asp catalytic triad (the respective module would be named “DE-H-R”, to indicate the presence an acid residue, Asp or Glu) is also present in phospholipase C and performs a different function (phosphodiester bond cleavage)[8] with the 3D arrangement of residues differing too. Furthermore, structural convergence can also apply to catalytically active modules, like the Ser-His-Asp catalytic triad, the most definitive example of active site convergent evolution in enzymology, which is present in multiple analogues of serine proteinases, all independently evolved[9–12].

Here we aim to collect generic and function specific modules that recur in enzymes. These can be derived from enzymes with different folds (convergent) or related enzymes with the same fold but different function (divergent). In addition, we elaborate on the concept of “partial” templates to decompose active sites into compact residue arrangements, identifying recurring instances of those arrangements observed to date, to catalogue, classify and functionally characterise most generic modules known to date governing enzyme catalysis.

## Results

### Definitions

First, it is important to distinguish the definitions for the terms “catalytic site”, “template” and “module” used throughout this paper:

- **Catalytic site**: A collection of three or more catalytically important and conserved residues, manually annotated on enzyme structures in the Mechanism and Catalytic Site Atlas.
- **Template**: An abstract representation of an active site that includes only three functional atoms per catalytic residue. Templates are used as 3D probes to identify similar motifs in a structure, and also to account for matching fuzziness in the atom or residue type. Templates can be “complete” if they represent all catalytic residues of a site, or “partial” if they represent a subset of densely grouped catalytic residues.
- **Module**: A set of recurring partial templates (identical or chemically similar), with defined residue composition and similar geometry, found in at least two evolutionary related (divergent) or unrelated proteins (convergent). A module might contain a mixture of divergent or convergent templates.

### Template library

Using the methods described further below, we derived templates from M-CSA, which contains 1003 enzymes at the time of this study, with 835 functions, as defined by the 4^th^ level of EC nomenclature[1]. The M-CSA was initially constructed to include examples of enzymes that are unique in at least one of the following characteristics: mechanism, overall reaction, or catalytic machinery[13]. With respect to mechanisms, divergent enzymes most often evolve to process different substrates (reflected in a different 4^th^ EC level). Only occasionally do they evolve to perform completely different mechanisms[14]. However, a new entry for a divergent enzyme would not be added in M-CSA if this only differs in substrate specificity, while catalytic residues and mechanism is conserved. The template library was derived to explore evolution of enzyme active sites and mechanisms.

#### Basic statistics of derived templates

Fig. 2a presents some statistics for the template library. In total, we extracted 3953 partial templates (to reflect 813 active sites), providing the principal dataset for all analyses. 3-residue templates are the most abundant (1474), with the number of larger templates decreasing with size (4-residue: 1042, 5-residue: 640, 6-residue: 417, 7-residue: 234, 8-residue: 146). Templates cover all seven EC classes (1: Oxidoreductases, 2: Transferases, 3: Hydrolases, 4: Lyases, 5: Isomerases, 6: Ligases, 7: Translocases), with hydrolases being clearly more numerous, and translocases (only recently added to E.C. nomenclature) occupying only a small fraction. This distribution, reflecting the enzyme functional space covered by UniProt and PDBe. Structural space is also diverse, covering 377 unique superfamilies, as defined at the H level of the CATH domain fold classification[15] and 259 unique topologies (T level). However, two topologies are overrepresented, the Rossmann Fold (3.40.50) and the TIM barrel (3.20.20), highlighting the dense network of evolutionary relationships in enzymes and the relatively limited but functionally diverse structural space[16].

**Fig. 2:**
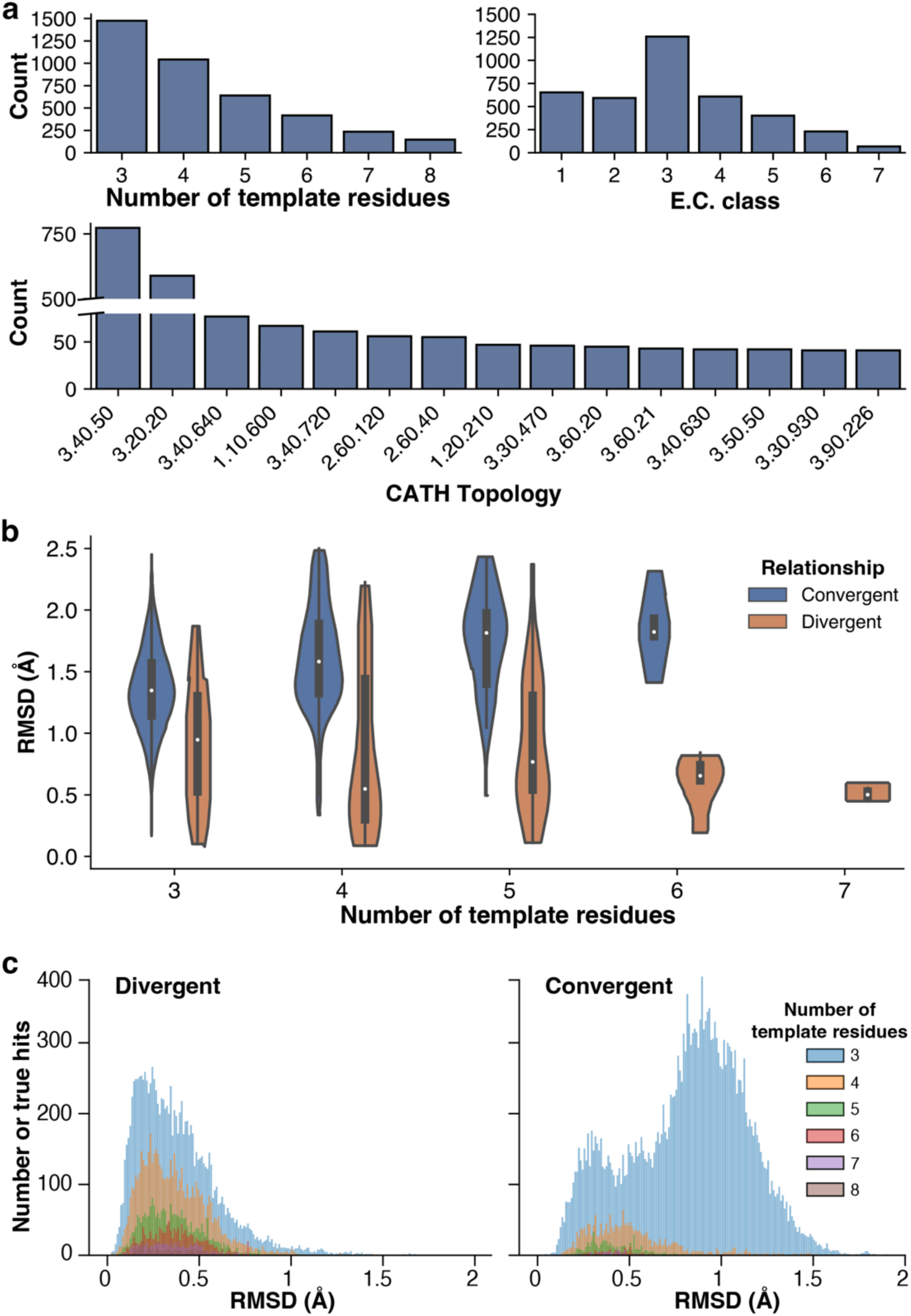
Analyses of the template library (total templates: 3953). a: Basic statistics. The upper left bar plot shows the distribution of template sizes while the upper right plot shows the functional coverage of templates in terms of their EC class (1: Oxidoreductases, 2: Transferases, 3: Hydrolases, 4: Lyases, 5: Isomerases, 6: Ligases, 7: Translocases). The bottom plot shows the structural coverage of the domains from which the templates derive, in terms of the <u>T</u>opology level in the CA<u>T</u>H hierarchy. b: Pairwise template superposition to quantify dissimilarity expressed as the RMSD over functional atoms. Violin plots illustrate the distribution of RMSD values for template pairs of different sizes (3,4,5,6 and 7 residues). Template pairs were split according to whether they derive from domains within the same CATH superfamily (divergent – coloured blue) or different families (convergent – coloured orange). c: Benchmarking of catalytic template search. 6405 non-redundant PDB chains for which catalytic residues are explicitly annotated in M-CSA were scanned by the catalytic template library. Histograms show the distribution of RMSD (Å) on hits containing at least 3 catalytic residues. The left panel corresponds to cases where the query protein is homologous with the protein from which the template has derived (common ancestor, same or divergent function), while the right panel shows cases of hits where template parent protein and query protein are non-homologous (unrelated lineage, convergence in parts of the active site and potentially in function). Histograms are further resolved by the number of residues in the template, shown in different colours.

#### Geometric analysis

To explore the landscape of enzyme active sites, we perfomed an all-by-all comparison of the coordinate templates, using RMSD as the comparison metric. The results are presented in Fig. 2b, where we explore the dependence of RMSD against the template size and the evolutionary relationship of the proteins from which the templates derive. We define a pair of templates as “divergent” if their parent proteins belong to the same CATH superfamily (H level), while in “convergent” templates, parent proteins are unrelated and belong to different CATH superfamilies. Out of 1572 pairs of matched templates, 1419 were convergent and 153 were divergent. It should be commented that those numbers reflect the construction of the M-CSA (see previous paragraph) and not the diversity in nature, where divergence dominates protein evolution (and the emergence of new functions) compared to convergence. Additionally, ∼37% of the template dataset consists of 3-residue templates for which, due to the presence of fewer matching constraints, it is easier to find matches from unrelated proteins. In contrast, larger templates requiring more geometrical constraints, are frequently found in active sites of related proteins. Divergent and convergent templates behave differently when their geometry is compared. In both cases, there is a broad spread of RMSD values. RMSD is sensitive to fluctuations when small coordinate sets (like templates) are compared. It would be expected for RMSD to increase when the template size increases, and this is the case in convergent template pairs, which overall exhibit higher RMSD values than divergent pairs. RMSD is also more variable in 3,4 and 5-residue template pairs, with significantly lower mean values (<1Å) than in convergent pairs, indicating that these functional residues tend to retain a stable geometry during evolution. In 6 and 7-residue divergent templates, RMSD tends to be much lower and inversely proportional with template size, in contrast to convergent templates in which RMSD and size are proportional. This can be explained since the likelihood of matching two templates by chance, within the set pairwise distance cutoff (3.5Å), dramatically decreases with the increase of the template size. Furthermore, RMSD for 4 and 5-residue pairs is distributed bimodally; values are distributed in a region around the mean value (∼0.5Å) and spread up to >2.0Å. The low RMSD region corresponds to nearly identical templates, while higher RMSD could indicate the presence of one or more flexible residues. It has been shown that residues of auxiliary function in catalysis, such as electrostatic stabilisers or ones with a steric role, tend to have more variable geometry compared to those involved directly in the formation/breakage of bonds)[17], therefore, their presence might influence template structural variability.

Templates are extensively used as tools to identify functional sites in protein structures and for functional annotation[12,18–20]. In Fig. 2c we benchmark the ability of our template library to identify catalytic residues and examine the effect of template size and template/query evolutionary relationship on RMSD. 6405 non-redundant chains from the PDB[21], for which catalytic residues are annotated in M-CSA, were scanned by our library of partial templates using the constraint based structural template matching program *Jess*[22]. Searching complete PDB structures means that we find template matches within the active sites, but also distributed around the protein, since some geometrical arrangements of three residues occur quite widely in structures but are not necessarily associated with enzyme mechanism. Such hits are called ‘spurious’ below.

Out of all matches, 4768 (3-residue: 2413, 4-residue: 1246, 5-residue: 563, 6-residue: 315, 7-residue: 155, 8-residue: 76) were divergent (template and target structure from the same superfamily) and 7223 (3-residue: 6477, 4-residue: 524, 5-residue: 133, 6-residue: 51, 7-residue: 30, 8-residue: 8) were convergent (different superfamilies). Unsurprisingly, matches where template parent protein and query protein are homologous (divergent) exhibit much lower RMSD values than convergent matches. This is useful when assessing the quality of template-template matches, indicating that convergent template geometries will be expected to be less similar than divergent ones. Also, template size is shown to be a definitive factor affecting template structural similarity. While in divergent matches, the shape of the RMSD distributions is retained when the overall number of hits increases with decrease in template size, in convergent examples, 3-residue matches are much more wide-spread, in a bimodal distribution, reflecting spurious hits. The global maximum of the histogram lies at ∼1Å, therefore, 3-residue convergent matches, either between templates and queries or between two templates, will be expected to exhibit higher RMSD values, with a higher likelihood of being spurious compared to matches from larger templates.

However, the chance of identifying a hit by chance drops as the query size is smaller; in the case of module identification discussed below, the queries are templates themselves, therefore the chance of two templates being structurally similar by chance (and not being functionally relevant) is relatively low.

### The modules of catalysis

#### Overview

We used the template cross comparison analysis to capture templates that recurred in more than one protein superfamily and to construct enzyme modules. We found the following matches, as illustrated in Fig. 3. 706 out of 3953 partial templates (3-residue: 579, 4-residue: 215, 5-residue: 69, 6-residue: 17, 7-residue: 4, 8-residue: 0) were grouped into 161 modules. Out of those, 87 modules were 3-residue, 44 were 4-residue, 22 were 5-residue, 6 were 6-residue and 2 were 7-residue. Fig. 3 represents the modules as a network of template-template relationships, where each node represents one or more templates from the same CATH homologous superfamily. The connections represent similarities with RMSD ≤ 2.5Å. The number of nodes in one module, represents the number of distinct superfamilies in which that module is found. As observed in the case of the Arg-His-Asp triad, the geometry of templates can vary within a module and multiple conformers could form structural clusters. Graph clustering allowed us to identify those clusters, using as a metric the connectivity (modularity) of each putative cluster. As clearly seen in Fig. 3, members of the same cluster tend to be interconnected (e.g. the two large clusters of the DE-DE-H module), but not necessarily fully connected (e.g. DE-H-NQ module).

**Fig. 3:**
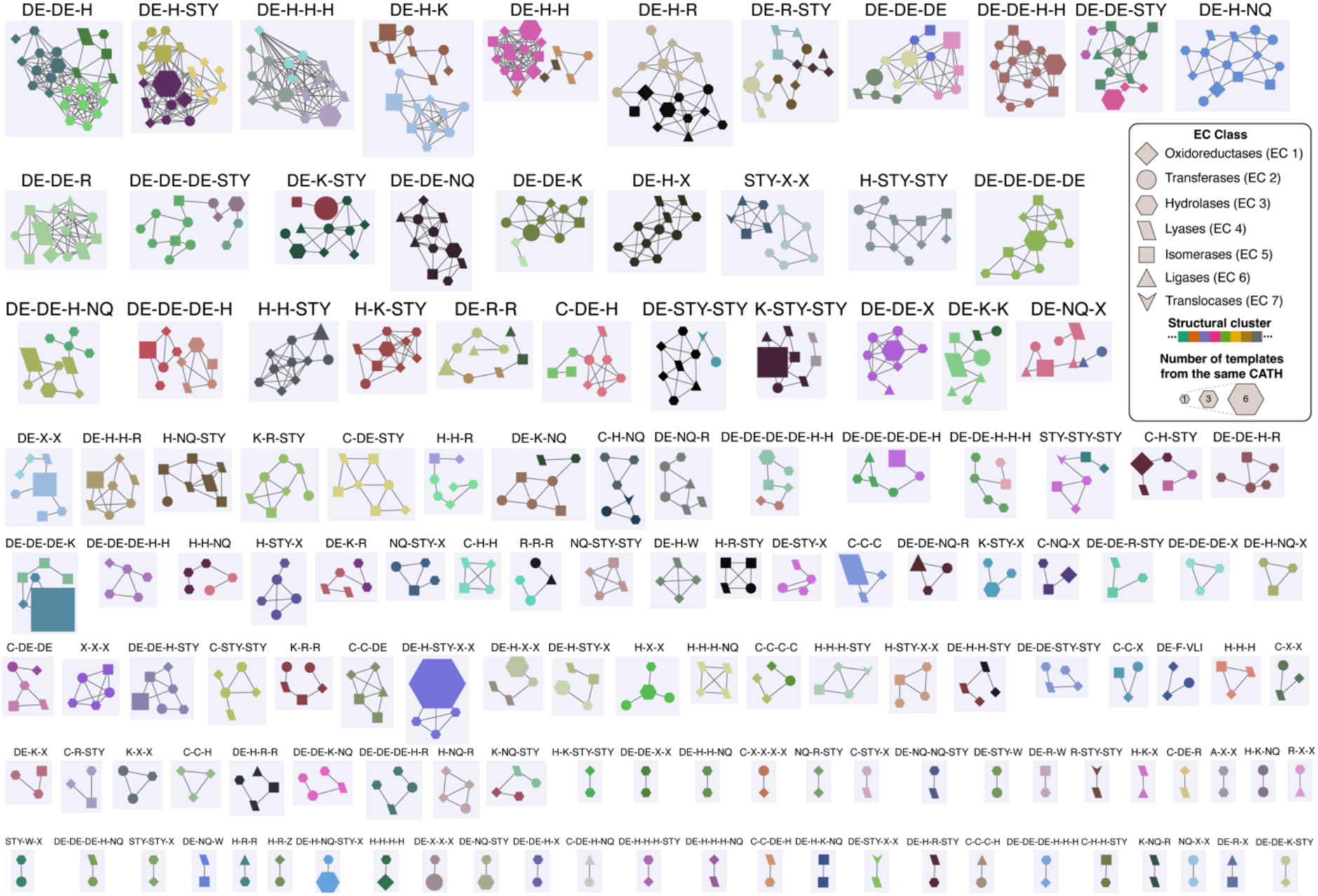
Graph representation of the modules landscape. Each node represents a template or a group of templates from the same CATH domain (size proportional to the number of merged nodes). Each subgraph represents a “module” named after its residue composition, where similar templates (RMSD ≤ 2.5Å) are connected by an edge. Different structural clusters within a module are indicated by a different arbitrary colour. Different node shapes correspond to different EC classes. For better visualisation, only modules with two or more templates from a different CATH domain are shown. For better visualisation and to reduce clutter, a sample of 300 random templates within the modules set was collected to draw the figure. Graph construction and clustering (Leiden algorithm) were performed in Cytoscape[23].

To visualise and illustrate the landscape of the modules of catalysis. Fig. 4 shows a hierarchical pie chart with five levels of classification: 1) Module size (number of catalytic residues), 2) fuzzy residue composition (the primary “module” level), 3) structural cluster 4) CATH homologous superfamily and 5) EC number of hosting enzyme. Structural variability within structural clusters is expressed by the average RMSD of the templates to a cluster representative.

**Fig. 4:**
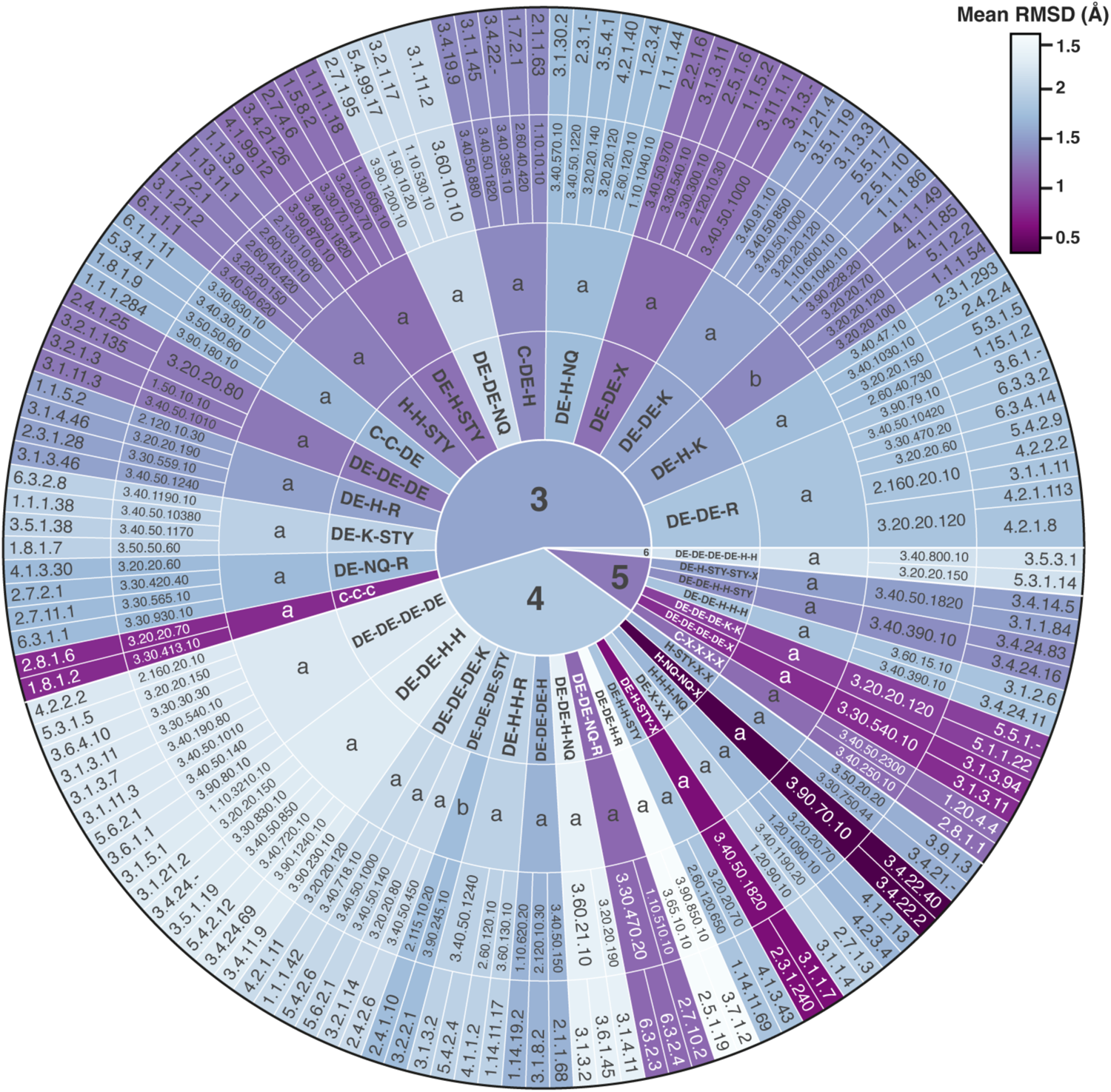
Classification of the modules. Levels of classification are represented as concentric circles, from the centre outwards, as follows: 1) Number of residues 2) Residue composition 3) Structural cluster 4) CATH superfamily 5) EC code. Each classification group is coloured according to the mean RMSD of its corresponding conformational clusters, quantifying the structural variability within. Lower RMSD values (darker colours) indicate relatively invariable motifs in 3D, while higher RMSD values (lighter colours) indicate variable motifs.

#### Divergent and convergent evolution in modules

It is important to discuss the structural and functional characteristics of modules in the context of divergent and convergent evolution. We find that most modules include both divergent and convergent templates, with their ratio (# divergent/# convergent) increasing with the number of residues. This is visible in Fig. 4, where four out of seven 5-residue modules in the figure derive from divergent proteins (same CATH superfamily). Divergence and convergence are also reflected in the graph landscape in Fig. 3. Several examples (DE-H-STY-X-X, DE-DE-DE-K, C-C-C, K-STY-STY, DE-DE-H-NQ) contain several divergent templates (larger nodes), as well as convergent ones. The question is whether the functional role of these modules is retained when enzymes diverge.

Table S2 is a collection of the divergent subset of templates in the module library (43 modules), grouped by CATH superfamily (52 groups). Most of them contain only a single superfamily group, where templates are likely to perform a similar role. For instance, the two templates from divergent enzymes in the C-NQ-X module (CATH 3.60.20.10), correspond to a triad of a catalytic nucleophile (Cys) and two oxyanion hole residues (Asn and Gly (backbone)). The two enzymes containing this module, Glutamate Synthase (EC: 1.4.7.1) and Amidophosphoribosyltransferase (EC: 2.4.2.14), catalyse different overall reactions. However, both retain part of the mechanism conserved (hydrolysis of glutamine to glutamate) using the same conserved triad, with the latter having an extra, coupled functionality facilitated by a separate group of catalytic residues[24,25]. This is likely a gain-of-function evolutionary scenario, where part of the active site remains conserved for maintaining a particular part of the mechanism, and an emerging new function utilises this site as a component.

Some modules in Table S2 contain multiple CATH superfamily groups. In those cases, residue function might be similar both within and across superfamilies, or similar within one superfamily but different across superfamilies. The first case is exemplified by the DE-DE-DE-H module, found in enzymes from two superfamilies (3.20.20.120 and 3.30.420.10). This is an acid residue triad binding Mg^2+^ and Mn^2+^ ions accompanied by a zwitterionic His acting as a proton shuttle[26–30]. The second scenario is represented by the DE-K-STY module. Here, five enzymes are grouped into two superfamilies, 3.40.50.300 and 3.40.640.10. In the first group, the module is a group of transition state electrostatic stabilisers[31,32], while in the second, it interacts with a PLP co-factor[33–35]. Many papers have been published discussing divergent evolution[36,37] and this dominates the emergence of new functions.

For the modules suggesting convergent evolution (131 or 83% of a total of 161 modules) we analysed whether structural convergence is accompanied by functional convergence or whether the change of EC class observed in proteins containing these modules is coincidental. This is the ultimate question addressed in this paper, with mostly convergent templates being under the spotlight.

#### Structural characterisation

We explored structural and functional diversity of the modules, by looking at them individually. Fig. 5a shows the number of unique CATH superfamilies (H level, left panel) and unique EC numbers (to the 4^th^ EC level, right panel) associated with each of the first 100 most populated modules. Most modules are highly diverse both structurally and functionally, with the numbers of unique domains and ECs being correlated (i.e. structure-function correlation). However, several do not to follow this trend; for instance, the second most abundant module (DE-H-STY, otherwise known as the Ser-His-Asp catalytic triad of serine proteinases), is seen in ∼30 different enzymes (21 unique EC subsubclasses), whose structural space covers ∼20 structural folds. This underlines the high functional diversity of this module as a catalytic entity[9,38–40]. Functional diversity is also visible when looking at the extended module DE-H-STY-X-X which includes the two residues forming the oxyanion hole using their backbone atoms. Similarly, generic modules that bind metal ions, for instance DE-DE-DE-K and K-STY-STY, have also remained structurally unaltered while enzyme functions diverged. Those two perfectly exemplify the evolutionary constraints placed by co-factor binding on catalytic residue conformation. The former module is almost always seen to bind Mg^2+^ or Mn^2+^ ions; three acidic residues (Asp or Glu) in any combination are directly coordinated by the metal, while an amide group contributed by an adjacent Lys residue, acts as a stabiliser of negative charges during this protein/ligand interaction. The need to bind organic co-factors might also lead to convergence during evolution, like in the K-STY-STY module that is observed in crystal structures to interact with NAD and its analogues, making it one of the few examples in our dataset where a consistent residue geometry is coordinated by a flexible organic co-factor.

**Fig. 5:**
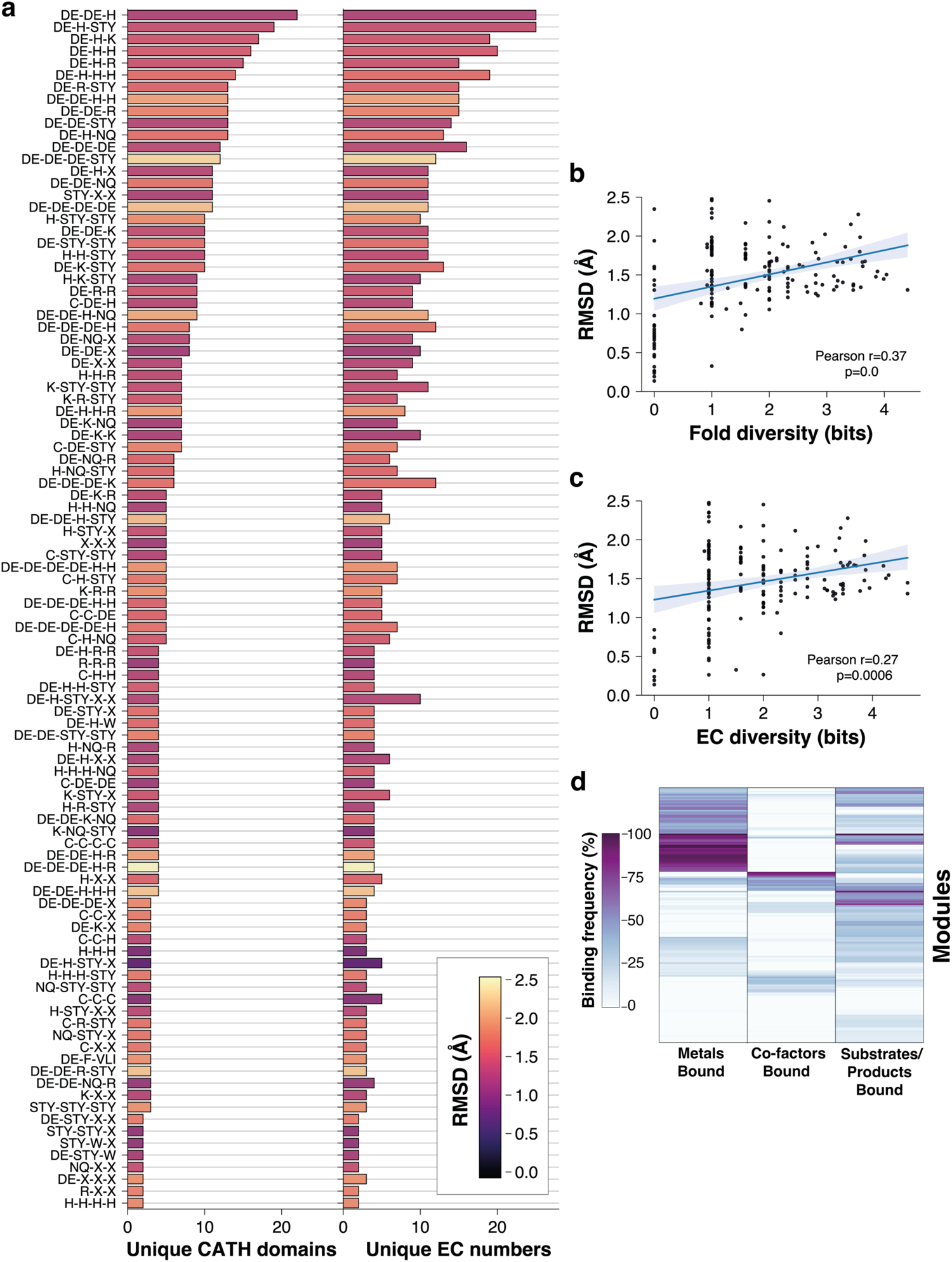
Structural characterisation of catalytic modules. a: Structural and functional diversity of the proteins accommodating the templates within each module. Diversity is expressed as the unique number of CATH domains and EC numbers respectively. To reduce clutter, only modules of ≥3 templates are shown. Average RMSD of all templates against a reference template is shown in colour scale. b and c: Average RMSD against the reference template for each module, plotted against the structural diversity (b) and functional diversity (c), expressed as the Shannon entropy of CATH and EC numbers respectively. Each point corresponds to a template and linear regression is drawn as a blue line with the respective confidence interval shaded. d: Heatmap of the bound ligand binding frequency scores (x-axis) for each module (y-axis).

Fig. 5b and c demonstrate the relationship between template conformational variation within modules (average RMSD) and diversity of the proteins from which the constituent templates derive from, in terms of structural folds (CATH domain diversity) and function (EC number diversity) respectively. In both cases, when no structure/function diversity is present, template conformational variation tends to be low. This is evident among enzymes of identical function (zero EC diversity), where average RMSD is always below 1Å. In cases of enzymes belonging to the same CATH superfamily (zero fold diversity), the average RMSD is slightly more variable, while most modules stay below 1Å. This behaviour is explained by the fact that proteins with an evolutionary link, sharing a similar overall fold, have diverged into enzymes of discrete function, where some key structural elements of their active site have been conserved in 3D, either perfectly (RMSD < 1Å) or partially (1Å < RMSD < 2Å). When diversity increases, in both structure and function, a subtle linear trend with RMSD is observed (Pearson correlation coefficient: 0.37 and 0.27 respectively). This highlights that templates with a defined function are either well conserved or have converged to a similar conformation (the geometry of functional atoms needs to be preserved in order for the chemistry to happen), and their structural evolution is relatively independent from the evolution of the overall protein fold.

A caveat should be stated here: By definition, modules only contain templates sharing the same fuzzy residue composition. This captures functional changes for an arrangement of conserved or conservatively mutated residues. It does not capture changes caused by non-conservative mutations (e.g. mutation of a charged catalytic residue to a hydrophobic one). These non-conservative mutations of active site residues often lead to change[41] or even loss of function[42,43].

#### Functional characterisation

Templates and modules have been identified with the exclusive use of geometrical and knowledge based criteria, so that further scrutiny is required to establish any potential relationship between module geometry (structure) and mechanism of action (function). Here, three approaches to achieve this are discussed (details in Methods section). The first is based on investigating which type of ligands lie in proximity to each template. The second is the enrichment of each residue position with functional role annotations and their frequencies, taken from the M-CSA entry for that enzyme. The third is the identification of mechanistic steps in which template residues participate, to search for common instances within a module. Exemplary 3D models of convergent modules are shown as superimposed templates in Fig. 6, while comprehensive module descriptions for the whole library are presented in Table S3. This is populated with examples of enzymes and folds represented by a module, the structural variation (average RMSD) between members of the module, annotations of ligands bound in proximity and catalytic residue functional roles, as discussed below.

**Fig. 6:**
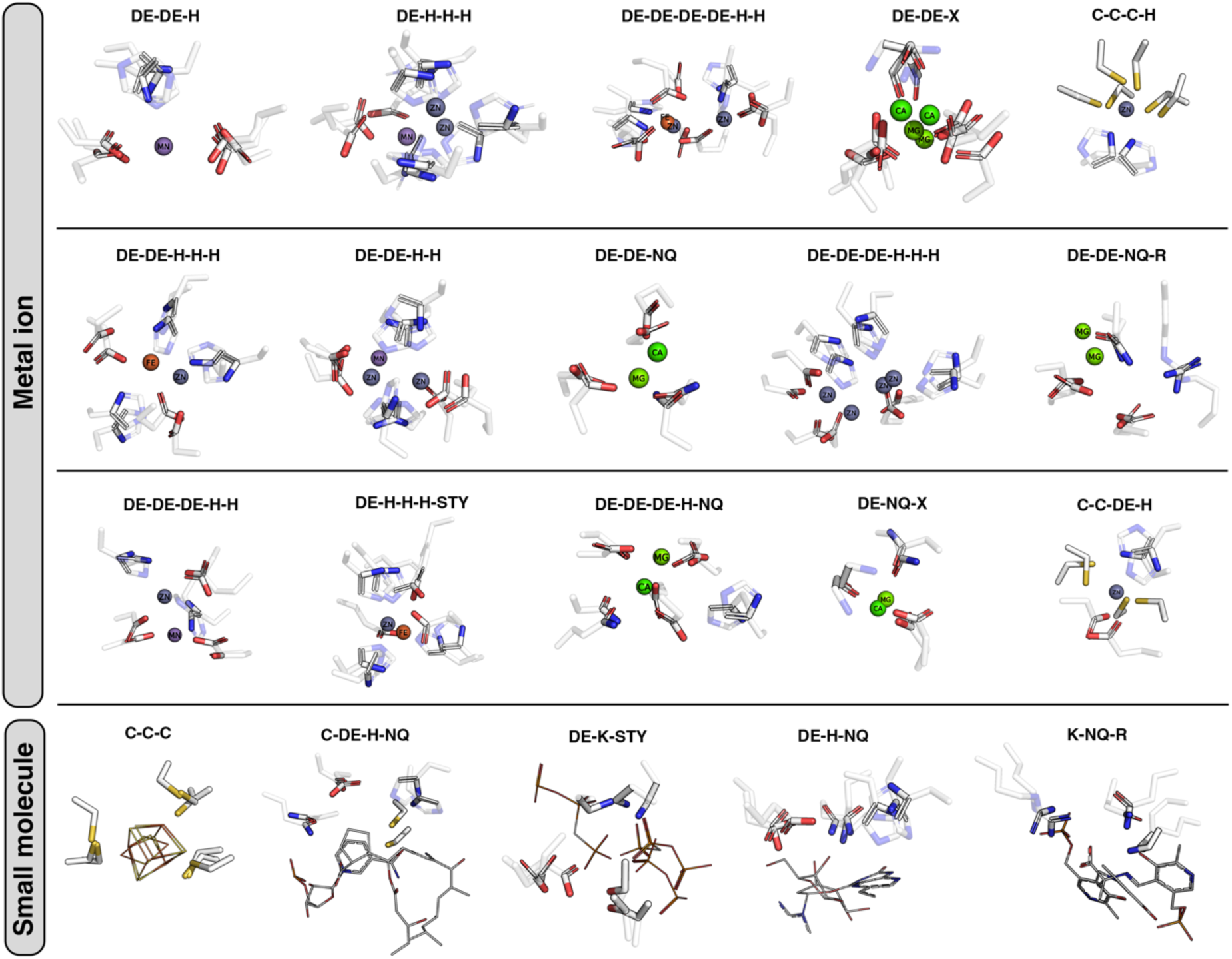
A collection of example modules, shown as superimposed convergent templates. Catalytic residues are shown as opaque and transparent sticks, indicating functional and non-functional atoms. For residues contributing side-chain atoms, backbone is removed, whereas for backbone contributing residues, side chain is removed. Ligands are shown as spheres (metals) and lines (small molecules). Atoms are coloured according to their type (white: C, red: O, blue: N, yellow: S, orange: P). Bulky small molecules were truncated to reduce clutter. 3D models were prepared in PyMol[49].

#### Modules can be characterised by bound ligand preferences

The first approach for functional characterisation explores the idea that a conserved/convergent arrangement of catalytic residues will be associated with a set of ligands with which it interacts (as taken from PDB structures for members of the module). For this, each module was assigned with a binding frequency score for each ligand category (substrate/product[44], co-factor[45], metal ion[46,47]) that approximates the propensity to bind a ligand falling in those categories. Those scores are presented as a heatmap in Fig. 5d. Almost all modules documented in Table S3 appear to interact with a ligand (given the distance cutoff we set). Of these, the majority involve metal binding, fewer bind cofactors and even fewer bind substrates.

At a glance, ∼1/3 of the modules bind a metal with 25% frequency or more. On the other hand, only a few modules seem to interact with organic co-factors, with respective propensities not exceeding 50% in most cases. This is surprising, since “generic” compounds like organic co-factors, are found across a plethora of domains and functions and would be expected to interact with a limited variety of local 3D motifs. However, this is not the case, as shown in the examples of Fig. 6. Not only can a particular co-factor bind and interact with different modules, but also a single module can accommodate one or more co-factors and their analogues, in multiple different poses.

On the contrary, metal ions and metal clusters[48] are precicely define the geometry of binding residues as shown in multiple examples in Fig. 6. This reflects the organometallic interactions driven by the ion coordination sphere, restricting the degree of side chain flexibility. For instance, in several modules consisting of His and Asp/Glu residues (Fig. 6), the ion-interacting functional atoms are fixed, and superpose well even in unrelated enzymes. A general conclusion drawn here is that metal binding motifs are frequently convergent in 3D, while motifs binding organic co-factors are significantly more variable. This is because large co-factors are flexible, therefore can bind in various ways, and some of them might contribute different chemical groups in different catalytic mechanisms.

With respect to the “substrate/product” ligand category, module annotations are not as clear as those in the “metal” and “co-factor” categories. Usually, a cognate ligand analogue will lie close to the catalytic residues, overlapping with residues involved in metal and co-factor binding. This is due to the relatively loose criteria we have set to identify neighbouring ligands (distance cut-off and the requirement of at least one ligand atom to lie close to the module residues) and the fact that usually enzymes are co-crystallised with a cognate ligand analogue that often interacts with co-factors. This highlights the multi-functional nature of several modules, as presented in multiple cases in Table S3. Based on these observations, it can be deduced that “substrate/product” annotation is typically of low confidence (substrate/product analogues of high similarity to the cognate ones are rarely seen in crystal structures[44]). For this reason, complementary approaches are necessary to characterise the modules, as discussed in the following paragraphs.

#### Module enrichment with catalytic residue functional roles

The second functional characterisation approach was the enrichment of each aligned residue position in a module with mechanistic role annotations from M-CSA. In M-CSA each residue is assigned a catalytic function in the mechanism – Broadly these can be classified as one of three roles - catalytic (directly involved in catalytic rearrangements), ligand binding or stabiliser/spectator role. In Fig. 7, we present the functional preferences of modules consisting of three or more convergent templates. For clarity, we decided to include only one instance of each CATH domain in each module, to reduce bias from over-represented folds (e.g. DE-DE-DE-K module, Fig. 3). Pie charts in Fig. 7 show the relative occupancies of role categories (for role classification see Table S1) on each residue position, and the occupancies for the entire module (leftmost pie charts).

**Fig. 7:**
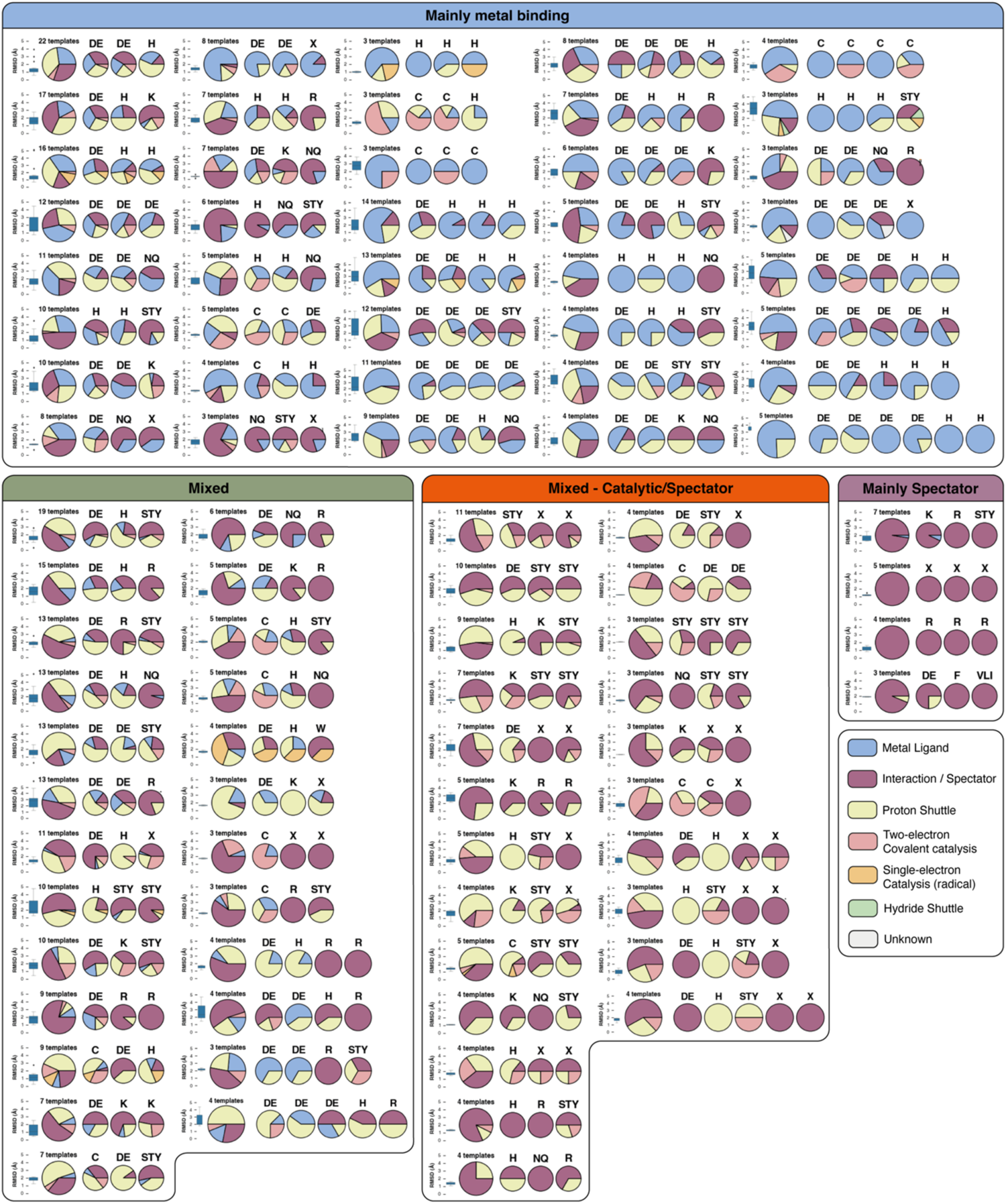
Functional characterisation of catalytic modules. For each module, the occupancies of functional role categories at each residue position are shown as pie charts. Residues at each position are named by the amino-acid one letter code, with multiple letters indicating different, chemically equivalent residues (the letter X is used for residues of unspecific type contributing backbone atoms). The leftmost pie chart shows the sum of occupancies over all residues. Conformational variation among constituent templates is shown as a boxplot of RMSD values over a representative template. To reduce bias from over-represented structural folds, one template from each unique CATH domain is used to collect annotations for this analysis. Modules with <3 residues are not shown in this figure. Categorisation was performed by manual inspection, considering the relative role frequencies.

Through visual inspection of functional occupancies, we defined four coarse categories that classify modules according to function: 1) Mainly metal binding – where at least 3 catalytic residues are annotated as “metal ligand”; 2) Mixed – where a mixture of functions including “metal ligand” are observed; 3) Mixed - Catalytic/Spectator – where a combination of non-radical covalent catalysis and interaction/spectator roles dominates; 4) Mainly Spectator – where Interaction/Spectator occupy ≥85% of roles.

Metal ligand binding modules are straightforward to characterise. This role is rarely ambiguous and most modules in the library are strongly correlated with co-crystallised metal ions in vicinity (see previous paragraph). However, conflation with roles of other categories, as well as the presence of multiple 3D conformers of the same residue composition, often introduce ambiguity. For instance, the DE-DE-H module is so generic that “proton shuttle”, “metal ligand” and “interaction/spectator” roles are observed in almost equal occupancies. In fact, this module serves multiple functions, one of which is Mn^2+^ and Mg^2+^ binding. The next two categories are of mixed function. Characterisation is not as trivial in those cases, and each module needs to be examined individually to elucidate its precise functions. However, a few broad observations can be made. For instance, His-Asp/Glu containing catalytic triads are captured and are, not unexpectedly, highly recurring. Modules corresponding to other triads (e.g. DE-H-R or DE-H-NQ) are also highly represented in this category. This reinforces the hypothesis of functional convergent evolution. Indeed, the acid-base pair (DE-H) is preserved and plays an activating role, while function is determined by the third residue, usually a catalytic nucleophile. We also defined a special category for modules of high interaction/spectator role content. Four modules are in this category (K-R-STY, X-X-X, R-R-R, DE-F-VLI) and all have an auxiliary role, such as charge stabilisation (X-X-X gamma-turn hole), stabilisation of radicals (DE-F-VLI) and interaction with co-factors (R-R-R and K-R-STY). A special comment should be made on the presence of oxyanion holes in the library. Usually those are formed by the backbone of two low-conserved residues. Since our method does not capture two-residue modules, all instances of oxyanion holes are found as two aspecific residue pairs (X-X) and a residue from the adjacent catalytic module. In almost all example, the X-X pair has interaction/spectator roles in high occupancies.

#### Modules as mechanistic components

The third functional characterisation approach based on the concept of the “rules of enzyme catalysis”[50]. Catalytic mechanisms can be broken down in a series of reactions involving catalytic residue functional atoms and substrate/co-factor atoms: the reaction centres of the mechanism. We applied a semi-automated method to identify recurring mechanistic steps in modules, as illustrated in the examples below.

The first example is the DE-DE-DE module. This functionally heterogenous module facilitates metal binding (see Table S3) as well as various catalytic and supporting functions (Fig. 7). We found that this triangular arrangement of acidic residues is a common motif in saccharide-cleaving enzymes. In Fig. 8a, it is shown that this module with this particular functionality is found in at least three functionally convergent and structurally diverse enzymes: Glucan 1,4-alpha-maltohydrolase (EC 3.2.1.133), Glucansucrase (E.C. 2.4.1.125) and Xyloglucan endotransglycosylase (EC 2.4.1.207). The latter two share three very similar rules, directly involving two Asp/Glu residues in the reaction centres, while the third acts as an electrostatic stabiliser. Co-crystallised substrate analogues (glucan disaccharides) bind close to the two catalytic Asp/Glu residues, though at slightly different poses, as seen in 3D in Fig. 8a. In Fig. 8b, the first step of the catalytic mechanism, taken from M-CSA entry 1002 (Glucantransferases), which is shared through the respective rule (rule ID: 799170) is illustrated in 3D (left panel). In this reference mechanism, elucidated by QM/MM simulations[51], the glucosyl group anomeric carbon of the sucrose incurs a nucleophilic attack from Asp477 (covalent catalysis role). Subsequently, the oxygen of the glycosidic bond is protonated by Glu515, causing cleavage, and leaving of the fructosyl group.

**Fig. 8:**
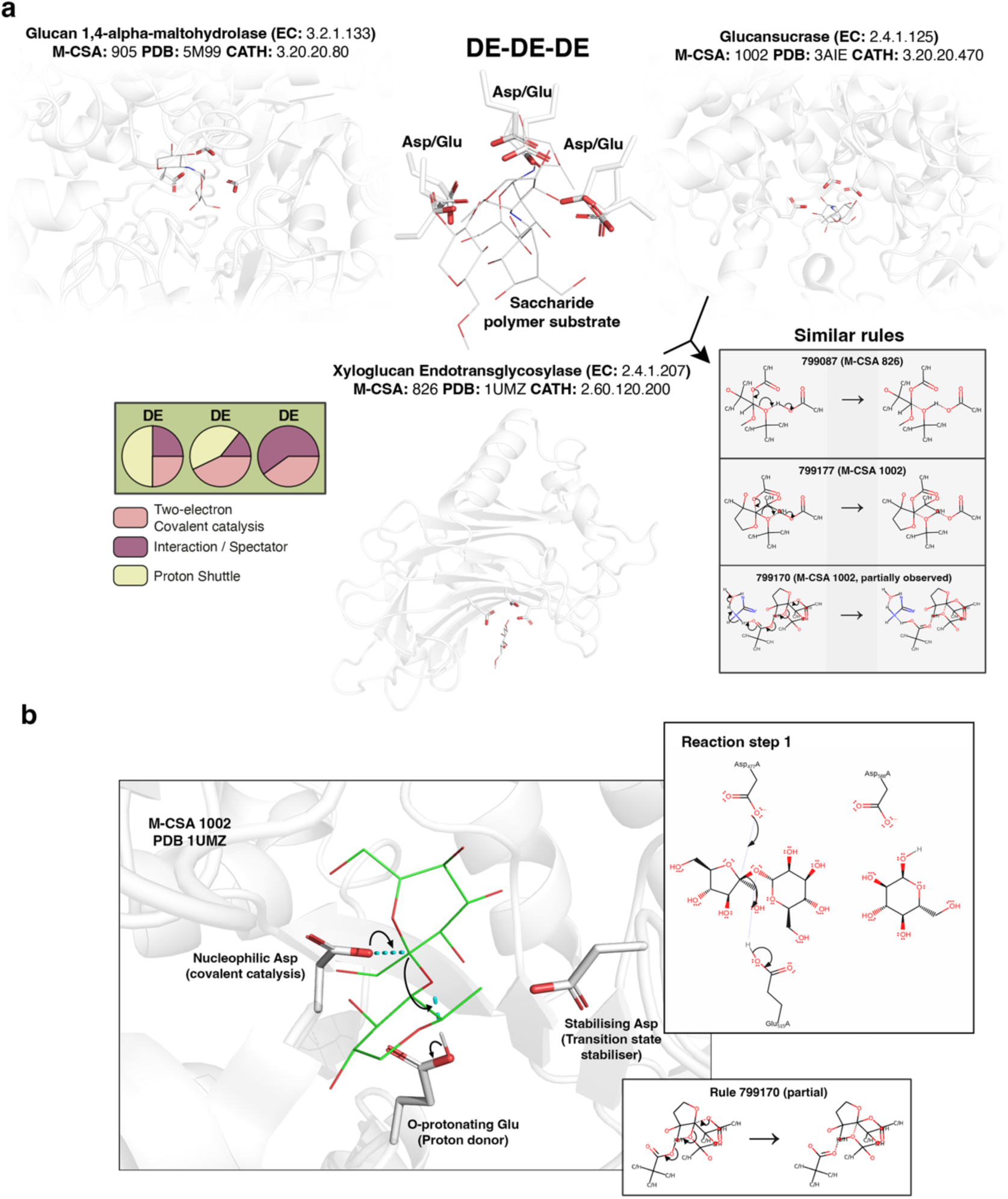
Mechanistic characterisation of the DE-DE-DE module. a: Superposition of three templates of the module over their functional atoms (center). Catalytic residue side chains, and bound ligands are shown as thick and light sticks respectively. Functional atoms are distinguished by increased opacity. Templates along with their structural context are shown individually around the central panel (context is shown in semi-transparent cartoon representation). Functional roles of each residue position are shown in the form of a pie chart in the lower left panel. Shared rules of highly chemical similarity are shown in the lower right panel, in the form of chemical equations. Here, since no identical rules are shared, the name of the M-CSA entry is shown next to each rule ID. b: Example of a catalytic step (initial step of saccharide bond cleavage), shared between two templates of the module. The left panel and upper right panels show the step in 3D and 2D representation, where curly arrows indicate electron transfers towards covalent bonds (dashed sticks/lines). For the 3D representation, the bound ligand has been slightly modified by substituting some atoms in order for its chemical structure to match the cognate ligand of the reaction. The lower right panel shows the respective rule of the step in the form of a chemical equation. Atoms throughout the figure are coloured according to their type (white: C, red: O, blue: N, yellow: S, orange: P). All 3D models were prepared in PyMol[49].

The second example (Fig. 9), is well studied. The Cys-His-Asp catalytic triad is one of the most profound examples of convergent evolution, found in several enzymes of the hydrolase (E.C. 3.-.-.-) and transferase (E.C. 2.-.-.-) classes. In Fig. 9a, we present three enzymes of those classes: Arylamine Acetyltransferase (E.C. 2.3.1.118), Ulp1 Peptidase (E.C. 3.4.22.68) and Glutamyl Hydrolase (E.C. 3.4.19.9). All have unrelated folds, and contain the Cys-His-Asp triad (C-DE-H) in their active sites. These three templates are mapped to several almost identical mechanistic steps, with the templates from Ulp1 Peptidase and Glutamyl Hydrolase sharing four consecutive and identical rules. As an example in Fig. 9b, we show the third step of the reference mechanism (Ulp1 Peptidase, M-CSA entry 820). The respective rule (rule ID: 828513) represents three electron transfers in a series, starting from the *δ* nitrogen atom of the His imidazole ring towards the catalytic Cys which is covalently attached to the peptide bond, and through a water molecule. The Asp/Glu serves consistently as pKa regulator to stabilise the His activator. A comment here regards the template from Glutamyl Hydrolase. As seen in the superimposed models in Fig. 9a, the side-chain conformation of the catalytic Cys is considerably different from the other two examples. However, it is important to note that the position of the endpoint functional sulfur atom is similar across all three analogues, regardless of the orientation of the side chain. This is a good example of how evolution might “invent” the same functional configuration multiple times: what is required, is the catalytically crucial atoms to be in a precise position, regardless of the geometry of the hosting residues and the overall structural context. This can also be reflected in the RMSD of fitted templates, exhibiting increased values in cases like this, although the function is the same.

**Fig. 9:**
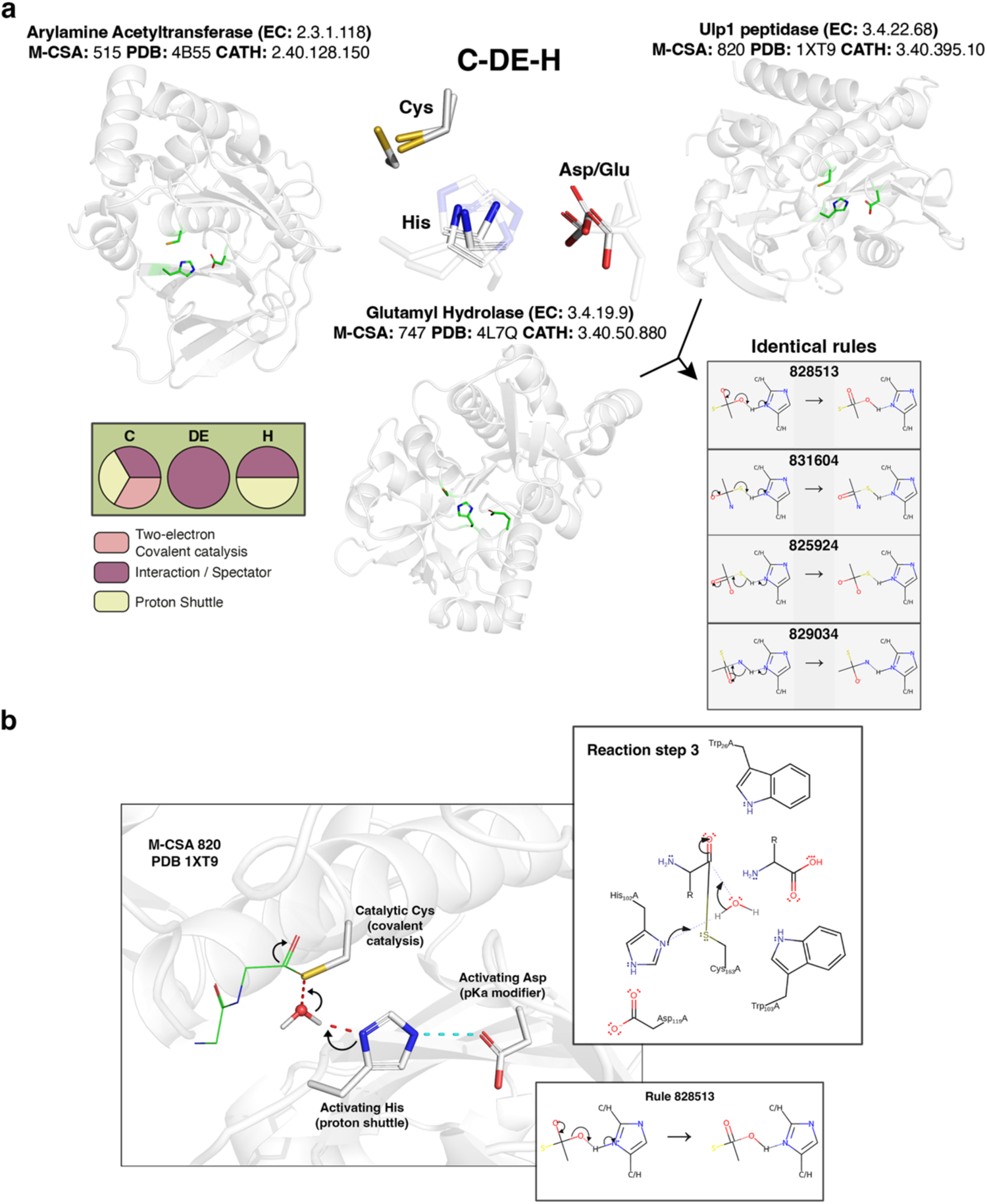
Mechanistic characterisation of the C-DE-H module. a: Superposition of three templates of the module over their functional atoms (center). Catalytic residue side chains are shown as sticks. Functional atoms are distinguished by increased opacity. Templates along with their structural context are shown individually around the central panel (context is shown in semi-transparent cartoon representation). Functional roles of each residue position are shown in the form of a pie chart in the lower left panel. Four identical and consecutive shared rules of are shown in the lower right panel, in the form of chemical equations. b: Example of a catalytic step (third step, involving electron transfers from the His to the covalently attached substrate to the catalytic Cys), shared between two templates of the module. The left panel and upper right panels show the step in 3D and 2D representation, where curly arrows indicate electron transfers. For the 3D representation, the bound ligand has been slightly modified by substituting some atoms in order for its chemical structure to match the cognate ligand of the reaction. The lower right panel shows the respective rule of the step in the form of a chemical equation. Atoms throughout the figure are coloured according to their type (white: C, red: O, blue: N, yellow: S, orange: P). All 3D models were prepared in PyMol[49].

## Discussion

Collecting and archiving information on the elements at the heart of biological catalysis has allowed the definition and identification of currently known “3D modules of enzyme catalysis”. By definition, a catalytic module is found in several enzymes, which are diverse in function and/or structure. Through a novel methodology involving 3D templates, we identified and catalogued 161 modules of varying abundance, and conformational/functional diversity.

The ultimate challenge of this analysis, was functional characterisation. A fully characterised and well-defined module would require each residue position to correspond to a precise functional role and/or a mechanistic step. Ideally, since catalytic modules participate in a series of reactions during the course of the catalytic transformation (steps), similarities in the detailed mode of action of each enzyme needs to be investigated .We show that a large number of catalytic modules facilitate binding of metal ions that are naturally not involved directly in electron/proton exchanges, making their characterisation relatively straightforward (annotation by ligand type or by role). Characterisation in mixed-role and non-metal binding modules however, as discussed, is more complex. Residue functional annotation, primarily for stabilising residues relies in manual curation and the available information of the literature, thus inconsistencies might be present. As can be summarized from Fig. 7, the striking majority of modules (apart from metal binders) are of mixed function, complicating automated characterisation. However, this also reflects the complexity of biological catalysis, the multitude of residue roles and also conformational variability. Another issue regards the sensitivity of template-template matching: Side chain conformations are rather variable and even in divergent enzymes[17] they often change their orientations and would not be picked up by template searching.

Looking at bound ligands in template neighbourhood provided good insight on module functionality. We characterised each module according to its residue-ligand interaction propensity and showed that ∼1/3 of modules in the library facilitate metal binding with high propensity and in several cases, ligands of more than one type can interact with a module. Several examples of convergent modules in Fig. 6 show that coordination by metals is a very important factor applying evolutionary pressure to the conformation of binding residues, resulting in finding instances of such modules in multiple, unlinked by evolution, enzymes. Functional atoms in these modules differ only slightly in geometry, something reflected in low RMSD values, despite the fact that residues contributing them might lie in completely different positions in the sequence and may not even share the same sequence order. Conversely, we found surprising that convergent patterns facilitating interaction with organic co-factors were not identified in this analysis. The explanation here is that large, flexible co-factors assisting multiple enzymatic functions (especially in pleiotropic enzymes[52]), might bind in several different ways and sometimes contribute different atoms according to what the mechanistic procedure requires. From our observations, two types of structural variation are distinguished: a flexible co-factor adopts several poses and can interact with different residue arrangements (for instance, several modules are found to interact with NAD, according to Table S3), and a residue arrangement can bind more than one co-factors. Finally, substrate binding is observed to be the least straightforward annotation, since it is highly likely that a substrate, especially a bulky one, will be located close to all catalytic residues and can co-exist with metals and co-factors.

Overall, our analysis shows that structurally similar modules can be involved in more than one enzymatic function, often being radically different in function themselves, and may have emerged more than once during evolution. A good example is the DE-DE-DE module, which is found to bind metals as well as being involved directly in catalysis, even though the residue geometry is relatively similar in both functions. Therefore, it is crucial to assess all three approaches of characterisation in order to assign the function of a module, taking also into account the structural diversity within. A limitation of this work is that this library is populated only by data stored in M-CSA. This automatically biases the analyses towards well studied enzymes that include detailed mechanistic information. However, as shown in Fig. 2a, the database covers a large functional landscape in terms of EC classes, therefore the amount and diversity of active site annotations in the current version provides a good start in examining the abundance of modules. There are two ways of expanding this coverage: 1) Use catalytic residue annotations from other resources like UniProt[53]. This though, comes with a compromise in reliability, since the advantage of M-CSA is the human-hand annotation of functionally important residues, compared to the automated ways used in general purpose resources. 2) Expansion of structural coverage and identification of further convergent evolution examples by scanning experimental and/or predicted[54–56] protein structures with the templates forming the modules. We are currently developing a protocol to perform exactly this, with the biggest challenge lying in distinguishing true, functional template matches from spurious ones, especially when functional information on the target proteins is limited (e.g. in uncharacterized predicted models from trEMBL[53]).

Beyond the academic interest that led us to investigate the modularity of active sites, these modules can be used as tools in computational enzymology. A library of functionally characterised, recurring small motifs can be used to identify functional sites in experimental or predicted protein structures. For each module, one or more templates can serve as probes to identify instances of the module in a protein structure[12,57–59], using template matching software like *Jess*[22]. Template matches can then benefit from the annotations provided in this study (see Table S3). This can be particularly useful, especially when investigating evolutionary relationships between enzymes and events like residue substitution and conservation of particular segments of the active site. Another application is the development of enzymes of novel function. We showed that active sites can be partially conserved in 3D, especially within residue groups coordinated by a ligand or serving a very precise catalytic function. This knowledge, in the form of the module library, can be used when developing a novel enzyme, incorporating *de novo* and directed evolution techniques[60,61]. A relevant concept, the concept of “theozymes”[62], can also use modules as templates for theoretical moulding of the active site, to perform a novel catalytic mechanism.

Although all data used here are available upon request, we are planning to release the template library in a detailed and entry-based structure within M-CSA in the close future. The immediate plan is to map residue-level mechanistic annotation in all modules. This will make modules an even more robust tool, to provide mechanistic insight in functional annotation and investigation of evolutionary relationships.

## Methods

### Template library methods

#### Catalytic site library

A library of enzyme catalytic site structures was generated by integrating data primarily from the Mechanism and Catalytic Site Atlas (M-CSA)[1] and the Protein Data Bank (PDBe)[63]. A detailed description of the method for building a catalytic site library is provided in the methods section of our recent work on catalytic site 3D variation[17]. In brief, curated catalytic residue annotations for 1003 enzyme families were collected from M-CSA, 734 of which included a detailed description of the catalytic mechanism in a computer readable format[50]. These annotations were used to extract atomic coordinates of catalytic residues from experimentally resolved protein structures (biological assemblies) and a clustering algorithm was applied to reconstruct all catalytic sites. The use of a clustering algorithm is important in this step: catalytic residues in identified homologues are annotated at the sequence level, and in cases of active sites consisting of residues from multiple identical chains, all residues sharing the same identifier will be annotated on the structure. Therefore, geometrical heuristics needed to be applied to identify the correct combination of catalytic residues forming an active site. Further information was added to the catalytic site library, such as co-factor and catalytic reaction annotations, cognate/bound ligand similarity scores[44] and mapping to protein sequences from SwissProt[53]. Lastly, active sites extracted from an M-CSA family, were overlayed onto their respective reference active site. Each ensemble of superimposed active sites effectively represents the different conformers captured in experimentally resolved structures.

#### Extraction of “complete” templates

In our recent review[12], we showed that each alternative conformational state of an active site can be represented by a consensus template –an abstract active site signature (“complete” template) containing only a subset of three functional atoms per catalytic residue. The selection of the atoms depends on the chemical nature of the residue, as well as on its function location (backbone or side chain atoms) within the context of the catalytic reaction, as annotated in M-CSA. Since templates are primarily used to probe protein structures for similar atom arrangements through a template matching algorithm[19,20,22,64,65], it is important to account for matching fuzziness in the residue and atom level. For instance, chemically equivalent amino acids should be able to be matched interchangeably (Asp ↔ Glu, Asn ↔ Gln, Ser ↔ Thr ↔ Tyr, Val ↔ Leu ↔ Ile), as well as carbonyl Ο atoms in Asp/Glu residues or N_δ1_ and C_δ2_ atoms in His residues, to account for side chain flipping and ambiguity in crystal structures[66].

Fig. S2 schematically describes the process for deriving “complete” templates for 813 M-CSA enzyme families. The steps are the following: Active sites from M-CSA families with ≥3 annotated catalytic residues were first constructed. Sites within a family, were then superimposed over their functional atoms in an all-vs-all fashion. For the purposes of this study, only active sites having all catalytic residues conserved (compared to the curated reference catalytic residues) or only containing chemically equivalent mutations were included. If aligned residues are known to function via their main chain or known to be post-translationally modified, any mutation is allowed. Rigid body superposition was RMSD-based and performed in an iterative manner (by updating the average structure at each cycle) using a modified, gaussian-weighted *Kabsch* algorithm[67,68] as introduced by Damm et al.[68]. Three functional atoms per catalytic residue are used for superposition with the exact definitions shown in Fig. S1. The next step was the calculation of an average active site structure and the selection of a representative template active site having the closest similarity to the average after superposition. After excluding those structures for which a template could not be generated due to insufficient electron density (some residues might be missing from the structure), we ended up with 813 “complete” templates. Those are formatted using a modified PDB nomenclature[64] and are compatible with the template matching software *Jess*[22,64].

#### Fragmentation of complete templates into “partial” templates

Complete templates were further fragmented into smaller, “partial” templates of 3,4,5,6,7 and 8 residues, lying in close proximity. The idea behind this is that residue groups contributing to a particular function, will be typically expected to form tight clusters. Thus, for a complete template consisting of *S* residues, we need to identify those partial templates which optimally cluster *S* residues into *k* partial templates. The fragmentation algorithm is purely geometrical and includes the following steps (Fig. 1): 1) Calculation of the centre of mass of the three functional atoms of each catalytic residue, 2) calculation of the number of partial templates (residue clusters) *k* ≈ *S*/*s*, where *S* is the size of the complete template (number of residues) and *s* is the desired size of the partial template. It should be noted that potential partial templates of all possible sizes were generated for each complete template. 3) Application of k-means algorithm in 3D cartesian space to define *k* clusters of catalytic residues’ centres of mass and 4) calculation of the centre of each cluster. 5) Finally, the first *s* catalytic residues closest to each centre are selected to define the partial template. Applying this method to all 813 complete templates led to a total of 3953 partial templates, with sizes spanning from 3 to 8 residues. Statistics on the template library (sizes, E.C. and fold distributions) are shown in Fig. 1c and are discussed further in the Results section.

### Module library methods

#### Template-template search

Templates of the same size were cross-compared to capture groups of similar templates. This template-template search was performed in an all-vs-all fashion using the software *Jess*[22]. The standard functionality of Jess involves a template and a query structure; in our case the query structures are replaced by templates themselves, permitting the identification of template pairs that share structural similarity up to a defined atom displacement (functional atoms’ RMSD). Template matching is fuzzy both at the residue and atom level as mentioned in the previous paragraph, allowing matching of chemically similar residues and atoms (Fig S1). Here again, for residues known to function via their main chain or those which are post-translationally modified, three backbone atoms of any residue can be matched. *Jess* requires two user-defined parameters to perform the search: a maximum template-query pairwise atom distance (set to 2.5Å) and a maximum RMSD after optimal superposition of the template to the query (set to 2.0Å). These cutoffs were set empirically, after performing multiple template-template search runs to determine a good balance between true positive and spurious matches. Using these parameters, *Jess* will try to identify a set of atoms satisfying the constraints coded within the template (atom and residue types) allowing a maximum variation equal to the atom distance cutoff, and then perform rigid body superposition, outputting matches with RMSD lower or equal than the RMSD cutoff. The final output of these template-template matching runs consisted of 15207 template pairs (*p-q* pairs). These hits were reduced to 1481 after removing redundant pairs (*p-q*/*q-p* permutations), and instances where at least one template in a pair is a subset of a larger one existing in the set. Furthermore, hits where both *p* and *q* derive from the same PDB structure were removed.

#### Graph representation and clustering

The 1481 template relationships constitute a network that can be drawn in the form of a graph (Fig. 3), consisting of a set of smaller subgraphs, within which all templates share the same residue composition (accounting for residue type fuzziness as explained previously). Subgraphs of the same residue composition define a set of “modules”. Structural clusters within each module were identified by graph clustering, using the Leiden algorithm and RMSD as similarity metric. Finally, a total number of 708 templates were assigned to 123 modules and 213 structural clusters.

#### Alignment of residues within modules and clusters

To facilitate precise residue level annotation of the templates within each module (e.g. to derive functional role statistics in each residue position), it was important for all residues to be aligned consistently. The approach is as follows: a reference template is found within each structural cluster of a module; in the corresponding subgraph, this is the node with the highest centrality (defined as the number of edges connected to a node). The residue order of the reference template is used to find the correct residue order of as many templates within the cluster as possible, from the raw output of the template-template search (before cleaning-up). Often, subgraphs are not fully connected (not all templates match all templates under the defined matching parameters), therefore for templates for which an alignment could not be found in the similarities set, this was found by freshly performing pairwise structural alignment to the reference, through applying the same gaussian-weighted superposition algorithm described further above in active site superposition. Last, to achieve residue alignment within the module, aligned residues within the clusters were reordered by finding the correct residue correspondence of each cluster reference to the reference of the largest cluster. A point of care here is the following: since structural similarity of functional atoms of chemically similar residues is the only criterion to find the ideal aligned positions, ambiguity is often unavoidable, especially in templates of convergent geometry containing more than one instance of a residue type (or equivalent residues such as D/E, S/T/Y or V/L/I).

#### Module annotation

After assignment to a module and structural cluster, and subsequent residue reordering, each template was annotated with structural and functional information. Basic information includes template size, mean inter-residue distance and RMSD to the corresponding reference template. The name of a module corresponds to its fuzzy residue composition (e.g. DE-DE-NQ-X), where each residue is written in one-letter code. Alternative residues in a position are written as multiple letters, with X and Z used for unspecific residue types contributing backbone atoms and post-translationally modified residues, respectively. Further annotations were collected from the M-CSA public API, such as EC number[69], CATH superfamily[15] and functional roles[70,71] for each residue position. The latter were grouped into empirical categories (metal ligand, proton shuttle, hydride shuttle, two-electron covalent catalysis, single-electron catalysis (radical), interaction/spectator) with the exact definitions[70,71] presented in Table S1. It should be noted that catalytic residue roles annotations rely on the underlying literature and discretion of the curator when describing the mechanism. Roles, especially those belonging to the Spectator/Interaction class, are often considered auxiliary and are subject to ambiguity. Also, residues might perform different roles in different steps during the course of the catalytic mechanism. For these reasons, “Metal Ligand” and “Reactant” classes are prioritised over “Spectator/Interaction” ones, when they co-exist at a residue, since the former are annotated more accurately using experimental/computational evidence.

In order to identify ligands interacting with module residues, we utilised the CSA-3D framework of our previous work[17], with the exact methodology being described thoroughly therein. For each template in a module, all ligands located 6Å or closer to all catalytic residue (minimum interatomic distance between ligand and residue in 3D space) are extracted and classified in four categories: Metal ions, co-factors (organic or inorganic co-factors composed of more than one atom, such as NAD and Fe-S metal clusters respectively), substrates (small molecule or polymer) and crystallographic artefacts (such as glycerol or ions irrelevant to the catalysed reaction). Classification involves an initial step of ion, co-factor and artefact identification by looking up their PDB three-letter codes in respective curated databases[45,47]. Any ligands not falling within these categories are classified as putative substrates/products. Subsequently, each ligand is compared as per its chemical similarity (PARITY score[44]) with all cognate ligands of the reaction that the respective enzyme is known to catalyse (reaction information is collected from KEGG[72,73], Rhea[73] or M-CSA). For non-polymer substrates/products, in order for a ligand to be classified as a substrate, it needs to resemble one of the cognate substrates/products at ≥70% chemical similarity. Crystallographic artefacts are ignored since they are usually biologically irrelevant. Since a template represents a cluster of similar, homologous and conserved active sites, ligand information is collected from all those sites. Frequency of occurrence for each ligand in each template is calculated, and frequencies are aggregated over all templates of a module. Additionally, a score is calculated for each of the three ligand categories in each template: if a site within a template contains at least one ligand of a given category, the corresponding score in the template is incremented by 1. An average score is then calculated at the module level.

Last, we added mechanistic information associated with the catalytic site from which each template derives; this information is coded in the form of “the rules of enzyme catalysis”, a concept introduced in our recent work[50]. Each rule represents a complete or partial step of the catalytic mechanism and is coded in a SMARTS string, representing a reaction involving the reaction centres of the catalytic step plus all atoms within two bonds of the reaction centre atoms. We compiled a database of these rules as part of M-CSA, that can be used to search an active site in 3D for possible mechanisms. We have also shown that rules can be a tool to investigate evolutionary relationships between enzymes at the mechanism level[14,74], by pairwise and stepwise rule comparison[75]. This is the basis of the characterisation method here: all rules mapped to respective M-CSA entries of the templates within a module are compared. However, as of yet, M-CSA rules are mapped on the active site level and not on the residue level, complicating the automation of the module annotation. Therefore, each case needs to be examined individually to check whether the rules shared between templates actually involve catalytic residues from these particular templates. A systematic analysis with precise residue-level rule mapping is currently in progress, and we plan to release a detailed, fully annotated library, where modules can be used as tools themselves for identifying enzyme mechanistic relationships in the context of convergent and divergent evolution. In the current analysis we used the version 1.0 rule set; in order to derive a set of generic rules able to capture mechanistic steps that are similar but not identical, rules were clustered according to three similarity features: reactants and products chemical similarity, bond changes and reaction centres similarity. The average of these scores was used as a similarity metric to perform hierarchical clustering and group together rules being ≥70% similar. If two rules share a chemical similarity of ≥70%[76], templates are considered to share a common mechanistic step, adding further evidence of functional convergence.

## Data availability

All data is available upon request.

## Author contributions

IGR: Conceptualisation, Software, Methodology, Validation, Formal analysis, Investigation, Visualisation, Writing – Original Draft preparation, Writing – Review & Editing

AJMR: Conceptualisation, Data Curation, Writing – Review & Editing NB: Conceptualisation, Writing – Review & Editing

JMT: Conceptualisation, Supervision, Resources, Funding Acquisition, Project Administration, Writing – Review & Editing

## Acknowledgements

The work was supported by: the EMBL International PhD Programme (IGR) and the European Molecular Biology Laboratory (AJMR, NB, JMT)

