## Supplementary Information for "The 3D modules of enzyme catalysis: deconstructing active sites into distinct functional entities"

#### Supporting Information

##### Figures

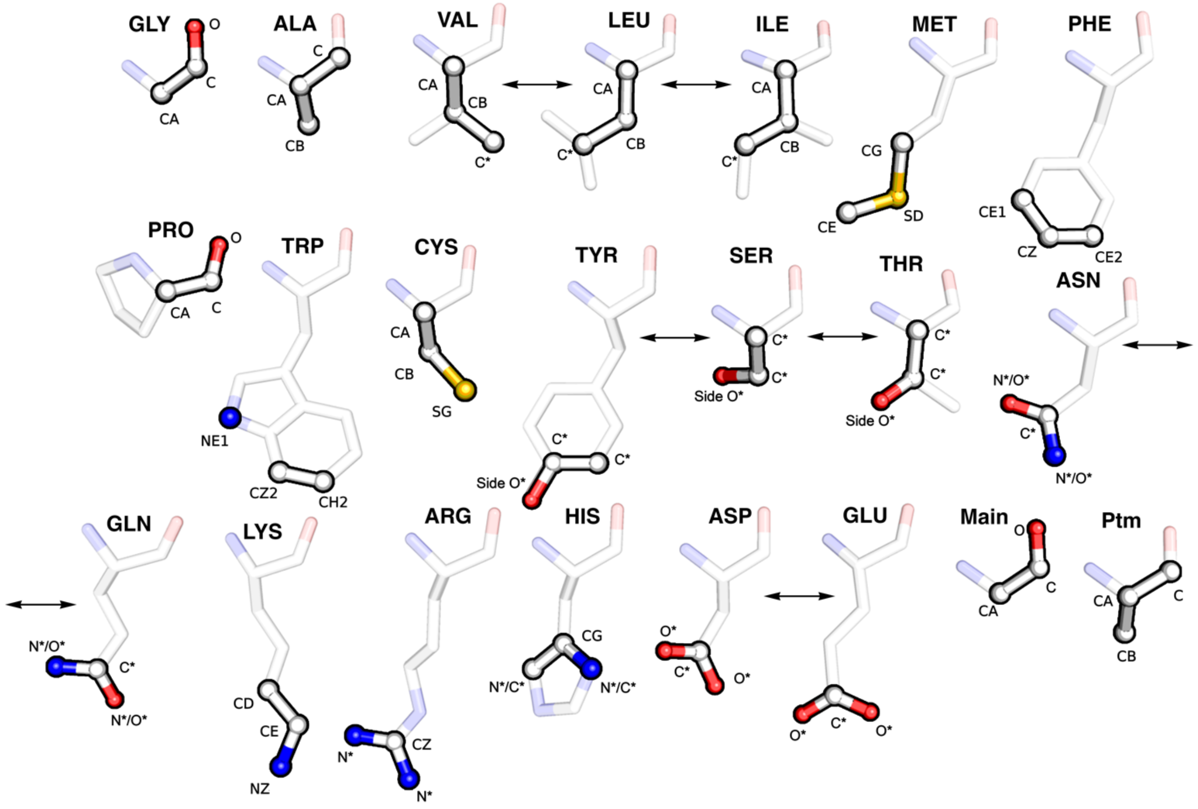

*Fig. S1:* ***Functional atoms used in superposition of homologous active sites****. For each residue type, three residue-specific atoms are selected (shown in opaque sticks) if the residue functions via its side chain. In cases where a residue functions via its main chain or undergoes post-translational modifications, explicit atom selections are defined, as shown in the lower right part of the figure. Bidirectional arrows indicate residues of equivalent properties that can be superposed interchangeably. Similarly, atoms of symmetrical chemical groups or atoms that are shared between equivalent residues are indicated with a * symbol. These atom selections are a modified version of the definitions from Wallace et al[1]. 3D models were generated in PyMol[2].*

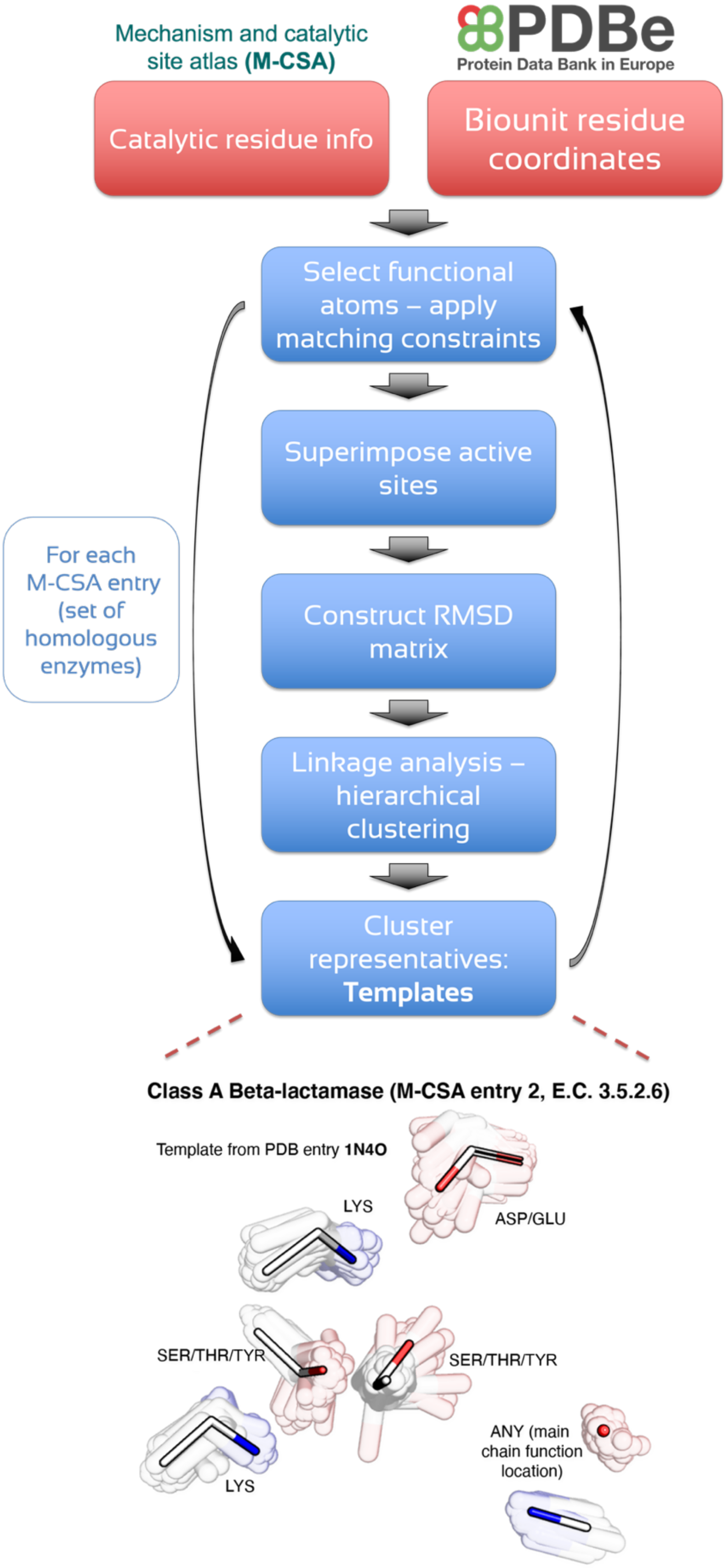

*Fig. S2:* ***"Complete" template extraction process.*** *The upper part of the figure is a flowchart of the method used to derive "complete" templates for an M-CSA family. Abstracted active sites from homologous enzymes are superimposed and clustered. A "complete" template (bottom part of figure) is the representative structure of each cluster. Functional atoms of active sites from homologous active sites of the same structural cluster are shown as transparent sticks in superposition. Template atoms are shown as opaque sticks. In this simple example, only a single cluster exists. Note that residues like Asp-Glu and Ser-Thr-Tyr are considered equivalent. 3D model was generated in PyMol*[2]*.*

### Tables

*Table S1: Catalytic residue functional roles and their classification*

| Role | Category | Class |
|---|---|---|
| Nucleofuge | Two-electron Covalent Catalysis | Reactant |
| Nucleophile | | |
| Electrophile | | |
| Electrofuge | | |
| Covalent Catalysis | | |
| Electron pair acceptor | | |
| Electron pair donor | | |
| Electron shuttle | | |
| Covalently attached | | |
| Leaving group (radical) | Single-electron Catalysis (radical) | |
| Radical combinant | | |
| Single electron acceptor | | |
| Single electron donor | | |
| Single electron relay | | |
| Electron tunnelling medium | | |
| Single electron shuttle | | |
| Hydrogen radical relay | | |
| Proton donor | Proton Shuttle | |
| Proton acceptor | | |
| Proton shuttle (general acid/base) | | |
| Proton relay | | |
| Hydride acceptor | Hydride Shuttle | |
| Hydrogen radical donor | | |
| Hydrogen radical acceptor | | |
| Metal ligand | Metal ligand | Metal ligand |
| Polar interaction | Interaction | Interaction/Spectator |
| Hydrogen bond acceptor | | |
| Hydrogen bond donor | | |
| Pi-pi interaction | | |
| Polar/non-polar interaction | | |
| Attractive charge-charge interaction | | |
| Electrostatic stabiliser | Spectator | |
| Activator | | |
| Increase basicity | | |
| Increase acidity | | |
| Promote heterolysis | | |
| Modifies pKa | | |
| Increase nucleophilicity | | |
| Steric role | | |
| Electrostatic interaction | | |
| Radical stabiliser | | |
| Promote homolysis | | |
| Alter redox potential | | |
| Enhance reactivity | | |
| Transition state stabiliser | | |
| Increase electrophilicity | | |
| Electrostatic destabiliser | | |
| Increase redox potential | | |
| Steric locator | | |
| Steric hindrance | | |
| Ground state destabiliser | | |
| unknown | Unknown | Unknown |

*Table S2: Divergent subset of templates in modules, grouped by CATH superfamily*

| Module | CATH domain | Enzymes (EC) | Mean RMSD (Å) |
|---|---|---|---|
| C-C-C | 3.20.20.70 | 7-carboxy-7-deazaguanine synthase (4.3.99.3)<br>biotin synthase (2.8.1.6)<br>RlmN (2.1.1.192) | 1.01 |
| C-NQ-X | 3.60.20.10 | amidophosphoribosyltransferase (2.4.2.14)<br>ferredoxin-glutamate synthase (1.4.7.1) | 1.37 |
| C-NQ-X-X-X | 3.60.20.10 | hexosephosphate aminotransferase (2.6.1.16)<br>GOGAT (1.4.1.13) | 1.58 |
| DE-DE-DE | 3.20.20.120 | enolase (4.2.1.11)<br>chloromuconate cycloisomerase (5.5.1.7) | 0.94 |
| | 3.20.20.80 | neopullulanase (3.2.1.135)<br>4-alpha-glucanotransferase (2.4.1.25) | 1.12 |
| DE-DE-DE-DE-H | 3.20.20.120 | L-fuconate dehydratase (4.2.1.68)<br>mandelate racemase (5.1.2.2)<br>muconate cycloisomerase (5.5.1.1) | 0.28 |
| DE-DE-DE-DE-H-H | 3.40.630.10 | bacterial leucyl aminopeptidase (3.4.11.10)<br>glutamate carboxypeptidase (3.4.17.11) | 0.66 |
| DE-DE-DE-DE-K | 3.20.20.120 | muconate cycloisomerase (5.5.1.1)<br>OSBS (4.2.1.113) | 2.35 |
| DE-DE-DE-H | 3.20.20.120 | L-fuconate dehydratase (4.2.1.68)<br>mandelate racemase (5.1.2.2)<br>muconate cycloisomerase (5.5.1.1) | 1.60 |
| | 3.30.420.10 | duplicase (2.7.7.7)<br>Unknown (3.1.-.-) | 1.71 |
| DE-DE-DE-H-H-H-STY | 3.40.720.10 | phosphomonoesterase (3.1.3.1)<br>phosphoglycerate mutase (5.4.2.12) | 0.45 |
| DE-DE-DE-H-K | 3.20.20.190 | Unknown (4.6.1.-)<br>glycerophosphodiester phosphodiesterase (3.1.4.46) | 0.77 |
| DE-DE-DE-K | 3.20.20.120 | enolase (4.2.1.11)<br>Unknown (5.5.1.-)<br>mandelate racemase (5.1.2.2) | 1.81 |
| DE-DE-DE-K-K | 3.20.20.120 | Hyp-B 2-epimerase (5.1.1.22)<br>Unknown (5.5.1.-) | 0.80 |
| DE-DE-H | 3.20.20.70 | aldolase (4.1.2.13)<br>PdxJ (2.6.99.2) | 1.63 |
| | 3.40.50.1820 | haloalkane dehalogenase (3.8.1.5)<br>Transferred to 3.3.2.9 and 3.3.2.10 (3.3.2.3) | 1.11 |
| DE-DE-H-NQ | 3.20.20.190 | 1-phosphatidylinositol-4 (3.1.4.11)<br>Unknown (4.6.1.-) | 1.64 |
| | 3.60.21.10 | phosphomonoesterase (3.1.3.2)<br>UDP-sugar diphosphatase (3.6.1.45) | 2.21 |
| DE-DE-H-STY | 3.40.720.10 | phosphomonoesterase (3.1.3.1)<br>phosphoglycerate mutase (5.4.2.12) | 1.81 |
| DE-DE-K | 3.20.20.70 | QAPRTase (2.4.2.19)<br>lysine 2 (5.4.3.2)<br>OMP-DC (4.1.1.23) | 1.36 |
| DE-DE-R | 2.160.20.10 | pectinesterase (3.1.1.11)<br>alpha-1 (4.2.2.2) | 1.76 |
| DE-DE-STY | 2.120.10.10 | anhydrosialidase (4.2.2.15)<br>exo-alpha-sialidase (3.2.1.18) | 0.21 |
| DE-DE-X | 3.40.50.1000 | phosphonatase (3.11.1.1)<br>phosphoserine phosphatase (3.1.3.3) | 1.21 |
| DE-F-K-STY | 3.40.640.10 | kynureninase (3.7.1.3)<br>L-lysine 6-transaminase (2.6.1.36) | 1.61 |
| DE-H-H | 2.60.120.330 | isopenicillin-N synthase (1.21.3.1)<br>leucocyanidin (1.14.20.4) | 1.13 |
| | 3.20.20.70 | acetyl-CoA carboxylase (6.4.1.2)<br>HOA (4.1.3.39) | 1.11 |
| DE-H-H-H | 2.60.120.10 | oxalate oxidase (1.2.3.4)<br>homogentisicase (1.13.11.5) | 1.40 |
| | 3.20.20.140 | urease (3.5.1.5)<br>aminodipeptidase (3.4.13.19)<br>adenosine deaminase (3.5.4.4) | 1.79 |
| DE-H-H-H-H-Z | 3.20.20.140 | urease (3.5.1.5)<br>Unknown (3.4.19.-) | 0.62 |
| DE-H-H-R | 3.40.50.1240 | phosphomonoesterase (3.1.3.2)<br>bisphosphoglyceromutase (5.4.2.4) | 1.75 |
| DE-H-K | 3.20.20.70 | UlaD (4.1.1.85)<br>DHQase (4.2.1.10) | |
| DE-H-NQ-STY-X | 3.40.50.1110 | alkylacetyl-GPC:acetylhydrolase (3.1.1.47)<br>palmitoyl-CoA hydrolase (3.1.2.2) | 0.69 |
| DE-H-STY | 3.20.20.70 | 4-enoyl-CoA reductase (1.3.1.34)<br>trimethylamine dehydrogenase (1.5.8.2)<br>HOA (4.1.3.39)<br>4HmO (1.1.3.46)<br>ribulose-phosphate 3-epimerase (5.1.3.1) | 1.24 |
| | 3.40.50.1820 | acetylxylan esterase (3.1.1.72)<br>Unknown (2.3.1.-)<br>endoprolylpeptidase (3.4.21.26) | 1.28 |
| DE-H-STY-STY-X | 3.40.50.1820 | Xaa-Pro-dipeptidyl-aminopeptidase (3.4.14.5)<br>CocE (3.1.1.84) | 1.43 |
| DE-H-STY-X | 3.40.50.1820 | acetylcholinesterase (3.1.1.7)<br>narbonolide synthase (2.3.1.240) | 0.50 |
| DE-H-STY-X-X | 3.40.50.1820 | butyrinase (3.1.1.3)<br>prolyl aminopeptidase (3.4.11.5)<br>palmitoyl[protein] hydrolase (3.1.2.22) | 1.28 |
| DE-H-X-X | 2.40.10.10 | trypsin (3.4.21.4)<br>nucleoside-triphosphatase (3.6.1.15) | 1.06 |
| DE-H-X-X-X | 3.10.129.10 | trans-2-decenoyl-ACP isomerase (5.3.3.14)<br>beta-hydroxyoctanoyl-ACP-dehydrase (4.2.1.59) | 0.54 |
| DE-K-K | 3.20.20.70 | aldolase (4.1.2.13)<br>indole-3-glycerol-phosphate synthase (4.1.1.48) | 1.04 |
| | 3.40.1160.10 | acetylglutamate kinase (2.7.2.8)<br>isopentenyl phosphate kinase (2.7.4.26) | 0.64 |
| DE-K-STY | 3.40.50.300 | dethiobiotin synthase (6.3.3.3)<br>AvDH1 (3.6.4.12) | 1.74 |
| | 3.40.640.10 | glycine hydroxymethyltransferase (2.1.2.1)<br>glutamate-1-semialdehyde 2 (5.4.3.8)<br>7,8-diaminonanoate transaminase (2.6.1.62) | 1.57 |
| DE-NQ-X | 3.40.50.970 | indolepyruvate decarboxylase (4.1.1.74)<br>CEAS (2.5.1.66) | 1.27 |

| | | | |
|---|---|---|---|
| **DE-R-STY** | 3.40.50.720 | PTR1 (1.5.1.33)<br>UGM (5.4.99.9) | 1.65 |
| **F-NQ-VLI** | 3.40.50.970 | pyruvate oxidase (1.2.3.3)<br>pyruvate:ubiquinone-8-oxidoreductase (1.2.5.1) | 0.86 |
| **H-H-K** | 3.10.130.10 | RNase (4.6.1.18)<br>Unknown (3.1.27.-) | 0.72 |
| **H-H-STY** | 3.40.50.620 | tyrosine---tRNA ligase (6.1.1.1)<br>ATP-sulfurylase (2.7.7.4) | 1.44 |
| **H-H-Z** | 3.20.20.140 | urease (3.5.1.5)<br>aryldialkylphosphatase (3.1.8.1) | 1.37 |
| **H-K-STY** | 3.40.50.720 | 6-phosphogluconic dehydrogenase (1.1.1.44)<br>UDP-sulfoquinovose synthase (3.13.1.1) | 1.38 |
| **K-STY-STY** | 3.40.50.720 | galactowaldenase (5.1.3.2)<br>dTDP-glucose 4 (4.2.1.46)<br>Unknown (1.1.1.30) | 1.46 |
| **K-STY-X** | 3.40.710.10 | beta-lactamase (3.5.2.6)<br>Unknown (3.1.1.-) | 0.55 |

*Table S3: The modules of catalysis and their functional characterisation*

| Module | Number of 3D clusters | Number of templates | Mean RMSD | Enzymes (Name (EC) (CATH)) | Ligand scores (%) | | | Role class propensities (%) | | | Observed ligands | | |
|---|---|---|---|---|---|---|---|---|---|---|---|---|---|
| | | | | | Metals | Co-factors | Substrates | Metal Ligand | Reactant | Spectator | Metals | Co-factors | Substrates |
| **A-X-X** | 1 | 2 | 1.9 | 2-hydroxy-6-oxohepta-2 (3.7.1.9) (3.40.50.1820)<br>omptin (3.4.23.49) (2.40.128.90) | 0 | 0 | 25 | 0 | 100 | 0 | | | HPK (13.04%)<br>MLI (8.7%)<br>C1E (8.7%)<br>BUA (4.35%)<br>PPI (4.35%)<br>KEM (4.35%)<br>PCS (4.35%)<br>C0E (4.35%)<br>BEZ (4.35%)<br>IVA (4.35%)<br>Other (39.15%) |
| **C-C-C** | 1 | 6 | 1.1 | Gfa (4.4.1.22) (3.90.1590.10)<br>NADPH-sulfite reductase (1.8.1.2) (3.30.413.10)<br>7-carboxy-7-deazaguanine synthase (4.3.99.3) (3.20.20.70)<br>biotin synthase (2.8.1.6) (3.20.20.70)<br>RlmN (2.1.1.192) (3.20.20.70)<br>biotin synthase (2.8.1.6) (3.20.20.70) | 5.5 | 91 | 36.8 | 100 | 0 | 0 | ZN (2.13%) | SF4 (53.19%)<br>SAM (19.15%)<br>GSH (4.26%)<br>FES (4.26%) | SF4 (17.02%) |
| **C-C-C-C** | 2 | 4 | 1.5 | Unknown (2.1.1.n11) (3.40.10.10)<br>ADH3 (1.1.1.284) (3.90.180.10)<br>NADPH-sulfite reductase (1.8.1.2) (3.30.413.10)<br>ferredoxin:thioredoxin reductase (1.8.7.2) (3.90.460.10) | 50 | 19.8 | 30.2 | 100 | 0 | 0 | ZN (47.92%) | SF4 (39.58%) | SF4 (12.5%) |
| **C-C-C-H** | 1 | 2 | 1.3 | MetRS (6.1.1.10) (2.170.220.10)<br>Q-insertase (ambiguous) (2.4.2.29) (3.20.20.105) | 99.5 | 0 | 0 | 100 | 0 | 0 | ZN (100.0%) | | |
| **C-C-DE** | 1 | 5 | 1.6 | diaminopimelate epimerase (5.1.1.7) (3.10.310.10)<br>thioredoxin-disulfide reductase (1.8.1.9) (3.50.50.60)<br>ADH3 (1.1.1.284) (3.90.180.10)<br>protein disulfide-isomerase (5.3.4.1) (3.40.30.10)<br>SerRS (6.1.1.11) (3.30.930.10) | 40 | 12.8 | 12 | 40 | 60 | 0 | ZN (27.91%) | FAD (37.21%)<br>NAD (4.65%)<br>FDA (2.33%) | FAD (16.28%)<br>SER (4.65%)<br>ATP (2.33%) |
| **C-C-DE-H** | 1 | 2 | 1.9 | cytidine deaminase (3.5.4.5) (3.40.140.10)<br>anhydrase (4.2.1.1) (3.40.1050.10) | 93 | 0 | 35.5 | 100 | 0 | 0 | ZN (88.0%)<br>CO (2.0%) | | DHZ (2.0%)<br>URI (2.0%)<br>ZEB (2.0%)<br>CTD (2.0%)<br>U (2.0%) |
| **C-C-H** | 1 | 3 | 1.4 | glutathione-disulfide reductase (1.8.1.7) (3.50.50.60)<br>ADH (1.1.1.1) (3.90.180.10)<br>ferredoxin:thioredoxin reductase (1.8.7.2) (3.90.460.10) | 33.3 | 44.7 | 46 | 66.7 | 33.3 | 0 | ZN (31.16%)<br>CD (0.59%)<br>AU (0.3%)<br>CU (0.3%) | FAD (19.29%)<br>NAD (12.17%)<br>NAJ (3.26%)<br>GSH (0.89%)<br>NAP (0.89%)<br>NAI (0.59%)<br>NDP (0.59%)<br>GDS (0.3%)<br>CND (0.3%)<br>8ID (0.3%)<br>Other (0%) | FAD (4.15%)<br>ETF (2.97%)<br>PFB (2.37%)<br>NAD (1.19%)<br>FU2 (0.89%)<br>NAJ (0.89%)<br>NAI (0.89%)<br>ELI (0.59%)<br>PYZ (0.59%)<br>ACT (0.59%)<br>Other (9.56%) |
| **C-C-X** | 1 | 3 | 1.8 | beta-ketothiolase (2.3.1.16) (3.40.47.10)<br>ArsC (ambiguous) (1.20.4.4) (3.40.50.2300)<br>prostaglandin-E synthase (5.3.99.3) (3.40.30.10) | 0 | 4.7 | 0.7 | 0 | 100 | 0 | | COA (33.33%)<br>ACO (33.33%) | PN5 (8.33%)<br>ACO (8.33%)<br>SCY (8.33%) |
| **C-DE-DE** | 2 | 4 | 1.2 | 3 (4.1.99.12) (3.90.870.10)<br>glucosylceramidase (3.2.1.45) (3.20.20.80)<br>MsrA (1.8.4.11) (3.30.1060.10)<br>thiamine pyridinylase (2.5.1.2) (3.40.190.10) | 0 | 0 | 32.8 | 0 | 100 | 0 | | | 5RP (7.41%)<br>3RK (7.41%)<br>IFM (7.41%)<br>HKW (7.41%)<br>RJR (7.41%)<br>NBV (3.7%)<br>3RI (3.7%)<br>NND (3.7%)<br>AMF (3.7%) |

| | | | | | | | | | | | | | |
|---|---|---|---|---|---|---|---|---|---|---|---|---|---|
| | | | | | | | | | | | | | LGS (3.7%)<br>Other (29.6%) |
| **C-DE-H** | 2 | 9 | 1.3 | NO-forming) (1.7.2.1) (2.60.40.420)<br>methylated-DNA---[protein]-cysteine S-methyltransferase (2.1.1.63) (1.10.10.10)<br>arylamine N-acetyltransferase (2.3.1.5) (2.40.128.150)<br>conjugase (3.4.19.9) (3.40.50.880)<br>Unknown (3.4.22.-) (3.40.395.10)<br>carboxymethylenebutenolidase (3.1.1.45) (3.40.50.1820)<br>selenocysteine lyase (4.4.1.16) (3.40.640.10)<br>glutamate racemase (5.1.1.3) (3.40.50.1860)<br>cytosine deaminase (3.5.4.1) (3.40.140.10) | 11.2 | 0 | 4.1 | 22.2 | 66.7 | 11.1 | ZN (55.56%)<br>NA (3.7%) | | DOD (11.11%)<br>FMT (7.41%)<br>DUC (7.41%)<br>HPY (7.41%)<br>ACT (3.7%)<br>GOL (3.7%) |
| **C-DE-H-NQ** | 1 | 2 | 2.5 | adenosylhomocysteinase (3.3.1.1) (3.40.50.720)<br>TRS (6.1.1.3) (3.30.930.10) | 36.5 | 50 | 20.5 | 50 | 50 | 0 | ZN (34.88%)<br>K (2.33%) | NAD (16.28%)<br>NAI (2.33%) | 2CR (9.3%)<br>TSB (9.3%)<br>SSA (6.98%)<br>1B2 (4.65%)<br>409 (2.33%)<br>THR (2.33%)<br>SO4 (2.33%)<br>1B3 (2.33%)<br>BC9 (2.33%)<br>X16 (2.33%)<br>Other (0%) |
| **C-DE-R** | 1 | 2 | 1.4 | MoCu-CODH (1.2.5.3) (3.30.365.10)<br>anhydrase (4.2.1.1) (3.40.1050.10) | 22.5 | 50 | 5 | 100 | 0 | 0 | ZN (35.48%)<br>CU (3.23%)<br>CO (3.23%) | MCN (22.58%)<br>CUB (6.45%) | BCT (12.9%)<br>CUM (6.45%)<br>CUN (6.45%)<br>OMO (3.23%) |
| **C-DE-STY** | 1 | 7 | 1.8 | glutamate racemase (5.1.1.3) (3.40.50.1860)<br>arginine kinase (2.7.3.3) (3.30.590.10)<br>ferredoxin-NADP+-oxidoreductase (1.18.1.2) (3.40.50.80)<br>7-carboxy-7-deazaguanine synthase (4.3.99.3) (3.20.20.70)<br>phosphopentosisomerase (5.3.1.6) (3.40.1400.10)<br>YkvM (1.7.1.13) (3.30.1130.10)<br>vinylacetyl-CoA Delta-isomerase (5.3.3.3) (2.40.110.10) | 0 | 18.3 | 33.3 | 14.3 | 85.7 | 0 | | FAD (32.61%)<br>SAM (13.04%)<br>SF4 (13.04%) | ARG (8.7%)<br>DAR (6.52%)<br>ORN (2.17%)<br>3S5 (2.17%)<br>IOM (2.17%)<br>ILO (2.17%)<br>R5P (2.17%)<br>RB5 (2.17%)<br>AOS (2.17%)<br>R52 (2.17%)<br>Other (6.51%) |
| **C-H-H** | 1 | 4 | 1.3 | GTP cyclohydrolase (3.5.4.16) (3.30.1130.10)<br>PPIase (5.2.1.8) (3.10.50.40)<br>neelaredoxin (1.15.1.2) (2.60.40.730)<br>mannonate dehydratase (4.2.1.8) (3.20.20.150) | 54.5 | 0 | 13 | 75 | 25 | 0 | ZN (33.33%)<br>FE (23.08%)<br>MN (12.82%)<br>CA (5.13%) | | GTP (7.69%)<br>8GT (5.13%)<br>CS2 (5.13%)<br>8DG (2.56%)<br>IPA (2.56%)<br>QBQ (2.56%) |
| **C-H-H-STY** | 1 | 2 | 2.2 | PPIase (5.2.1.8) (3.10.50.40)<br>N-acetylmuramoyl-L-alanine amidase (3.5.1.28) (3.40.80.10) | 50 | 0 | 0 | 0 | 100 | 0 | ZN (100.0%) | | |
| **C-H-NQ** | 2 | 6 | 1.5 | DOCS (2.3.1.74) (3.40.47.10)<br>F1-ATPase (7.1.2.2) (2.170.16.10)<br>cathepsin L (3.4.22.15) (3.90.70.10)<br>beta-lactamase (3.5.2.6) (3.60.15.10)<br>GTP cyclohydrolase (3.5.4.16) (3.30.1130.10)<br>cathepsin K (3.4.22.38) (3.90.70.10) | 10.8 | 0 | 12.3 | 33.3 | 66.7 | 0 | ZN (35.39%)<br>CD (0.91%)<br>CL (0.68%)<br>AU (0.23%) | | MRI (12.79%)<br>RTD (4.34%)<br>*P* (1.6%)<br>E64 (1.37%)<br>ZZ7 (1.14%)<br>OH (0.91%)<br>QUE (0.91%)<br>MYC (0.91%)<br>COA (0.68%)<br>0IW (0.68%)<br>Other (29.66%) |
| **C-H-STY** | 2 | 7 | 1.7 | protein-tyrosine-phosphatase (3.1.3.48) (3.90.190.10)<br>ADH3 (1.1.1.284) (3.90.180.10)<br>MDR (1.1.1.105) (3.90.180.10)<br>ADH (1.1.1.1) (3.90.180.10)<br>3 (4.1.99.12) (3.90.870.10)<br>vinylacetyl-CoA Delta-isomerase (5.3.3.3) (1.20.140.10)<br>destabilase (3.5.1.44) (3.30.1330.200) | 14.3 | 17.9 | 5 | 42.9 | 57.1 | 0 | ZN (6.48%) | NAD (42.59%)<br>NAJ (10.19%)<br>NAP (3.7%)<br>NAI (1.85%)<br>NDP (1.85%)<br>CND (0.93%)<br>8ID (0.93%)<br>PAD (0.93%) | NAD (3.7%)<br>CXF (2.78%)<br>NAJ (2.78%)<br>NAI (2.78%)<br>MRD (1.85%)<br>ETF (1.85%)<br>COD (1.85%)<br>SF4 (1.85%)<br>PVS (0.93%) |

| | | | | | | | | | | | | | |
|---|---|---|---|---|---|---|---|---|---|---|---|---|---|
| | | | | | | | | | | | | | PSY (0.93%)<br>Other (4.65%) |
| **C-NQ-X** | 1 | 3 | 1.6 | peptide deformylase (3.5.1.88) (3.90.45.10)<br>amidophosphoribosyltransferase (2.4.2.14) (3.60.20.10)<br>ferredoxin-glutamate synthase (1.4.7.1) (3.60.20.10) | 33.3 | 0 | 24.3 | 33.3 | 66.7 | 0 | ZN (32.33%)<br>NI (17.24%)<br>CD (5.6%)<br>CO (4.74%)<br>FE (1.72%)<br>FE2 (0.86%)<br>CL (0.43%) | | BB2 (19.4%)<br>FMT (3.02%)<br>ONL (1.72%)<br>LHY (1.29%)<br>K3U (1.29%)<br>K1U (1.29%)<br>K2U (1.29%)<br>SB7 (0.86%)<br>MLN (0.86%)<br>PO4 (0.86%)<br>Other (3.87%) |
| **C-NQ-X-X-X** | 1 | 2 | 1.6 | hexosephosphate aminotransferase (2.6.1.16) (3.60.20.10)<br>GOGAT (1.4.1.13) (3.60.20.10) | 0 | 0 | 19.5 | 0 | 100 | 0 | | | ONL (100.0%) |
| **C-R-STY** | 1 | 3 | 1.7 | sulfite oxidase (1.8.3.1) (3.90.420.10)<br>McyF (5.1.1.13) (3.40.50.1860)<br>protein-tyrosine-phosphatase (3.1.3.48) (3.90.190.10) | 33.3 | 0 | 42.7 | 33.3 | 66.7 | 0 | MO (5.41%) | | SO4 (5.41%)<br>073 (5.41%)<br>PO4 (5.41%)<br>UA1 (5.41%)<br>234 (2.7%)<br>OTA (2.7%)<br>IX1 (2.7%)<br>F20 (2.7%)<br>F32 (2.7%)<br>DBD (2.7%)<br>Other (45.9%) |
| **C-STY-STY** | 1 | 5 | 1.3 | pyruvate (2.7.9.1) (3.20.20.60)<br>(S)-oxynitrilase (4.1.2.47) (3.40.50.1820)<br>GDP-4-keto-6-deoxy-D-mannose-3 (1.1.1.271) (3.40.50.720)<br>riboflavin synthase (2.5.1.9) (2.40.30.20)<br>ribonucleoside-triphosphate reductase (formate) (1.1.98.6)<br>(3.20.70.20) | 0 | 2.8 | 34.6 | 0 | 100 | 0 | | NAP (8.7%) | ACN (13.04%)<br>SCN (8.7%)<br>GOL (8.7%)<br>GFB (8.7%)<br>CNH (4.35%)<br>MNN (4.35%)<br>FAC (4.35%)<br>ACT (4.35%)<br>CLX (4.35%)<br>ICN (4.35%)<br>Other (26.1%) |
| **C-STY-X** | 1 | 2 | 1.2 | N-carbamoylsarcosine amidase (3.5.1.59) (3.40.50.850)<br>adenosylmethionine decarboxylase (4.1.1.50) (3.60.90.10) | 0 | 0 | 0 | 0 | 100 | 0 | | | SO4 (100.0%) |
| **C-X-X** | 1 | 3 | 1.7 | D-camphor-exo-hydroxylase (1.14.15.1) (1.10.630.10)<br>beta-hydroxyoctanoyl-ACP-dehydrase (4.2.1.59) (3.40.47.10)<br>rhodanese (2.8.1.1) (3.40.250.10) | 0 | 27 | 2 | 33.3 | 66.7 | 0 | | HEM (53.26%)<br>7HE (1.09%)<br>MNR (1.09%) | HEM (11.96%)<br>CER (5.43%)<br>OCA (2.17%)<br>6NA (2.17%)<br>MRJ (2.17%)<br>PNS (2.17%)<br>TLM (2.17%)<br>HEC (1.09%)<br>CMO (1.09%)<br>1XG (1.09%)<br>Other (7.63%) |
| **C-X-X-X-X** | 1 | 2 | 1.1 | ArsC (ambiguous) (1.20.4.4) (3.40.50.2300)<br>rhodanese (2.8.1.1) (3.40.250.10) | 0 | 0 | 12.5 | 0 | 100 | 0 | | | SO4 (50.0%)<br>ACT (50.0%) |
| **DE-DE-DE** | 4 | 18 | 1.4 | calcium pump (7.2.2.10) (3.40.50.1000)<br>neopullulanase (3.2.1.135) (3.20.20.80)<br>4-alpha-glucanotransferase (2.4.1.25) (3.20.20.80)<br>exodeoxyribonuclease (lambda-induced) (3.1.11.3) (3.40.50.1010)<br>glucoamylase (3.2.1.3) (1.50.10.10)<br>fumarylacetoacetase (3.7.1.2) (3.90.850.10)<br>xylose isomerase (5.3.1.5) (3.20.20.150)<br>enolase (4.2.1.11) (3.20.20.120)<br>2-chloro-2 (5.5.1.7) (3.20.20.120)<br>duplicase (2.7.7.7) (3.30.460.10)<br>transphosphoribosidase (2.4.2.8) (3.40.50.2020)<br>Unknown (2.4.2.-) (3.40.50.2020)<br>protoporphyrin ferrochelatase (4.99.1.1) (3.40.50.1400)<br>glycopeptidase (3.5.1.52) (2.60.120.230)<br>glucoamylase (3.2.1.3) (1.50.10.10)<br>isoamylase (3.2.1.68) (3.20.20.80)<br>exodeoxyribonuclease (lambda-induced) (3.1.11.3) (3.40.50.1010)<br>UDP-N-acetyl-D-glucosamine 2-epimerase (5.1.3.14)<br>(3.40.50.2000) | 24.4 | 0 | 26.8 | 50 | 50 | 0 | MG (26.19%)<br>MN (10.48%)<br>CA (1.75%)<br>CO (0.58%)<br>ZN (0.23%)<br>D8U (0.12%)<br>CL (0.12%) | | *P* (9.31%)<br>GLC (3.96%)<br>MF4 (2.44%)<br>ALF (1.51%)<br>XYL (1.51%)<br>BGC (0.93%)<br>EDO (0.93%)<br>DUP (0.93%)<br>8OG (0.81%)<br>ACP (0.7%)<br>Other (27.7%) |

| | | | | | | | | | | | | |
|---|---|---|---|---|---|---|---|---|---|---|---|---|
| **DE-DE-DE-DE** | 1 | 12 | 2.2 | NAD+-malic enzyme (1.1.1.38) (3.40.50.10380)<br>xylose isomerase (5.3.1.5) (3.20.20.150)<br>Xaa-Pro aminopeptidase (3.4.11.9) (3.90.230.10)<br>3'(2') (3.1.3.7) (3.40.190.80)<br>alpha-1 (4.2.2.2) (2.160.20.10)<br>3'(2') (3.1.3.7) (3.40.190.80)<br>fructose-bisphosphatase (3.1.3.11) (3.30.540.10)<br>swivelase (5.6.2.1) (3.40.50.140)<br>exodeoxyribonuclease (lambda-induced) (3.1.11.3) (3.40.50.1010)<br>lysozyme (3.2.1.17) (3.20.20.80)<br>inorganic diphosphatase (3.6.1.1) (3.90.80.10)<br>non-chaperonin molecular chaperone ATPase (3.6.4.10)<br>(3.30.30.30) | 53.5 | 2.1 | 38.2 | 83.3 | 8.3 | 8.3 | MN (22.11%)<br>MG (20.35%)<br>CA (4.27%)<br>ZN (4.02%)<br>TL (2.01%)<br>CO (1.76%)<br>NA (0.75%)<br>K (0.25%) | NAD (1.01%)<br>NAP (0.25%) | PO4 (6.28%)<br>XYL (3.27%)<br>GLC (2.76%)<br>EDO (1.51%)<br>PRO (1.51%)<br>AMP (1.51%)<br>XLS (1.26%)<br>OXL (1.01%)<br>TTN (1.01%)<br>GOL (1.01%)<br>Other (11.51%) |
| **DE-DE-DE-DE-H** | 2 | 7 | 1.7 | neopullulanase (3.2.1.135) (3.20.20.80)<br>L-fuconate dehydratase (4.2.1.68) (3.20.20.120)<br>GS (6.3.1.2) (3.30.590.10)<br>arginase (3.5.3.1) (3.40.800.10)<br>mandelate racemase (5.1.2.2) (3.20.20.120)<br>exodeoxyribonuclease III (3.1.11.2) (3.60.10.10)<br>cis (5.5.1.1) (3.20.20.120) | 39.4 | 0 | 24.3 | 85.7 | 14.3 | 0 | MN (48.1%)<br>MG (11.96%)<br>CO (2.72%)<br>NI (1.09%)<br>CA (0.82%)<br>ZN (0.54%) | | ABH (2.99%)<br>P3S (2.17%)<br>GLC (1.9%)<br>S2C (1.9%)<br>ORN (1.36%)<br>NNH (1.09%)<br>X7A (1.09%)<br>NOJ (0.82%)<br>AC1 (0.82%)<br>HAR (0.82%)<br>Other (17.83%) |
| **DE-DE-DE-DE-H-H** | 2 | 7 | 1.9 | bacterial leucyl aminopeptidase (3.4.11.10) (3.40.630.10)<br>glutamate carboxypeptidase (3.4.17.11) (3.40.630.10)<br>arginase (3.5.3.1) (3.40.800.10)<br>L-rhamnose isomerase (5.3.1.14) (3.20.20.150)<br>catalase (1.11.1.6) (1.20.1260.10)<br>methane hydroxylase (1.14.13.25) (1.10.620.20)<br>redoxyendonuclease (3.1.21.2) (3.20.20.150) | 71.9 | 0 | 32.3 | 100 | 0 | 0 | MN (31.67%)<br>FE (23.58%)<br>ZN (11.02%)<br>CO (2.41%)<br>FE2 (1.55%)<br>MN3 (1.38%)<br>NI (0.69%)<br>O (0.34%) | | ABH (1.89%)<br>S2C (1.2%)<br>OH (1.2%)<br>ORN (0.86%)<br>LEU (0.69%)<br>BES (0.69%)<br>NNH (0.69%)<br>X7A (0.69%)<br>TRS (0.52%)<br>PLU (0.52%)<br>Other (12.43%) |
| **DE-DE-DE-DE-K** | 1 | 2 | 2.3 | cis (5.5.1.1) (3.20.20.120)<br>OSBS (4.2.1.113) (3.20.20.120) | 73.5 | 0 | 29.5 | 100 | 0 | 0 | MG (39.74%)<br>MN (11.54%)<br>CL (1.28%) | | MUC (7.69%)<br>DGL (6.41%)<br>TYR (3.85%)<br>ACY (2.56%)<br>AME (2.56%)<br>DLY (1.28%)<br>SUG (1.28%)<br>VAL (1.28%)<br>ALA (1.28%)<br>SMG (1.28%)<br>Other (7.68%) |
| **DE-DE-DE-DE-X** | 1 | 2 | 0.7 | fructose-bisphosphatase (3.1.3.11) (3.30.540.10)<br>D-galactose 1-phosphate phosphatase (3.1.3.94) (3.30.540.10) | 81 | 2.5 | 69 | 100 | 0 | 0 | MG (26.4%)<br>CA (13.71%)<br>MN (10.15%)<br>ZN (7.11%)<br>TL (5.08%)<br>GD (2.03%)<br>CL (0.51%) | NAP (1.52%) | PO4 (16.75%)<br>IPD (4.57%)<br>F6P (3.05%)<br>SO4 (2.03%)<br>PI (1.02%)<br>PO3 (1.02%)<br>FBP (1.02%)<br>FDP (0.51%)<br>INS (0.51%)<br>LIP (0.51%)<br>Other (1.02%) |
| **DE-DE-DE-H** | 2 | 12 | 1.7 | stearoyl-[acyl-carrier-protein] 9-desaturase (1.14.19.2)<br>(1.10.620.20)<br>L-fuconate dehydratase (4.2.1.68) (3.20.20.120)<br>caffeate O-methyltransferase (2.1.1.68) (3.40.50.150)<br>creatininase (3.5.2.10) (3.40.50.10310)<br>diisopropyl-fluorophosphatase (3.1.8.2) (2.120.10.30)<br>mandelate racemase (5.1.2.2) (3.20.20.120)<br>exo-cellobiohydrolase (3.2.1.91) (2.70.100.10)<br>methane hydroxylase (1.14.13.25) (1.10.620.20)<br>3 (4.1.99.12) (3.90.870.10)<br>duplicase (2.7.7.7) (3.30.420.10)<br>Unknown (3.1.-.-) (3.30.420.10)<br>cis (5.5.1.1) (3.20.20.120) | 42.3 | 0 | 18.4 | 83.3 | 16.7 | 0 | FE (35.46%)<br>MG (13.52%)<br>MN (4.59%)<br>FE2 (2.55%)<br>ZN (1.28%)<br>CA (0.77%)<br>CO (0.51%)<br>NA (0.51%)<br>CL (0.26%)<br>ER3 (0.26%)<br>Other (0%) | | BGC (4.34%)<br>FMT (0.77%)<br>*P* (0.77%)<br>MSE (0.51%)<br>MCR (0.51%)<br>AZI (0.51%)<br>PER (0.51%)<br>PCR (0.51%)<br>SO4 (0.51%)<br>TMP (0.51%)<br>Other (14.0%) |

| | | | | | | | | | | | | | |
|---|---|---|---|---|---|---|---|---|---|---|---|---|---|
| **DE-DE-DE-H-H** | 1 | 5 | 1.6 | DNase (3.1.21.1) (3.60.10.10)<br>exo-1 (3.2.1.114) (3.20.110.10)<br>tripeptidase (3.4.11.4) (1.10.390.10)<br>arginase (3.5.3.1) (3.40.800.10)<br>methane hydroxylase (1.14.13.25) (1.10.620.20) | 79 | 0 | 18.4 | 100 | 0 | 0 | ZN (22.43%)<br>FE (21.34%)<br>MN (14.8%)<br>FE2 (1.4%)<br>CO (1.09%)<br>NI (0.31%)<br>MG (0.16%) | | ABH (1.71%)<br>BES (1.4%)<br>S2C (1.09%)<br>LYS (0.78%)<br>ORN (0.78%)<br>NNH (0.62%)<br>X7A (0.62%)<br>HAR (0.47%)<br>MSN (0.31%)<br>SWA (0.31%)<br>Other (24.2%) |
| **DE-DE-DE-H-H-H** | 1 | 2 | 1.4 | Unknown (1.-.-.-) (3.60.15.10)<br>creatininase (3.5.2.10) (3.40.50.10310) | 61 | 0 | 5 | 100 | 0 | 0 | ZN (29.55%)<br>MN (15.91%)<br>FE (9.09%)<br>O (2.27%)<br>CL (2.27%) | | FEO (15.91%)<br>OXY (9.09%)<br>CRN (2.27%) |
| **DE-DE-DE-H-H-H-STY** | 1 | 2 | 0.4 | phosphomonoesterase (3.1.3.1) (3.40.720.10)<br>2 (5.4.2.12) (3.40.720.10) | 31 | 0 | 92.5 | 100 | 0 | 0 | ZN (29.63%)<br>CO (7.41%) | | PO4 (33.33%)<br>2PG (7.41%)<br>3PG (7.41%)<br>PAE (3.7%)<br>MMQ (3.7%)<br>AF3 (3.7%)<br>WO4 (3.7%) |
| **DE-DE-DE-H-K** | 1 | 2 | 0.8 | Unknown (4.6.1.-) (3.20.20.190)<br>glycerophosphodiester phosphodiesterase (3.1.4.46)<br>(3.20.20.190) | 61.5 | 0 | 7.5 | 100 | 0 | 0 | MG (33.33%)<br>CA (11.11%)<br>FE (3.7%) | | GOL (11.11%)<br>IPD (3.7%) |
| **DE-DE-DE-H-NQ** | 1 | 2 | 1.9 | beta-methylaspartase (4.3.1.2) (3.20.20.120)<br>N-sulfoglucosamine sulfohydrolase (3.10.1.1) (3.40.720.10) | 87.5 | 0 | 6 | 100 | 0 | 0 | MG (46.15%)<br>CA (23.08%) | | FGP (23.08%)<br>2AS (7.69%) |
| **DE-DE-DE-H-R** | 1 | 4 | 2.5 | konchizaimu (2.4.1.19) (3.20.20.80)<br>MICL (4.1.3.30) (3.20.20.60)<br>Xaa-Pro aminopeptidase (3.4.11.9) (3.90.230.10)<br>endo-polygalacturonase (3.2.1.15) (2.160.20.10) | 0 | 0 | 13.8 | 50 | 50 | 0 | | | GLC (16.67%)<br>PRO (10.0%)<br>ACI (6.67%)<br>*P* (6.67%)<br>BGC (5.0%)<br>NOJ (5.0%)<br>ADH (5.0%)<br>M44 (3.33%)<br>GTR (3.33%)<br>OPG (1.67%)<br>Other (10.02%) |
| **DE-DE-DE-K** | 3 | 13 | 1.7 | enolase (4.2.1.11) (3.20.20.120)<br>OSBS (4.2.1.113) (3.20.20.120)<br>Unknown (3.6.1.74) (3.20.100.10)<br>OSBS (4.2.1.113) (3.20.20.120)<br>Unknown (5.5.1.-) (3.20.20.120)<br>NADP+-ICDH (1.1.1.42) (3.40.718.10)<br>L-fuconate dehydratase (4.2.1.68) (3.20.20.120)<br>mandelate racemase (5.1.2.2) (3.20.20.120)<br>swivelase (5.6.2.1) (3.40.50.140)<br>2-chloro-2 (5.5.1.7) (3.20.20.120)<br>beta-phosphoglucomutase (5.4.2.6) (3.40.50.1000)<br>licheninase (3.2.1.73) (3.20.20.80)<br>endo-1 (3.2.1.39) (3.20.20.80) | 44.2 | 0.2 | 23.4 | 84.6 | 15.4 | 0 | MG (40.82%)<br>MN (5.1%)<br>CA (3.32%)<br>CL (0.51%) | NAD (0.77%) | IPM (4.59%)<br>MUC (2.55%)<br>ICT (2.55%)<br>DGL (2.04%)<br>ALF (2.04%)<br>MGF (2.04%)<br>TYR (1.79%)<br>APG (1.28%)<br>AKG (1.02%)<br>LGT (1.02%)<br>Other (20.28%) |
| **DE-DE-DE-K-K** | 1 | 2 | 0.8 | Hyp-B 2-epimerase (5.1.1.22) (3.20.20.120)<br>Unknown (5.5.1.-) (3.20.20.120) | 60.5 | 0 | 27.5 | 100 | 0 | 0 | MG (48.31%)<br>MN (2.25%)<br>CL (1.12%) | | 0XW (5.62%)<br>TYR (4.49%)<br>GLU (3.37%)<br>MUC (3.37%)<br>PBE (2.25%)<br>5CR (2.25%)<br>DGL (2.25%)<br>NPQ (2.25%)<br>4OP (1.12%)<br>NSK (1.12%)<br>Other (12.32%) |
| **DE-DE-DE-NQ-Z** | 1 | 2 | 0.3 | cerebroside-sulfatase (3.1.6.8) (3.40.720.10)<br>arylsulfatase (3.1.6.1) (3.40.720.10) | 96 | 0 | 23 | 100 | 0 | 0 | CA (35.71%)<br>MG (2.38%) | | DDZ (19.05%)<br>FGP (9.52%)<br>SO4 (9.52%)<br>ALS (4.76%)<br>SV7 (4.76%)<br>FGL (2.38%)<br>62Y (2.38%) |

| | | | | | | | | | | | | | |
|---|---|---|---|---|---|---|---|---|---|---|---|---|---|
| **DE-DE-DE-STY** | 3 | 13 | 2.3 | (+)-bornylpyrophosphate cyclase (5.5.1.8) (1.10.600.10)<br>levansucrase (2.4.1.10) (2.115.10.20)<br>IAG-NH (3.2.2.1) (3.90.245.10)<br>cellulase (3.2.1.4) (1.50.10.10)<br>duplicase (2.7.7.7) (3.30.70.270)<br>beta-glucuronidase (3.2.1.31) (2.60.120.260)<br>exo-cellobiohydrolase (3.2.1.91) (3.20.20.40)<br>chitinase (3.2.1.14) (3.20.20.80)<br>biliverdin reductase (1.3.1.24) (3.30.360.10)<br>chitinase (3.2.1.14) (3.20.20.80)<br>trans-deoxyribosylase (2.4.2.6) (3.40.50.450)<br>sedolisin (3.4.21.100) (3.40.50.200)<br>inorganic diphosphatase (3.6.1.1) (3.90.80.10) | 16.5 | 0 | 25.6 | 30.8 | 69.2 | 0 | CA (4.35%)<br>MG (1.45%)<br>IOD (0.48%) | | AMI (5.8%)<br>IPT (4.83%)<br>NAG (4.83%)<br>149 (4.35%)<br>*P* (3.38%)<br>VR0 (2.9%)<br>NGT (2.9%)<br>A1L (2.42%)<br>GLC (1.93%)<br>3AG (1.45%)<br>Other (45.85%) |
| **DE-DE-DE-X** | 1 | 3 | 1.8 | hybridase (3.1.26.4) (3.30.420.10)<br>thermonuclease (3.1.31.1) (2.40.50.90)<br>ADPR-PPase (3.6.1.13) (3.90.79.10) | 57.7 | 0 | 54 | 100 | 0 | 0 | CA (48.0%)<br>MN (1.6%)<br>MG (0.8%)<br>CO (0.8%) | | THP (47.2%)<br>PO4 (0.8%)<br>ADV (0.8%) |
| **DE-DE-H** | 3 | 25 | 1.3 | glycine amidinotransferase (2.1.4.1) (3.75.10.10)<br>haloalkane dehalogenase (3.8.1.5) (3.40.50.1820)<br>exo-cellobiohydrolase (3.2.1.91) (3.20.20.80)<br>glyoxylate reductase (1.1.1.26) (3.40.50.720)<br>Unknown (3.4.24.-) (3.30.830.10)<br>Transferred to 3.3.2.9 and 3.3.2.10 (3.3.2.3) (3.40.50.1820)<br>PRMT1 (gene name) (2.1.1.319) (3.40.50.150)<br>YkvM (1.7.1.13) (3.30.1130.10)<br>3'-phosphoadenylyl-sulfate:[heparan sulfate]-glucosamine 3-sulfotransferase (2.8.2.30) (3.40.50.300)<br>chloroperoxidase (1.11.1.10) (1.10.489.10)<br>phosphatidase (3.1.1.4) (1.20.90.10)<br>glucosamine-6-phosphate deaminase (3.5.99.6) (3.40.50.1360)<br>L-fucose isomerase (5.3.1.25) (3.20.14.10)<br>carboxypeptidase T (3.4.17.18) (3.40.630.10)<br>Unknown (1.-.-.-) (3.60.15.10)<br>galactarate dehydratase (D-threo-forming) (4.2.1.158) (3.20.20.120)<br>Unknown (2.4.1.-) (3.90.550.10)<br>UDP-sugar diphosphatase (3.6.1.45) (3.60.21.10)<br>arylsulfatase (3.1.6.1) (3.40.720.10)<br>Unknown (3.1.5.-) (1.10.3210.10)<br>protoporphyrin ferrochelatase (4.99.1.1) (3.40.50.1400)<br>aldolase (4.1.2.13) (3.20.20.70)<br>enolase (4.2.1.11) (3.20.20.120)<br>creatinase (3.5.3.3) (3.90.230.10)<br>PdxJ (2.6.99.2) (3.20.20.70) | 25.5 | 5.5 | 25.1 | 48 | 52 | 0 | ZN (30.61%)<br>MN (5.54%)<br>CA (3.5%)<br>FE (0.58%)<br>MG (0.58%)<br>CL (0.29%)<br>HG (0.29%)<br>O (0.29%) | HEM (4.37%)<br>PXP (0.87%) | *P* (4.66%)<br>DDZ (2.33%)<br>FEO (1.75%)<br>FOC (1.46%)<br>CXA (1.17%)<br>OXY (1.17%)<br>SO4 (1.17%)<br>IPA (0.87%)<br>3K0 (0.87%)<br>GOL (0.58%)<br>Other (30.16%) |
| **DE-DE-H-H** | 1 | 16 | 2 | redoxyendonuclease (3.1.21.2) (3.20.20.150)<br>thermolysin (3.4.24.27) (1.10.390.10)<br>isopentenyl-diphosphate Delta-isomerase (5.3.3.2) (3.90.79.10)<br>deuterolysin (3.4.24.39) (3.40.390.10)<br>neprilysin (3.4.24.11) (3.40.390.10)<br>bontoxilysin (3.4.24.69) (3.90.1240.10)<br>peptidyl-Lys metalloendopeptidase (3.4.24.20) (3.40.390.10)<br>dGTPase (3.1.5.1) (1.10.3210.10)<br>redoxyendonuclease (3.1.21.2) (3.20.20.150)<br>alpha-toxin (3.1.4.3) (1.10.575.10)<br>DNase (3.1.21.1) (3.60.10.10)<br>NDO (1.14.12.12) (3.90.380.10)<br>nicotinamidase (3.5.1.19) (3.40.50.850)<br>Xaa-Pro aminopeptidase (3.4.11.9) (3.90.230.10)<br>Unknown (3.4.24.-) (3.30.830.10)<br>2 (5.4.2.12) (3.40.720.10) | 69.3 | 0 | 32.4 | 100 | 0 | 0 | ZN (48.85%)<br>MN (2.52%)<br>CD (1.61%)<br>CU (0.46%)<br>FE2 (0.46%)<br>FE (0.46%)<br>CU1 (0.23%)<br>CL (0.23%)<br>NA (0.23%)<br>RU (0.23%)<br>Other (0.23%) | | *P* (4.82%)<br>VAL (3.21%)<br>RDF (1.38%)<br>LEU (0.92%)<br>GOL (0.92%)<br>BNL (0.92%)<br>OXY (0.92%)<br>MRD (0.69%)<br>NX6 (0.69%)<br>ASP (0.69%)<br>Other (23.92%) |
| **DE-DE-H-H-H** | 1 | 4 | 2.2 | alpha-toxin (3.1.4.3) (1.10.575.10)<br>hydroxyacylglutathione hydrolase (3.1.2.6) (3.60.15.10)<br>neprilysin (3.4.24.11) (3.40.390.10)<br>adenosine deaminase (3.5.4.4) (3.20.20.140) | 94 | 2.5 | 30.8 | 100 | 0 | 0 | ZN (60.65%)<br>FE (12.26%)<br>CD (2.58%)<br>CO (0.65%)<br>NI (0.65%) | GBP (0.65%) | RDF (3.23%)<br>PRH (1.94%)<br>DCF (1.29%)<br>ADE (1.29%)<br>MCF (1.29%)<br>LAC (0.65%)<br>TLA (0.65%)<br>CAC (0.65%)<br>ACY (0.65%)<br>OIR (0.65%)<br>Other (10.4%) |

| | | | | | | | | | | | | | |
|---|---|---|---|---|---|---|---|---|---|---|---|---|---|
| **DE-DE-H-H-STY** | 1 | 2 | 1.3 | neurolysin (3.4.24.16) (3.40.390.10)<br>lethal toxin (3.4.24.83) (3.40.390.10) | 93.5 | 0 | 13 | 50 | 50 | 0 | ZN (53.49%) | | *P* (2.33%)<br>2ZL (2.33%)<br>GOL (2.33%)<br>56R (2.33%)<br>GM6 (2.33%)<br>407 (2.33%)<br>30H (2.33%)<br>56Q (2.33%)<br>0LX (2.33%)<br>30P (2.33%)<br>Other (23.3%) |
| **DE-DE-H-NQ** | 2 | 13 | 2 | 1-phosphatidylinositol-4 (3.1.4.11) (3.20.20.190)<br>phosphomonoesterase (3.1.3.2) (3.60.21.10)<br>exodeoxyribonuclease III (3.1.11.2) (3.60.10.10)<br>Unknown (4.6.1.-) (3.20.20.190)<br>1-phosphatidylinositol-4 (3.1.4.11) (3.20.20.190)<br>Unknown (3.1.5.-) (1.10.3210.10)<br>histidinol dehydrogenase (1.1.1.23) (3.40.50.1980)<br>beta-methylaspartase (4.3.1.2) (3.20.20.120)<br>UDP-sugar diphosphatase (3.6.1.45) (3.60.21.10)<br>cerebroside-sulfatase (3.1.6.8) (3.40.720.10)<br>UDP-sugar diphosphatase (3.6.1.45) (3.60.21.10)<br>Unknown (3.1.-.-) (3.40.1800.10)<br>exo-cellobiohydrolase (3.2.1.91) (3.20.20.80) | 53.1 | 0 | 26.2 | 84.6 | 15.4 | 0 | CA (15.32%)<br>MG (11.91%)<br>ZN (8.51%)<br>FE (5.96%)<br>MN (2.55%)<br>SM (1.7%)<br>LA (0.85%)<br>PB (0.85%)<br>BA (0.43%)<br>F (0.43%)<br>Other (0%) | | XYP (8.94%)<br>3DR (5.96%)<br>*P* (4.68%)<br>I3P (2.55%)<br>DV3 (2.55%)<br>PO4 (2.13%)<br>SO4 (1.28%)<br>XDN (1.28%)<br>XIF (1.28%)<br>IP2 (0.85%)<br>Other (15.81%) |
| **DE-DE-H-R** | 1 | 4 | 2 | fumarylacetoacetase (3.7.1.2) (3.90.850.10)<br>3-phosphoshikimate 1-carboxyvinyltransferase (2.5.1.19) (3.65.10.10)<br>beta-galactosidase (3.2.1.23) (3.20.20.80)<br>neprilysin (3.4.24.11) (3.40.390.10) | 20.2 | 0 | 24.8 | 75 | 25 | 0 | ZN (13.68%) | | FMT (7.37%)<br>GPJ (5.26%)<br>BGC (5.26%)<br>G2F (5.26%)<br>RDF (5.26%)<br>BGP (4.21%)<br>GIM (3.16%)<br>GOX (3.16%)<br>GPF (2.11%)<br>BG6 (2.11%)<br>Other (33.61%) |
| **DE-DE-H-STY** | 1 | 6 | 2.1 | DNase (3.1.21.1) (3.60.10.10)<br>phosphomonoesterase (3.1.3.1) (3.40.720.10)<br>stearoyl-[acyl-carrier-protein] 9-desaturase (1.14.19.2) (1.10.620.20)<br>thermolysin (3.4.24.27) (1.10.390.10)<br>2 (5.4.2.12) (3.40.720.10)<br>SPT (2.3.1.50) (3.40.640.10) | 46.8 | 6.2 | 35.2 | 83.3 | 16.7 | 0 | ZN (19.82%)<br>FE (7.21%)<br>FE2 (5.41%)<br>MN (3.6%)<br>CO (1.8%) | PLP (6.31%) | LYS (10.81%)<br>PO4 (8.11%)<br>PLP (3.6%)<br>*P* (2.7%)<br>2PG (1.8%)<br>3PG (1.8%)<br>PAE (0.9%)<br>MMQ (0.9%)<br>AF3 (0.9%)<br>WO4 (0.9%)<br>Other (22.5%) |
| **DE-DE-H-X** | 1 | 2 | 2 | Unknown (3.2.-.-) (3.90.245.10)<br>phosphatidase (3.1.1.4) (1.20.90.10) | 16.5 | 0 | 20 | 100 | 0 | 0 | CA (20.0%) | | PIR (20.0%)<br>DNB (20.0%)<br>NOS (10.0%)<br>BDR (10.0%)<br>RIB (10.0%) |
| **DE-DE-K** | 2 | 12 | 1.4 | QAPRTase (2.4.2.19) (3.20.20.70)<br>methionine adenosyltransferase (2.5.1.6) (3.30.300.10)<br>(2E (2.5.1.10) (1.10.600.10)<br>lysine 2 (5.4.3.2) (3.20.20.70)<br>type II site-specific deoxyribonuclease (3.1.21.4) (3.40.91.10)<br>phosphoserine phosphatase (3.1.3.3) (3.40.50.1000)<br>isomeroreductase (1.1.1.86) (1.10.1040.10)<br>type II site-specific deoxyribonuclease (3.1.21.4) (3.40.210.10)<br>2-chloro-2 (5.5.1.7) (3.20.20.120)<br>nicotinamidase (3.5.1.19) (3.40.50.850)<br>OMP-DC (4.1.1.23) (3.20.20.70)<br>NADPH-hydroxymethylglutaryl-CoA reductase (1.1.1.34) (3.90.770.10) | 29 | 8.3 | 56.8 | 66.7 | 33.3 | 0 | MG (42.21%)<br>MN (4.02%)<br>CL (2.01%)<br>CA (1.01%)<br>K (0.5%)<br>NA (0.5%) | NAP (1.01%)<br>HMG (0.5%) | NCN (3.52%)<br>*P* (3.02%)<br>MSE (2.01%)<br>PO4 (2.01%)<br>HIO (2.01%)<br>MUC (2.01%)<br>TYR (2.01%)<br>40E (1.51%)<br>DGL (1.51%)<br>U5P (1.51%)<br>Other (23.59%) |
| **DE-DE-K-K-NQ** | 1 | 2 | 0.7 | mandelate racemase (5.1.2.2) (3.20.20.120)<br>glucarate dehydratase (4.2.1.40) (3.20.20.120) | 51 | 0 | 23 | 100 | 0 | 0 | MG (44.16%)<br>NA (2.6%)<br>CA (1.3%)<br>MN (1.3%) | | GLR (5.19%)<br>GKR (3.9%)<br>APG (2.6%)<br>DXL (2.6%)<br>LGT (2.6%)<br>XYH (2.6%)<br>ACT (1.3%)<br>BHO (1.3%)<br>0YR (1.3%) |

| | | | | | | | | | | | | | |
|---|---|---|---|---|---|---|---|---|---|---|---|---|---|
| | | | | | | | | | | | | | 3PY (1.3%)<br>Other (15.6%) |
| **DE-DE-K-NQ** | 1 | 4 | 1.6 | cerebroside-sulfatase (3.1.6.8) (3.40.720.10)<br>beta-methylaspartase (4.3.1.2) (3.20.20.120)<br>catechol O-methyltransferase (2.1.1.6) (3.40.50.150)<br>D-N-carbamoylase (3.5.1.77) (3.60.110.10) | 47.5 | 3.8 | 5 | 75 | 25 | 0 | MG (37.72%)<br>NA (3.51%)<br>CA (2.63%)<br>K (1.75%) | SAM (6.14%)<br>SAH (0.88%)<br>SFG (0.88%) | DNC (2.63%)<br>619 (2.63%)<br>FGP (1.75%)<br>FGL (0.88%)<br>ALS (0.88%)<br>2AS (0.88%)<br>72W (0.88%)<br>611 (0.88%)<br>76T (0.88%)<br>7JX (0.88%)<br>Other (33.44%) |
| **DE-DE-K-STY** | 1 | 2 | 1.6 | NADP+-ICDH (1.1.1.42) (3.40.718.10)<br>AvDH1 (3.6.4.12) (3.40.50.300) | 14.5 | 0 | 16.5 | 100 | 0 | 0 | MG (23.44%)<br>CA (10.94%)<br>ZN (3.12%)<br>MN (3.12%) | | ICT (17.19%)<br>IPM (17.19%)<br>SO4 (6.25%)<br>AKG (3.12%)<br>ICA (3.12%)<br>OXL (1.56%)<br>XYM (1.56%)<br>48Y (1.56%)<br>CIT (1.56%)<br>GOL (1.56%)<br>Other (1.56%) |
| **DE-DE-K-STY-STY-X** | 1 | 2 | 0.8 | SRR (5.1.1.18) (3.40.50.1100)<br>SRR (5.1.1.18) (3.40.50.1100) | 0 | 0 | 0 | 100 | 0 | 0 | | | |
| **DE-DE-NQ** | 1 | 12 | 1.7 | adenosylhomocysteinase (3.3.1.1) (3.40.50.1480)<br>exodeoxyribonuclease III (3.1.11.2) (3.60.10.10)<br>glucarate dehydratase (4.2.1.40) (3.20.20.120)<br>exodeoxyribonuclease III (3.1.11.2) (3.60.10.10)<br>kanamycin kinase (2.7.1.95) (3.90.1200.10)<br>SerRS (6.1.1.11) (3.30.930.10)<br>methanol dehydrogenase (1.1.2.7) (2.140.10.10)<br>biotin carboxylase (6.3.4.14) (3.30.470.20)<br>catechol O-methyltransferase (2.1.1.6) (3.40.50.150)<br>lysozyme (3.2.1.17) (1.10.530.10)<br>anhydrase (4.2.1.1) (2.160.10.10)<br>squalene---hopene cyclase (5.4.99.17) (1.50.10.20) | 28 | 5.8 | 21.9 | 66.7 | 33.3 | 0 | MG (28.94%)<br>CA (8.06%)<br>NA (1.1%)<br>CD (1.1%)<br>PR (0.73%)<br>RU (0.73%)<br>CE (0.37%)<br>EU (0.37%)<br>CO (0.37%)<br>MN (0.37%)<br>Other (0%) | PQQ (6.23%)<br>SAM (1.83%)<br>SAH (0.37%) | PQQ (3.66%)<br>KAN (2.93%)<br>GLR (1.47%)<br>ADP (1.47%)<br>GKR (1.1%)<br>ATP (1.1%)<br>DNC (1.1%)<br>619 (1.1%)<br>SO4 (1.1%)<br>BCT (1.1%)<br>Other (20.69%) |
| **DE-DE-NQ-NQ** | 1 | 2 | 1.9 | diisopropyl-fluorophosphatase (3.1.8.2) (2.120.10.30)<br>aryldialkylphosphatase (3.1.8.1) (2.120.10.30) | 100 | 0 | 0 | 100 | 0 | 0 | CA (69.57%) | | DOD (21.74%)<br>DI9 (4.35%) |
| **DE-DE-NQ-R** | 1 | 4 | 1.1 | glutathione synthase (6.3.2.3) (3.30.470.20)<br>ABL (2.7.10.2) (1.10.510.10)<br>endo-1 (3.2.1.78) (3.20.20.80)<br>D-alanine---D-alanine ligase (6.3.2.4) (3.30.470.20) | 48 | 0 | 49 | 75 | 25 | 0 | MG (37.78%) | | ADP (8.89%)<br>*P* (6.67%)<br>ANP (6.67%)<br>ACP (4.44%)<br>ATP (4.44%)<br>DAL (4.44%)<br>SO4 (2.22%)<br>N42 (2.22%)<br>2P5 (2.22%)<br>1Q4 (2.22%)<br>Other (17.76%) |
| **DE-DE-R** | 1 | 17 | 1.8 | TruB (5.4.99.25) (3.30.2350.10)<br>limonene-1 (3.3.2.8) (3.10.450.50)<br>mannonate dehydratase (4.2.1.8) (3.20.20.120)<br>xanthine oxidase (1.17.3.2) (3.30.365.10)<br>biotin carboxylase (6.3.4.14) (3.30.470.20)<br>pectinesterase (3.1.1.11) (2.160.20.10)<br>xanthine oxidase (1.17.3.2) (3.30.365.10)<br>Unknown (3.6.1.-) (3.90.79.10)<br>PEPPM (5.4.2.9) (3.20.20.60)<br>squalene---hopene cyclase (5.4.99.17) (1.50.10.20)<br>endo-1 (3.2.1.89) (3.20.20.80)<br>cholesterol oxidase (1.1.3.6) (3.40.462.10)<br>alpha-1 (4.2.2.2) (2.160.20.10) | 21.2 | 0 | 27.2 | 47.1 | 47.1 | 5.9 | MG (21.6%)<br>ZN (0.8%)<br>NA (0.8%) | | URC (9.6%)<br>SAL (6.4%)<br>3ZS (3.2%)<br>CO3 (3.2%)<br>GCO (3.2%)<br>UNX (3.2%)<br>3ZQ (2.4%)<br>ADA (2.4%)<br>HPN (1.6%)<br>HYH (1.6%)<br>Other (28.0%) |

| | | | | | | | | | | | | | |
|---|---|---|---|---|---|---|---|---|---|---|---|---|---|
| | | | | mannonate dehydratase (4.2.1.8) (3.20.20.120)<br>thymidine kinase (2.7.1.21) (3.40.50.300)<br>5 (6.3.3.2) (3.40.50.10420)<br>OSBS (4.2.1.113) (3.20.20.120) | | | | | | | | | |
| **DE-DE-R-STY** | 1 | 3 | 2.2 | swivelase (5.6.2.1) (3.40.50.140)<br>exo-cellobiohydrolase (3.2.1.91) (3.20.20.40)<br>galactarate dehydratase (D-threo-forming) (4.2.1.158) (3.20.20.120) | 37.7 | 0 | 0 | 66.7 | 33.3 | 0 | MG (55.56%)<br>CA (11.11%) | | SSG (11.11%)<br>RCB (11.11%)<br>XYS (11.11%) |
| **DE-DE-STY** | 3 | 15 | 1.3 | dethiobiotin synthase (6.3.3.3) (3.40.50.300)<br>isopentenyl-diphosphate Delta-isomerase (5.3.3.2) (3.90.79.10)<br>EAS (4.2.3.61) (1.10.600.10)<br>6-phosphofructokinase (2.7.1.11) (3.40.50.450)<br>hyaluronoglucosaminidase (3.2.1.35) (3.20.20.70)<br>exodeoxyribonuclease III (3.1.11.2) (3.60.10.10)<br>endothiapepsin (3.4.23.22) (2.40.70.10)<br>steroid Delta-isomerase (5.3.3.1) (3.10.450.50)<br>anhydrosialidase (4.2.2.15) (2.120.10.10)<br>exo-alpha-sialidase (3.2.1.18) (2.120.10.10)<br>P450(eryF) (1.14.15.35) (1.10.630.10)<br>exo-alpha-sialidase (3.2.1.18) (2.120.10.10)<br>formyltetrahydrofolate dehydrogenase (1.5.1.6) (3.40.605.10)<br>tripeptidase (3.4.11.4) (1.10.390.10)<br>chitosanase (3.2.1.132) (1.20.141.10) | 10.1 | 5.1 | 28.9 | 33.3 | 66.7 | 0 | MG (3.89%)<br>MN (1.8%) | NDP (0.9%)<br>NAD (0.6%)<br>HEM (0.3%)<br>NAP (0.3%) | DAN (5.99%)<br>G39 (5.09%)<br>SIA (2.99%)<br>ZMR (2.69%)<br>BES (2.69%)<br>ADP (1.5%)<br>DPO (1.5%)<br>EQP (1.5%)<br>FSI (1.2%)<br>BCZ (1.2%)<br>Other (54.9%) |
| **DE-DE-STY-STY** | 1 | 4 | 1.8 | phosphomonoesterase (3.1.3.1) (3.40.720.10)<br>dTDP-glucose 4 (4.2.1.46) (3.40.50.720)<br>peroxidase (1.11.1.7) (1.10.640.10)<br>endo-1 (3.2.1.8) (2.60.120.180) | 50.5 | 9 | 16 | 50 | 50 | 0 | CA (59.21%)<br>MG (7.24%)<br>ZN (2.63%)<br>CO (0.66%) | NAD (3.29%) | DOD (8.55%)<br>XYP (2.63%)<br>X2F (1.97%)<br>TDX (1.32%)<br>GDD (1.32%)<br>DAU (1.32%)<br>DFX (0.66%)<br>C5X (0.66%)<br>07E (0.66%)<br>C3X (0.66%)<br>Other (0.66%) |
| **DE-DE-X** | 1 | 10 | 1.2 | methionine adenosyltransferase (2.5.1.6) (3.30.300.10)<br>Unknown (3.2.-.-) (3.90.245.10)<br>mGDH (1.1.5.2) (2.120.10.30)<br>phosphonatase (3.11.1.1) (3.40.50.1000)<br>fructose-bisphosphatase (3.1.3.11) (3.30.540.10)<br>acetolactate synthase (2.2.1.6) (3.40.50.970)<br>phosphoserine phosphatase (3.1.3.3) (3.40.50.1000)<br>Unknown (3.1.3.-) (3.40.50.1000)<br>type II site-specific deoxyribonuclease (3.1.21.4) (3.40.600.20)<br>aspartate---ammonia ligase (6.3.1.1) (3.30.930.10) | 86.7 | 14.3 | 45.4 | 100 | 0 | 0 | MG (34.02%)<br>CA (9.28%)<br>TL (7.73%)<br>ZN (4.64%)<br>MN (2.06%)<br>CL (1.55%)<br>K (1.03%)<br>NA (0.52%) | SAM (0.52%)<br>EN0 (0.52%)<br>5GY (0.52%)<br>TPP (0.52%) | PO4 (13.4%)<br>HE3 (1.55%)<br>2O2 (1.55%)<br>PIR (1.03%)<br>DNB (1.03%)<br>PI (1.03%)<br>PO3 (1.03%)<br>ALF (1.03%)<br>APO (1.03%)<br>PPK (0.52%)<br>Other (7.8%) |
| **DE-DE-X-X** | 1 | 2 | 2.4 | vesicle-fusing ATPase (3.6.4.6) (3.40.50.300)<br>nicotinamidase (3.5.1.19) (3.40.50.850) | 4 | 0 | 42.5 | 100 | 0 | 0 | MG (13.33%) | | ATP (40.0%)<br>ADP (13.33%)<br>NIO (13.33%)<br>ANP (6.67%)<br>VGL (6.67%)<br>CAF (6.67%) |
| **DE-F-K-STY** | 1 | 2 | 1.6 | kynureninase (3.7.1.3) (3.40.640.10)<br>L-lysine 6-transaminase (2.6.1.36) (3.40.640.10) | 0 | 83 | 1 | 0 | 100 | 0 | | PLP (71.11%)<br>PMP (2.22%) | RW2 (6.67%)<br>PLP (4.44%)<br>PFM (2.22%)<br>P00 (2.22%)<br>PXG (2.22%)<br>IF1 (2.22%)<br>PSZ (2.22%)<br>POI (2.22%)<br>PPE (2.22%) |
| **DE-F-VLI** | 1 | 3 | 1.9 | DHFR (1.5.1.3) (3.40.430.10)<br>pyruvate oxidase (1.2.3.3) (3.40.50.970)<br>histone acetyltransferase (2.3.1.48) (3.40.630.30) | 0 | 0 | 17 | 0 | 33.3 | 66.7 | | | TRR (40.14%)<br>MTX (16.9%)<br>TMQ (15.49%)<br>BDM (13.38%)<br>MMV (2.11%)<br>QKJ (2.11%)<br>FOL (1.41%)<br>TOP (1.41%)<br>WPD (1.41%)<br>WFA (0.7%)<br>Other (4.9%) |

| | | | | | | | | | | | | | |
|---|---|---|---|---|---|---|---|---|---|---|---|---|---|
| **DE-H-H** | 3 | 21 | 1.5 | beta-lactamase (3.5.2.6) (3.60.15.10)<br>alpha-toxin (3.1.4.3) (1.10.575.10)<br>phosphomonoesterase (3.1.3.2) (3.60.21.10)<br>L-fuculose-phosphate aldolase (4.1.2.17) (3.40.225.10)<br>luciferase (1.14.14.3) (3.20.20.30)<br>phenylalaninase (1.14.16.1) (1.10.800.10)<br>isopenicillin-N synthase (1.21.3.1) (2.60.120.330)<br>biotin carboxylase (6.3.4.14) (3.20.20.70)<br>mutarotase (5.1.3.3) (2.70.98.10)<br>[histone H3]-trimethyl-L-lysine36 demethylase (1.14.11.69) (2.60.120.650)<br>single-stranded-nucleate endonuclease (3.1.30.1) (1.10.575.10)<br>single-stranded-nucleate endonuclease (3.1.30.1) (1.10.575.10)<br>rhamnulose-1-phosphate aldolase (4.1.2.19) (3.40.225.10)<br>leucocyanidin (1.14.20.4) (2.60.120.330)<br>BphI (4.1.3.43) (3.20.20.70)<br>G6PDH (1.1.1.363) (3.30.360.10)<br>4a-hydroxytetrahydrobiopterin dehydratase (4.2.1.96) (3.30.1360.20)<br>methylglyoxal synthase (4.2.3.3) (3.40.50.1380)<br>phospholipase D (3.1.4.4) (3.30.870.10)<br>NDO (1.14.12.12) (3.90.380.10)<br>methylglyoxalase (4.4.1.5) (3.10.180.10) | 61 | 1 | 31 | 71.4 | 28.6 | 0 | ZN (41.22%)<br>FE (6.62%)<br>FE2 (3.89%)<br>MN (1.47%)<br>CO (1.16%)<br>CD (0.95%)<br>NI (0.84%)<br>CL (0.21%)<br>AU (0.21%)<br>MG (0.21%)<br>Other (0.11%) | HBI (0.74%)<br>H4B (0.42%)<br>H2B (0.11%) | MRI (5.89%)<br>RTD (2.1%)<br>OGA (1.79%)<br>OH (0.84%)<br>PO4 (0.84%)<br>AKG (0.63%)<br>ZZ7 (0.53%)<br>ACV (0.53%)<br>QUE (0.42%)<br>MYC (0.42%)<br>Other (21.48%) |
| **DE-H-H-H** | 3 | 21 | 1.7 | ToyB (4.1.2.50) (3.30.479.10)<br>anhydrase (4.2.1.1) (3.10.200.10)<br>glycerol dehydrogenase (1.1.1.6) (1.20.1090.10)<br>astacin (3.4.24.21) (3.40.390.10)<br>metalloelastase (3.4.24.65) (3.40.390.10)<br>glycoprotein gp63 (3.4.24.36) (3.10.170.20)<br>metalloelastase (3.4.24.65) (3.40.390.10)<br>rhamnulose-1-phosphate aldolase (4.1.2.19) (3.40.225.10)<br>oxalate oxidase (1.2.3.4) (2.60.120.10)<br>urease (3.5.1.5) (3.20.20.140)<br>beta-lactamase (3.5.2.6) (3.20.20.140)<br>Unknown (3.1.5.-) (1.10.3210.10)<br>2 (1.13.11.2) (3.10.180.10)<br>hydroxyacylglutathione hydrolase (3.1.2.6) (3.60.15.10)<br>homogentisicase (1.13.11.5) (2.60.120.10)<br>DD-carboxypeptidase (3.4.17.14) (3.30.1380.10)<br>protocatechuate 4 (1.13.11.8) (3.40.830.10)<br>beta-lactamase (3.5.2.6) (3.60.15.10)<br>SOD (1.15.1.1) (2.60.40.200)<br>adenosine deaminase (3.5.4.4) (3.20.20.140)<br>quercetinase (1.13.11.24) (2.60.120.10) | 79.7 | 2.2 | 20.9 | 90.5 | 4.8 | 0 | ZN (55.76%)<br>FE (3.15%)<br>CO (1.54%)<br>FE2 (1.35%)<br>CU (0.71%)<br>MN (0.71%)<br>NI (0.26%)<br>CD (0.26%)<br>MG (0.06%)<br>HG (0.06%)<br>Other (0.12%) | BIO (0.06%)<br>NAD (0.06%)<br>GBP (0.06%) | CGS (2.06%)<br>HSI (1.8%)<br>0DS (1.67%)<br>HAV (1.29%)<br>RTD (1.29%)<br>*P* (0.9%)<br>SO4 (0.77%)<br>R47 (0.45%)<br>NGH (0.39%)<br>GDE (0.39%)<br>Other (21.81%) |
| **DE-H-H-H-H-Z** | 1 | 2 | 0.6 | urease (3.5.1.5) (3.20.20.140)<br>Unknown (3.4.19.-) (3.20.20.140) | 88 | 0 | 11 | 100 | 0 | 0 | NI (41.27%)<br>ZN (11.64%)<br>F (2.65%)<br>MN (1.06%) | | KCX (25.93%)<br>OH (3.17%)<br>PO4 (2.65%)<br>2PA (2.12%)<br>HAE (1.59%)<br>BME (1.06%)<br>BO3 (0.53%)<br>DJM (0.53%)<br>URE (0.53%)<br>9XN (0.53%)<br>Other (4.24%) |
| **DE-H-H-H-NQ** | 1 | 2 | 1.9 | aldolase (4.1.2.13) (3.20.20.70)<br>oxalate oxidase (1.2.3.4) (2.60.120.10) | 100 | 0 | 100 | 100 | 0 | 0 | ZN (37.5%)<br>MN (12.5%) | | PGH (37.5%)<br>GLV (12.5%) |
| **DE-H-H-H-NQ-R** | 1 | 2 | 0.2 | calcineurin (3.1.3.16) (3.60.21.10)<br>calcineurin (3.1.3.16) (3.60.21.10) | 88 | 0 | 37 | 100 | 0 | 0 | MN (45.04%)<br>ZN (12.98%)<br>FE (9.16%)<br>MG (2.29%)<br>NI (2.29%)<br>CL (0.76%) | | PO4 (12.98%)<br>ENL (2.29%)<br>NHC (2.29%)<br>MES (1.53%)<br>SO4 (1.53%)<br>MLI (1.53%)<br>AGS (0.76%)<br>8D4 (0.76%)<br>NHE (0.76%)<br>4TF (0.76%)<br>Other (2.28%) |

| | | | | | | | | | | | | | |
|---|---|---|---|---|---|---|---|---|---|---|---|---|---|
| **DE-H-H-H-STY** | 1 | 2 | 1.9 | 2 (1.13.11.2) (3.10.180.10)<br>thermolysin (3.4.24.27) (3.10.170.10) | 93.5 | 0 | 31 | 100 | 0 | 0 | ZN (47.89%)<br>FE2 (6.1%)<br>FE (4.23%)<br>MN (3.29%)<br>CD (1.41%)<br>CO (0.94%)<br>CU (0.47%)<br>CU1 (0.47%)<br>CL (0.47%)<br>RU (0.47%)<br>Other (0%) | | VAL (1.88%)<br>DHY (1.41%)<br>LEU (1.41%)<br>NX6 (1.41%)<br>ASP (1.41%)<br>ACN (0.94%)<br>M3P (0.94%)<br>FEL (0.94%)<br>MCT (0.94%)<br>*P* (0.94%)<br>Other (19.27%) |
| **DE-H-H-NQ** | 1 | 2 | 2.3 | cytosine deaminase (3.5.4.1) (3.20.20.140)<br>aryldialkylphosphatase (3.1.8.1) (2.120.10.30) | 10 | 0 | 30 | 100 | 0 | 0 | MG (25.0%) | | FPY (25.0%)<br>O7U (16.67%)<br>1LD (8.33%)<br>IGA (8.33%)<br>HPY (8.33%)<br>17E (8.33%) |
| **DE-H-H-R** | 1 | 8 | 1.9 | phosphatidylinositol diacylglycerol-lyase (4.6.1.13) (3.20.20.190)<br>taurine dioxygenase (1.14.11.17) (3.60.130.10)<br>carboxypeptidase T (3.4.17.18) (3.40.630.10)<br>single-stranded-nucleate endonuclease (3.1.30.1) (1.10.575.10)<br>phosphomonoesterase (3.1.3.2) (3.40.50.1240)<br>oxalate decarboxylase (4.1.1.2) (2.60.120.10)<br>VanX (3.4.13.22) (3.30.1380.10)<br>bisphosphoglyceromutase (5.4.2.4) (3.40.50.1240) | 55.2 | 0.2 | 24.1 | 62.5 | 37.5 | 0 | ZN (33.65%)<br>FE2 (3.77%)<br>MN (2.83%)<br>FE (2.52%)<br>CO (2.2%)<br>MG (0.31%)<br>HG (0.31%) | DG2 (0.31%) | AKG (5.97%)<br>*P* (3.77%)<br>INS (2.52%)<br>3PG (2.2%)<br>VVO (1.26%)<br>CXA (1.26%)<br>SO4 (1.26%)<br>3K0 (0.94%)<br>VO4 (0.94%)<br>SIN (0.63%)<br>Other (23.96%) |
| **DE-H-H-STY** | 2 | 4 | 1.5 | Unknown (1.-.-.-) (3.60.15.10)<br>beta-hydroxyoctanoyl-ACP-dehydrase (4.2.1.59) (3.10.129.110)<br>[histone H3]-trimethyl-L-lysine36 demethylase (1.14.11.69) (2.60.120.650)<br>BphI (4.1.3.43) (3.20.20.70) | 30.8 | 0 | 42.8 | 75 | 25 | 0 | FE (3.64%)<br>MN (2.73%)<br>MG (1.82%)<br>ZN (0.91%) | | OGA (15.45%)<br>FEO (6.36%)<br>OXY (4.55%)<br>AKG (2.73%)<br>M3L (2.73%)<br>PYR (2.73%)<br>OXL (2.73%)<br>2HG (1.82%)<br>MMK (1.82%)<br>PD2 (1.82%)<br>Other (41.86%) |
| **DE-H-K** | 2 | 19 | 1.5 | methionine adenosyltransferase (2.5.1.6) (3.30.300.10)<br>PEPK (4.1.1.49) (3.90.228.20)<br>UlaD (4.1.1.85) (3.20.20.70)<br>Unknown (2.4.1.-) (3.90.550.10)<br>allyl-alcohol dehydrogenase (1.1.1.54) (3.20.20.100)<br>isochorismatase (3.3.2.1) (3.40.50.850)<br>phosphoenolpyruvate carboxykinase (GTP) (4.1.1.32) (3.40.449.10)<br>phosphomonoesterase (3.1.3.1) (3.40.720.10)<br>mandelate racemase (5.1.2.2) (3.20.20.120)<br>methylmalonyl-CoA mutase (5.4.99.2) (3.40.50.280)<br>SpeA (4.1.1.19) (3.20.20.10)<br>DHQase (4.2.1.10) (3.20.20.70)<br>(S)-oxynitrilase (4.1.2.47) (3.40.50.1820)<br>xylose isomerase (5.3.1.5) (3.20.20.150)<br>L-rhamnose isomerase (5.3.1.14) (3.20.20.150)<br>kasA (gene name) (2.3.1.293) (3.40.47.10)<br>SpeC (4.1.1.17) (3.40.640.10)<br>gliostatins (2.4.2.4) (3.40.1030.10)<br>neelaredoxin (1.15.1.2) (2.60.40.730) | 36.9 | 11.4 | 30.2 | 47.4 | 47.4 | 0 | MN (21.67%)<br>MG (11.74%)<br>ZN (3.16%)<br>CA (2.26%)<br>CO (1.81%)<br>FE (1.35%)<br>CL (0.23%)<br>NA (0.23%) | PLP (3.84%)<br>NAP (0.23%) | GTP (4.29%)<br>XYL (2.93%)<br>PO4 (2.71%)<br>GLC (2.48%)<br>ATP (2.03%)<br>OXL (2.03%)<br>PEP (1.81%)<br>SPV (1.81%)<br>EDO (1.35%)<br>PGA (1.13%)<br>Other (22.96%) |
| **DE-H-K-NQ** | 1 | 2 | 1.8 | triose-phosphate isomerase (5.3.1.1) (3.20.20.70)<br>phosphohexomutase (5.3.1.8) (2.60.120.10) | 50 | 0 | 11.5 | 50 | 50 | 0 | ZN (6.67%) | | PGA (26.67%)<br>PO4 (20.0%)<br>EDO (13.33%)<br>TPS (6.67%)<br>13P (6.67%)<br>PGH (6.67%)<br>129 (6.67%) |
| **DE-H-NQ** | 1 | 14 | 1.7 | steroid Delta-isomerase (5.3.3.1) (3.50.50.60)<br>6-phosphogluconic dehydrogenase (1.1.1.44) (1.10.1040.10)<br>Unknown (3.1.5.-) (1.10.3210.10)<br>barley nuclease (3.1.30.2) (3.40.570.10)<br>glucarate dehydratase (4.2.1.40) (3.20.20.120)<br>cytosine deaminase (3.5.4.1) (3.20.20.140)<br>peroxidase (1.11.1.7) (1.10.640.10)<br>peroxidase (1.11.1.7) (1.10.640.10) | 8.6 | 18.8 | 27.1 | 42.9 | 57.1 | 0 | MG (2.69%)<br>K (2.39%)<br>IOD (0.6%)<br>NA (0.6%) | HEM (51.34%)<br>FAD (2.39%)<br>FDA (0.3%)<br>FAE (0.3%)<br>SFD (0.3%)<br>NDP (0.3%) | OSM (3.28%)<br>PEO (2.99%)<br>CYN (2.39%)<br>6PG (1.79%)<br>SHA (1.79%)<br>HEC (1.19%)<br>3TR (1.19%)<br>PZA (1.19%) |

| | | | | | | | | | | | | | |
|---|---|---|---|---|---|---|---|---|---|---|---|---|---|
| | | | | oxalate oxidase (1.2.3.4) (2.60.120.10)<br>DHQase (4.2.1.10) (3.40.50.9100)<br>Unknown (2.3.1.-) (3.40.50.1220)<br>alpha-galactosyltransferase (2.4.1.87) (3.90.550.10)<br>glycine amidinotransferase (2.1.4.1) (3.75.10.10)<br>aminoacyl-tRNA hydrolase (3.1.1.29) (3.40.50.1470) | | | | | | | | | MMZ (1.19%)<br>OXY (0.9%)<br>Other (16.2%) |
| DE-H-NQ-STY-X | 1 | 4 | 1.1 | alkylacetyl-GPC:acetylhydrolase (3.1.1.47) (3.40.50.1110)<br>CA (3.5.1.93) (3.60.20.10)<br>RGAE (3.1.1.86) (3.40.50.1110)<br>palmitoyl-CoA hydrolase (3.1.2.2) (3.40.50.1110) | 0 | 0 | 0 | 0 | 100 | 0 | | | GD7 (100.0%) |
| DE-H-NQ-X | 1 | 2 | 1 | adenain (3.4.22.39) (3.40.395.10)<br>nucleoside-triphosphatase (3.6.1.15) (3.90.70.10) | 0 | 0 | 0 | 0 | 100 | 0 | | | E69 (100.0%) |
| DE-H-R | 2 | 17 | 1.5 | FRD (1.3.5.4) (3.90.700.10)<br>glycerophosphodiester phosphodiesterase (3.1.4.46) (3.20.20.190)<br>Xaa-Pro aminopeptidase (3.4.11.9) (3.90.230.10)<br>GS (6.3.1.2) (3.30.590.10)<br>Unknown (3.1.-.-) (3.90.540.10)<br>fructose-2 (3.1.3.46) (3.40.50.1240)<br>chloramphenicol O-acetyltransferase (2.3.1.28) (3.30.559.10)<br>mGDH (1.1.5.2) (2.120.10.30)<br>glycerophosphodiester phosphodiesterase (3.1.4.46) (3.20.20.190)<br>bromoperoxidase (1.11.1.18) (1.10.606.10)<br>bromoperoxidase (1.11.1.18) (1.10.606.10)<br>citrogenase (2.3.3.16) (1.10.230.10)<br>isocitrase (4.1.3.1) (3.20.20.60)<br>L(+)-nLDH (1.1.1.27) (3.90.110.10)<br>bontoxilysin (3.4.24.69) (3.90.1240.10)<br>barnase (4.6.1.24) (3.10.450.30)<br>PdxJ (2.6.99.2) (3.20.20.70) | 10.9 | 3.2 | 30.1 | 47.1 | 52.9 | 0 | MG (5.15%)<br>ZN (5.15%)<br>CA (1.47%)<br>MN (0.37%)<br>FE (0.37%) | NAI (2.21%)<br>NAD (2.21%)<br>CMC (1.47%)<br>CMX (0.74%)<br>ACO (0.74%)<br>HAX (0.37%)<br>SDX (0.37%)<br>FCX (0.37%)<br>COA (0.37%)<br>AMX (0.37%)<br>Other (2.59%) | *P* (8.09%)<br>OAA (5.51%)<br>P3S (2.94%)<br>PO4 (2.94%)<br>CIT (2.94%)<br>PRO (2.21%)<br>GOL (2.21%)<br>OXM (1.84%)<br>FUM (1.47%)<br>ADP (1.47%)<br>Other (32.13%) |
| DE-H-R-R | 1 | 4 | 1.4 | barnase (4.6.1.24) (3.10.450.30)<br>carboxy-cis (5.5.1.5) (2.130.10.10)<br>endo-alpha-sialidase (3.2.1.129) (2.120.10.10)<br>GS (6.3.1.2) (3.30.590.10) | 22.5 | 0 | 23 | 25 | 75 | 0 | MG (17.78%)<br>MN (2.22%) | | P3S (17.78%)<br>2GP (8.89%)<br>ADP (8.89%)<br>PO4 (8.89%)<br>3GP (6.67%)<br>SGP (4.44%)<br>*P* (2.22%)<br>ANP (2.22%) |
| DE-H-R-STY | 1 | 2 | 1.3 | Xaa-Pro aminopeptidase (3.4.11.9) (3.90.230.10)<br>hyaluronate lyase (4.2.2.1) (1.50.10.100) | 0 | 0 | 11.5 | 50 | 50 | 0 | | | PRO (26.09%)<br>*P* (17.39%)<br>M44 (8.7%)<br>GCT (8.7%)<br>IPA (4.35%)<br>MRD (4.35%)<br>PVC (4.35%)<br>ASG (4.35%)<br>IDR (4.35%) |
| DE-H-STY | 3 | 25 | 1.4 | 2,4-dienoyl-CoA reductase (1.3.1.34) (3.20.20.70)<br>trimethylamine dehydrogenase (1.5.8.2) (3.20.20.70)<br>Methylenetetrahydrofolate reductase (1.5.1.20) (3.20.20.220)<br>nucleoside-diphosphate kinase (2.7.4.6) (3.30.70.141)<br>destabilase (3.5.1.44) (3.40.50.180)<br>bromoperoxidase (1.11.1.18) (1.10.606.10)<br>3-mercaptopyruvate sulfurtransferase (2.8.1.2) (3.40.250.10)<br>acetylxylan esterase (3.1.1.72) (3.40.50.1820)<br>DHBP synthase (4.1.99.12) (3.90.870.10)<br>Unknown (2.3.1.-) (3.40.50.1820)<br>endoprolylpeptidase (3.4.21.26) (3.40.50.1820)<br>AlaDH (1.4.1.1) (3.40.50.720)<br>BphI (4.1.3.43) (3.20.20.70)<br>peptidyl-Lys metalloendopeptidase (3.4.24.20) (3.40.390.10)<br>citrogenase (2.3.3.16) (1.10.580.10)<br>phosphatidase (3.1.1.4) (1.20.90.10)<br>scytalone dehydratase (4.2.1.94) (3.10.450.50)<br>D-arginine dehydrogenase (1.4.99.6) (3.50.50.60)<br>barnase (4.6.1.24) (3.10.450.30)<br>4HmO (1.1.3.46) (3.20.20.70)<br>ribulose-phosphate 3-epimerase (5.1.3.1) (3.20.20.70)<br>endo-1 (3.2.1.78) (3.20.20.80) | 11.2 | 9.2 | 14.6 | 28 | 60 | 12 | MG (7.6%)<br>ZN (2.28%)<br>MN (1.14%)<br>CA (1.14%)<br>FE2 (1.14%)<br>CD (0.38%)<br>NA (0.38%) | FMN (12.55%)<br>FNS (0.76%)<br>MDE (0.38%)<br>SDX (0.38%)<br>CIC (0.38%)<br>FNR (0.38%) | 2GP (11.41%)<br>FMN (10.65%)<br>OAA (4.94%)<br>3GP (3.8%)<br>CIT (2.66%)<br>SGP (1.9%)<br>LMR (1.52%)<br>MPD (1.52%)<br>2AM (1.52%)<br>PYR (1.14%)<br>Other (22.42%) |

| | | | | | | | | | | | | | |
|---|---|---|---|---|---|---|---|---|---|---|---|---|---|
| | | | | galactarate dehydratase (D-threo-forming) (4.2.1.158) (3.30.390.10)<br>nucleoside-triphosphatase (3.6.1.15) (2.40.10.10)<br>MetH (2.1.1.13) (3.40.50.280) | | | | | | | | | |
| **DE-H-STY-STY-X** | 1 | 2 | 1.4 | Xaa-Pro-dipeptidyl-aminopeptidase (3.4.14.5) (3.40.50.1820)<br>CocE (3.1.1.84) (3.40.50.1820) | 0 | 0 | 1 | 0 | 100 | 0 | | | *P* (100.0%) |
| **DE-H-STY-X** | 2 | 5 | 0.8 | acetylcholinesterase (3.1.1.7) (3.40.50.1820)<br>narbonolide synthase (2.3.1.240) (3.40.50.1820)<br>DBP2 (3.6.4.13) (2.40.10.120)<br>chymotrypsin (3.4.21.1) (2.40.10.10)<br>coagulation factor XIa (3.4.21.27) (2.40.10.10) | 0 | 0 | 3 | 0 | 100 | 0 | | | 2ZF (21.21%)<br>*P* (15.15%)<br>2R9 (4.04%)<br>SV6 (4.04%)<br>1BV (3.03%)<br>0G6 (3.03%)<br>ELT (2.02%)<br>SUE (2.02%)<br>HTB (1.01%)<br>57H (1.01%)<br>Other (43.43%) |
| **DE-H-STY-X-X** | 1 | 11 | 1.3 | deamidase (3.4.16.5) (3.40.50.11320)<br>butyrinase (3.1.1.3) (3.40.50.1820)<br>prolyl aminopeptidase (3.4.11.5) (3.40.50.1820)<br>butyrinase (3.1.1.3) (3.40.50.1820)<br>palmitoyl[protein] hydrolase (3.1.2.22) (3.40.50.1820)<br>Unknown (3.1.2.-) (3.40.50.1820)<br>dipeptidase E (3.4.13.21) (3.40.50.880)<br>cutinase (3.1.1.74) (3.40.50.1820)<br>acetylxylan esterase (3.1.1.72) (3.40.50.1820)<br>ClpP (3.4.21.92) (3.90.226.10)<br>cholinesterase (3.1.1.8) (3.40.50.1820) | 0 | 0 | 0.7 | 0 | 100 | 0 | | | MUP (14.29%)<br>M3D (14.29%)<br>PLC (14.29%)<br>EMM (14.29%)<br>ELT (14.29%)<br>AT3 (14.29%)<br>NTJ (14.29%) |
| **DE-H-W** | 1 | 4 | 1.6 | stearoyl-[acyl-carrier-protein] 9-desaturase (1.14.19.2) (1.10.620.20)<br>L-ascorbate peroxidase (1.11.1.11) (1.10.420.10)<br>ribonuclease T2 (4.6.1.19) (3.90.730.10)<br>FGD1 (1.1.98.2) (3.20.20.30) | 0 | 10.2 | 0 | 50 | 50 | 0 | | HEM (72.6%)<br>ZNH (6.85%)<br>FMI (2.74%)<br>HEB (2.74%)<br>PP9 (1.37%)<br>DDH (1.37%) | DOD (5.48%)<br>522 (1.37%)<br>HEC (1.37%)<br>F42 (1.37%) |
| **DE-H-X** | 1 | 11 | 1.4 | 4-chlorobenzoyl-CoA dehalogenase (3.8.1.7) (3.90.226.10)<br>methylglyoxal synthase (4.2.3.3) (3.40.50.1380)<br>RNase (4.6.1.18) (3.10.130.10)<br>Unknown (4.6.1.-) (3.20.20.190)<br>acyl-[acyl-carrier-protein]---UDP-N-acetylglucosamine O-acyltransferase (2.3.1.129) (2.160.10.10)<br>transglutaminase (2.3.2.13) (3.90.260.10)<br>5-oxoprolyl-peptidase (3.4.19.3) (3.40.630.20)<br>C_overbar_42_ (3.4.21.43) (2.40.10.10)<br>peptidase K (3.4.21.64) (3.40.50.200)<br>CocE (3.1.1.84) (3.40.50.1820)<br>Unknown (3.1.-.-) (3.40.960.10) | 0.2 | 0 | 16.2 | 18.2 | 81.8 | 0 | CL (2.17%) | | *P* (19.57%)<br>DOD (6.52%)<br>PO4 (4.35%)<br>CGP (3.26%)<br>BCA (2.17%)<br>C3P (2.17%)<br>U3P (2.17%)<br>IMP (2.17%)<br>C2P (2.17%)<br>AMP (2.17%)<br>Other (28.3%) |
| **DE-H-X-X** | 2 | 6 | 1.3 | trypsin (3.4.21.4) (2.40.10.10)<br>nucleoside-triphosphatase (3.6.1.15) (2.40.10.10)<br>coagulation factor XIa (3.4.21.27) (2.40.10.10)<br>Lys-gingipain (3.4.22.47) (3.40.50.1460)<br>AtECH2 (4.2.1.119) (3.10.129.10)<br>beta-hydroxyoctanoyl-ACP-dehydrase (4.2.1.59) (3.10.129.110) | 2.3 | 0 | 4.2 | 0 | 100 | 0 | CL (1.06%) | | *P* (24.47%)<br>AG7 (8.51%)<br>0G6 (2.13%)<br>15U (2.13%)<br>DRX (2.13%)<br>5GI (2.13%)<br>G85 (2.13%)<br>G83 (2.13%)<br>K7J (1.06%)<br>BBL (1.06%)<br>Other (49.82%) |
| **DE-H-X-X-X** | 2 | 4 | 0.3 | alpha-lytic endopeptidase (3.4.21.12) (2.40.10.10)<br>alpha-lytic endopeptidase (3.4.21.12) (2.40.10.10)<br>trans-2 (5.3.3.14) (3.10.129.10)<br>beta-hydroxyoctanoyl-ACP-dehydrase (4.2.1.59) (3.10.129.10) | 13.2 | 0.8 | 5.8 | 0 | 100 | 0 | CL (31.37%) | PNS (1.96%) | *P* (13.73%)<br>B2F (7.84%)<br>AES (7.84%)<br>0EG (3.92%)<br>2A1 (3.92%)<br>B2V (3.92%)<br>GOL (1.96%)<br>C9H (1.96%)<br>3MQ (1.96%)<br>7SB (1.96%)<br>Other (9.8%) |

| | | | | | | | | | | | | | |
|---|---|---|---|---|---|---|---|---|---|---|---|---|---|
| **DE-K-K** | 2 | 11 | 1.3 | glutathione synthase (6.3.2.3) (3.30.470.20)<br>glutathione synthase (6.3.2.3) (3.30.470.20)<br>aldolase (4.1.2.13) (3.20.20.70)<br>glucocycloaldolase (5.5.1.4) (3.40.50.720)<br>phosphodeoxyriboaldolase (4.1.2.4) (3.20.20.70)<br>acetylglutamate kinase (2.7.2.8) (3.40.1160.10)<br>isopentenyl phosphate kinase (2.7.4.26) (3.40.1160.10)<br>indole-3-glycerol-phosphate synthase (4.1.1.48) (3.20.20.70)<br>beta-lactamase (3.5.2.6) (3.40.710.10)<br>D-ribulose-1 (4.1.1.39) (3.20.20.110)<br>3'-phosphoadenylyl-sulfate:[heparan sulfate]-glucosamine 3-sulfotransferase (2.8.2.30) (3.40.50.300) | 5.5 | 5.9 | 50.9 | 27.3 | 45.5 | 27.3 | MG (10.44%)<br>ZN (0.55%) | NAD (3.3%)<br>NAI (2.75%) | CAP (24.73%)<br>ADP (8.24%)<br>PO4 (4.4%)<br>RUB (3.3%)<br>SO4 (2.75%)<br>ACY (1.65%)<br>NLG (1.65%)<br>MSE (1.65%)<br>137 (1.65%)<br>13P (1.1%)<br>Other (24.2%) |
| **DE-K-NQ** | 2 | 7 | 1.3 | PEPPM (5.4.2.9) (3.20.20.60)<br>3-phosphoshikimate 1-carboxyvinyltransferase (2.5.1.19) (3.65.10.10)<br>PDHK (2.7.11.2) (3.30.565.10)<br>Dp38 (2.7.11.24) (1.10.510.10)<br>Unknown (5.5.1.-) (3.20.20.120)<br>cyclohydrolase (3.5.4.9) (3.40.50.10860)<br>selenocysteine lyase (4.4.1.16) (3.40.640.10) | 16 | 14.3 | 15 | 57.1 | 28.6 | 14.3 | MG (25.96%)<br>MN (2.88%) | PLP (7.69%) | ANP (9.62%)<br>FMT (5.77%)<br>GPJ (4.81%)<br>ATP (3.85%)<br>*P* (2.88%)<br>L34 (2.88%)<br>OXL (1.92%)<br>PO4 (1.92%)<br>GPF (1.92%)<br>ADP (1.92%)<br>Other (22.08%) |
| **DE-K-R** | 2 | 5 | 1.4 | 3-dehydroquinate-forming) (4.2.3.4) (1.20.1090.10)<br>pentalenene synthase (4.2.3.7) (1.10.600.10)<br>3'-deoxynucleotidase (3.1.3.34) (3.40.50.300)<br>endo-polygalacturonase (3.2.1.15) (2.160.20.10)<br>exodeoxyribonuclease (lambda-induced) (3.1.11.3) (1.10.150.20) | 0 | 0 | 18.6 | 40 | 40 | 20 | | | CRB (45.45%)<br>GTR (18.18%)<br>GLY (9.09%)<br>GLU (9.09%)<br>GTK (9.09%) |
| **DE-K-STY** | 2 | 13 | 1.7 | glutathione-disulfide reductase (1.8.1.7) (3.50.50.60)<br>NAD+-malic enzyme (1.1.1.38) (3.40.50.10380)<br>transaldolase (2.2.1.2) (3.20.20.70)<br>ingensin (3.4.25.1) (3.60.20.10)<br>glutamin-(asparagin-)ase (3.5.1.38) (3.40.50.1170)<br>NADP+-ICDH (1.1.1.42) (3.40.718.10)<br>dethiobiotin synthase (6.3.3.3) (3.40.50.300)<br>diphosphate---fructose-6-phosphate 1-phosphotransferase (2.7.1.90) (3.40.50.450)<br>AvDH1 (3.6.4.12) (3.40.50.300)<br>UDP-N-acetylmuramate---L-alanine ligase (6.3.2.8) (3.40.1190.10)<br>glycine hydroxymethyltransferase (2.1.2.1) (3.40.640.10)<br>glutamate-1-semialdehyde 2 (5.4.3.8) (3.40.640.10)<br>7 (2.6.1.62) (3.40.640.10) | 8.6 | 19.5 | 41.7 | 30.8 | 46.2 | 15.4 | MG (5.12%)<br>CA (1.97%)<br>MN (0.98%) | PLP (23.82%)<br>FAD (12.8%)<br>PMP (2.36%) | PLG (5.51%)<br>PLP (4.92%)<br>IPM (3.54%)<br>FAD (2.76%)<br>ASP (2.76%)<br>ICT (1.97%)<br>ADP (1.57%)<br>F6R (1.38%)<br>PMP (1.38%)<br>04C (1.18%)<br>Other (25.1%) |
| **DE-K-W** | 1 | 2 | 1 | PSAT (2.6.1.52) (3.40.640.10)<br>AAT (2.6.1.1) (3.40.640.10) | 0 | 74 | 0 | 0 | 100 | 0 | | PLP (60.18%)<br>PMP (7.08%)<br>PLR (0.88%)<br>NOP (0.88%)<br>MPL (0.88%) | PLP (5.31%)<br>PLA (5.31%)<br>PPD (1.77%)<br>PGU (1.77%)<br>IK2 (1.77%)<br>PMP (1.77%)<br>PP3 (1.77%)<br>PY6 (0.88%)<br>77E (0.88%)<br>PY5 (0.88%)<br>Other (3.52%) |
| **DE-K-X** | 1 | 3 | 1.8 | 2-chloro-2 (5.5.1.7) (3.20.20.120)<br>asparaginase (3.5.1.1) (3.40.50.1170)<br>exodeoxyribonuclease (lambda-induced) (3.1.11.3) (3.90.320.10) | 26.3 | 0 | 34 | 66.7 | 33.3 | 0 | CA (8.22%)<br>MG (4.11%)<br>CL (1.37%) | | ASP (9.59%)<br>DGL (8.22%)<br>TYR (5.48%)<br>NPQ (4.11%)<br>GLU (2.74%)<br>5CR (2.74%)<br>NLQ (2.74%)<br>MUC (2.74%)<br>AME (2.74%)<br>OSB (2.74%)<br>Other (15.07%) |
| **DE-NQ-NQ-STY** | 1 | 2 | 1.5 | lysozyme (3.2.1.17) (1.10.530.10)<br>meso-2 (4.2.1.28) (3.20.20.350) | 50 | 0 | 38 | 50 | 50 | 0 | K (40.91%)<br>CA (13.64%) | | PGO (31.82%)<br>*P* (4.55%)<br>GOL (4.55%)<br>PGR (4.55%) |

| | | | | | | | | | | | | | |
|---|---|---|---|---|---|---|---|---|---|---|---|---|---|
| **DE-NQ-R** | 1 | 6 | 1.6 | aspartate---ammonia ligase (6.3.1.1) (3.30.930.10)<br>adenylylcyclase (4.6.1.1) (1.10.400.10)<br>acetokinase (2.7.2.1) (3.30.420.40)<br>A-kinase (2.7.11.1) (3.30.565.10)<br>MICL (4.1.3.30) (3.20.20.60)<br>AvDH1 (3.6.4.12) (3.40.50.300) | 10 | 0 | 47.2 | 33.3 | 50 | 16.7 | MG (3.03%) | | GSP (26.26%)<br>GDP (26.26%)<br>ALF (26.26%)<br>ASN (4.04%)<br>ATP (4.04%)<br>ADP (3.03%)<br>AMP (2.02%)<br>B4P (2.02%)<br>SIN (1.01%)<br>APC (1.01%)<br>Other (1.01%) |
| **DE-NQ-STY** | 1 | 3 | 1.4 | L-DEX (3.8.1.2) (3.40.50.1000)<br>beta-glucosidase (3.2.1.21) (3.20.20.80)<br>beta-galactosidase (3.2.1.23) (3.20.20.80) | 0 | 0 | 27.3 | 0 | 100 | 0 | | | BGC (8.05%)<br>BG6 (5.75%)<br>G2F (5.75%)<br>BGP (4.6%)<br>GIM (3.45%)<br>GOX (3.45%)<br>GLC (2.3%)<br>S55 (2.3%)<br>GTL (2.3%)<br>IFM (2.3%)<br>Other (28.75%) |
| **DE-NQ-W** | 1 | 2 | 1.9 | photolyase (4.1.99.3) (1.25.40.80)<br>2-hydroxychromene-2-carboxylate isomerase (5.99.1.4)<br>(3.40.30.10) | 0 | 41 | 3 | 0 | 50 | 50 | | FAD (66.67%) | FAD (14.29%)<br>TCP (4.76%)<br>*P* (4.76%)<br>TDR (4.76%)<br>GSH (4.76%) |
| **DE-NQ-X** | 2 | 10 | 1.4 | formyl-CoA transferase (2.8.3.16) (3.40.50.10540)<br>Delta3-Delta2-enoyl-CoA isomerase (5.3.3.8) (3.90.226.10)<br>glutathione synthase (6.3.2.3) (3.30.1490.50)<br>Delta3-Delta2-enoyl-CoA isomerase (5.3.3.8) (3.90.226.10)<br>indolepyruvate decarboxylase (4.1.1.74) (3.40.50.970)<br>CEAS (2.5.1.66) (3.40.50.970)<br>isoamylase (3.2.1.68) (3.20.20.80)<br>pantothenate synthetase (6.3.2.1) (3.40.50.620)<br>adenylosuccinate synthase (6.3.4.4) (3.40.440.10)<br>ErmC' (2.1.1.184) (3.40.50.150) | 41.1 | 37.4 | 18.7 | 60 | 30 | 10 | MG (23.29%)<br>CA (0.68%) | TPP (13.01%)<br>COA (2.74%)<br>CO8 (1.37%)<br>SAM (1.37%)<br>CAA (0.68%)<br>3HC (0.68%)<br>3H9 (0.68%)<br>GSH (0.68%)<br>TDL (0.68%)<br>TDP (0.68%)<br>Other (2.04%) | GDP (7.53%)<br>IMP (6.16%)<br>IMO (4.11%)<br>SO4 (3.42%)<br>HDA (3.42%)<br>PO4 (2.74%)<br>NO3 (2.74%)<br>AMP (2.05%)<br>ADP (1.37%)<br>ACT (1.37%)<br>Other (12.26%) |
| **DE-R-R** | 3 | 10 | 1.5 | thermonuclease (3.1.31.1) (2.40.50.90)<br>glutathione synthase (6.3.2.3) (3.30.470.20)<br>M.BsuRIa (2.1.1.37) (3.40.50.150)<br>duplicase (2.7.7.7) (2.30.40.20)<br>E1 (6.2.1.45) (3.50.50.80)<br>E1 (6.2.1.45) (3.50.50.80)<br>TruA (5.4.99.12) (3.30.70.660)<br>SerRS (6.1.1.11) (3.30.930.10)<br>7 (4.1.2.25) (3.30.70.560)<br>thymidine kinase (2.7.1.21) (3.40.50.300) | 12.3 | 0 | 56.7 | 50 | 10 | 40 | CA (2.4%)<br>MG (1.6%) | | THP (72.8%)<br>SO4 (7.2%)<br>*P* (4.8%)<br>PO4 (2.4%)<br>C49 (2.4%)<br>ATP (1.6%)<br>C36 (0.8%)<br>C37 (0.8%)<br>PYO (0.8%)<br>DTP (0.8%)<br>Other (1.6%) |
| **DE-R-STY** | 4 | 16 | 1.6 | PTR1 (1.5.1.33) (3.40.50.720)<br>hexokinase (2.7.1.1) (3.30.420.40)<br>hexokinase (2.7.1.1) (3.30.420.40)<br>trans (4.2.3.6) (1.10.600.10)<br>NAD+ glycohydrolase (3.2.2.5) (3.90.176.10)<br>D-camphor-exo-hydroxylase (1.14.15.1) (1.10.630.10)<br>uridine-5'-diphospho-N-acetyl-2-amino-2-deoxy-3-O-<br>lactylglucose:NADP-oxidoreductase (1.3.1.98) (3.90.78.10)<br>3-hydroxybenzoate 4-monooxygenase (1.14.13.23) (3.30.9.10)<br>SerRS (6.1.1.11) (3.30.930.10)<br>gliostatins (2.4.2.4) (3.40.1030.10)<br>biliverdin reductase (1.3.1.24) (3.30.360.10)<br>exo-alpha-sialidase (3.2.1.18) (2.120.10.10)<br>Unknown (2.4.2.-) (3.90.210.10)<br>UGM (5.4.99.9) (3.40.50.720)<br>uroporphyrinogen decarboxylase (4.1.1.37) (3.20.20.210)<br>aspartate---tRNA ligase (6.1.1.12) (3.30.930.10) | 0.7 | 16.2 | 16.7 | 18.8 | 56.2 | 25 | K (0.3%) | NAP (15.41%)<br>FAD (2.72%)<br>NDP (2.42%)<br>HBI (0.6%)<br>H4B (0.3%)<br>TAD (0.3%) | NAP (9.97%)<br>G39 (4.23%)<br>ZMR (2.42%)<br>SSA (2.11%)<br>SIA (2.11%)<br>DAN (1.81%)<br>MTX (1.51%)<br>G6P (1.21%)<br>AMP (1.21%)<br>BCZ (1.21%)<br>Other (46.56%) |
| **DE-R-X** | 1 | 2 | 1.4 | phosphohexomutase (5.3.1.9) (3.40.50.10490)<br>Unknown (6.3.1.21) (3.30.470.20) | 0 | 0 | 9 | 50 | 50 | 0 | | | 6PG (16.67%)<br>F6P (16.67%)<br>PA5 (10.0%)<br>E4P (10.0%)<br>S6P (10.0%)<br>A5P (6.67%)<br>DER (6.67%)<br>RI2 (3.33%) |

| | | | | | | | | | | | | | |
|---|---|---|---|---|---|---|---|---|---|---|---|---|---|
| | | | | | | | | | | | | | G6Q (3.33%)<br>G6P (3.33%)<br>Other (3.33%) |
| **DE-STY-STY** | 3 | 11 | 1.7 | phosphatidase (3.1.1.4) (1.20.90.10)<br>ArsC (ambiguous) (1.20.4.4) (3.40.50.2300)<br>ingensin (3.4.25.1) (3.60.20.10)<br>dTDP-glucose 4 (4.2.1.46) (3.40.50.720)<br>cytochrome P450nor (1.7.1.14) (1.10.630.10)<br>D-alanine---D-alanine ligase (6.3.2.4) (3.30.1490.20)<br>Transferred to 5.6.1.7 (3.6.4.9) (1.10.560.10)<br>PARP (2.4.2.30) (3.90.228.10)<br>indophenolase (7.1.1.9) (1.20.210.10)<br>glutamin-(asparagin-)ase (3.5.1.38) (3.40.50.1170)<br>asparaginase (3.5.1.1) (3.40.50.1170) | 4.7 | 1.6 | 34.2 | 0 | 54.5 | 36.4 | K (0.77%)<br>NA (0.38%)<br>MG (0.38%) | NAD (1.54%) | *P* (28.08%)<br>ASP (8.08%)<br>ADP (4.23%)<br>DAL (2.69%)<br>CNQ (1.54%)<br>2YQ (1.54%)<br>09L (1.54%)<br>78P (1.54%)<br>3BV (1.15%)<br>POL (1.15%)<br>Other (43.09%) |
| **DE-STY-W** | 1 | 2 | 1.3 | thymidylate synthase (2.1.1.45) (3.30.572.10)<br>AlkA (3.2.2.21) (1.10.340.30) | 0 | 2 | 15 | 0 | 50 | 50 | | C2F (7.32%)<br>THG (2.44%) | UMP (41.46%)<br>UFP (19.51%)<br>UMC (4.88%)<br>NOH (4.88%)<br>NDU (2.44%)<br>CB3 (2.44%)<br>VLD (2.44%)<br>FGT (2.44%)<br>CF9 (2.44%) |
| **DE-STY-X** | 1 | 4 | 1.7 | DNA[cytosine-N4]methyltransferase (2.1.1.113) (3.40.50.150)<br>phosphatidase (3.1.1.4) (3.40.1090.10)<br>limonene-1 (3.3.2.8) (3.10.450.50)<br>cis-aconitase (4.2.1.3) (3.20.19.10) | 0 | 0 | 18.2 | 0 | 100 | 0 | | | 3ZS (18.52%)<br>3ZQ (11.11%)<br>HPN (7.41%)<br>HYH (7.41%)<br>6VV (3.7%)<br>VPR (3.7%)<br>40O (3.7%)<br>ATH (3.7%)<br>MIC (3.7%)<br>FLC (3.7%)<br>Other (18.5%) |
| **DE-STY-X-X** | 1 | 2 | 1.8 | 1 (4.1.3.36) (3.90.226.10)<br>arsenite-transporting ATPase (7.3.2.7) (3.40.50.300) | 12.5 | 37.5 | 50 | 0 | 0 | 100 | MG (12.5%) | 1HA (25.0%)<br>2NE (12.5%) | ADP (50.0%) |
| **DE-STY-Z** | 1 | 2 | 0.5 | L-cysteine desulfidase (4.4.1.28) (3.40.640.10)<br>homocysteine desulfhydrase (4.4.1.2) (3.40.640.10) | 0 | 19 | 0 | 0 | 100 | 0 | | PLP (18.8%) | LLP (35.04%)<br>PLP (6.84%)<br>4LM (6.84%)<br>3LM (5.13%)<br>2LM (3.42%)<br>5OW (3.42%)<br>HEN (1.71%)<br>PPJ (1.71%)<br>KOU (1.71%)<br>LPI (1.71%)<br>Other (13.68%) |
| **DE-X-X** | 2 | 9 | 1.4 | beta-lactamase (3.5.2.6) (3.40.710.10)<br>Delta3-Delta2-enoyl-CoA isomerase (5.3.3.8) (3.90.226.10)<br>Transferred to 4.1.2.61 (4.2.1.101) (3.90.226.10)<br>triose-phosphate isomerase (5.3.1.1) (3.20.20.70)<br>Transferred to 5.6.1.3 (3.6.4.4) (3.40.850.10)<br>Unknown (5.3.3.-) (3.90.226.10)<br>crotonase (4.2.1.17) (3.90.226.40)<br>isovaleryl-CoA dehydrogenase (1.3.8.4) (1.20.140.10)<br>ArsC (ambiguous) (1.20.4.4) (3.40.50.2300) | 0.1 | 5.7 | 26.9 | 0 | 66.7 | 22.2 | K (0.84%) | ACO (4.2%)<br>CO8 (1.68%) | NXL (5.04%)<br>DOD (5.04%)<br>PGA (5.04%)<br>SO4 (4.2%)<br>PO4 (3.36%)<br>CB4 (2.52%)<br>ISS (2.52%)<br>2UL (1.68%)<br>C8V (1.68%)<br>IM2 (1.68%)<br>Other (50.4%) |
| **DE-X-X-X** | 1 | 3 | 1.9 | phosphatidase (3.1.1.4) (1.20.90.10)<br>ribokinase (2.7.1.15) (3.40.1190.20)<br>ketohexokinase (2.7.1.3) (3.40.1190.20) | 16.7 | 0 | 21.7 | 33.3 | 66.7 | 0 | CA (28.18%)<br>NA (1.36%)<br>CL (0.91%)<br>CD (0.45%) | | ADN (14.55%)<br>ADP (12.73%)<br>ATP (5.0%)<br>ACP (4.55%)<br>ANP (3.18%)<br>KDG (1.82%)<br>FRU (1.36%)<br>ACY (0.91%)<br>U8D (0.91%) |

| | | | | | | | | | | | | |
|---|---|---|---|---|---|---|---|---|---|---|---|---|
| | | | | | | | | | | | | GEL (0.91%)<br>Other (15.79%) |
| **F-NQ-VLI** | 1 | 2 | 0.9 | pyruvate oxidase (1.2.3.3) (3.40.50.970)<br>pyruvate:ubiquinone-8-oxidoreductase (1.2.5.1) (3.40.50.970) | 0 | 25 | 0 | 0 | 0 | 100 | | TDP (100.0%) | |
| **H-H-H** | 1 | 3 | 1 | HGAD (4.3.2.5) (2.60.120.310)<br>SOD (1.15.1.1) (2.60.40.200)<br>isopentenyl-diphosphate Delta-isomerase (5.3.3.2) (3.90.79.10) | 85.3 | 0 | 0.3 | 100 | 0 | 0 | CU (41.62%)<br>CU1 (27.17%)<br>ZN (10.98%)<br>MN (5.78%)<br>CD (0.58%) | | AZI (1.16%)<br>CO3 (0.58%)<br>PEO (0.58%) |
| **H-H-H-H** | 1 | 3 | 1.9 | phosphomonoesterase (3.1.3.2) (3.60.21.10)<br>laccase (1.10.3.2) (2.60.40.420)<br>laccase (1.10.3.2) (2.60.40.420) | 69 | 0 | 50 | 100 | 0 | 0 | CU (82.63%)<br>NA (0.28%)<br>F (0.28%) | | OXY (10.36%)<br>PER (2.8%)<br>SO4 (0.84%)<br>PO4 (0.28%)<br>0K7 (0.28%)<br>WO4 (0.28%)<br>AD9 (0.28%)<br>0LV (0.28%)<br>H1T (0.28%)<br>ANP (0.28%)<br>Other (0%) |
| **H-H-H-NQ** | 1 | 4 | 1.6 | 3-dehydroquinate-forming) (4.2.3.4) (1.20.1090.10)<br>aldolase (4.1.2.13) (3.20.20.70)<br>anhydrase (4.2.1.1) (2.160.10.10)<br>13-lipoxidase (1.13.11.12) (1.20.245.10) | 78.5 | 0 | 59.5 | 75 | 0 | 25 | FE2 (29.79%)<br>FE (24.47%)<br>ZN (13.83%)<br>CO (3.19%)<br>MN (3.19%) | | CRB (5.32%)<br>BCT (4.26%)<br>PGH (3.19%)<br>SO4 (3.19%)<br>PO4 (1.06%)<br>GLU (1.06%)<br>9OH (1.06%)<br>13S (1.06%)<br>13R (1.06%)<br>EGT (1.06%)<br>Other (1.06%) |
| **H-H-H-STY** | 1 | 3 | 1.7 | indophenolase (7.1.1.9) (1.20.210.10)<br>beta-lactamase (3.5.2.6) (3.60.15.10)<br>primary-amine oxidase (1.4.3.21) (2.70.98.20) | 92.3 | 31.3 | 41.7 | 100 | 0 | 0 | CU (38.23%)<br>ZN (8.03%)<br>CU1 (1.39%)<br>CO (1.39%)<br>NI (0.83%)<br>O (0.28%)<br>CD (0.28%)<br>CL (0.28%) | HEA (16.9%) | TYQ (6.65%)<br>AZI (3.6%)<br>TPQ (3.6%)<br>PER (3.32%)<br>CMO (2.49%)<br>PEO (1.66%)<br>OH (1.11%)<br>HEA (1.11%)<br>MX1 (1.11%)<br>NO (0.83%)<br>Other (5.59%) |
| **H-H-K** | 1 | 2 | 0.7 | RNase (4.6.1.18) (3.10.130.10)<br>Unknown (3.1.27.-) (3.10.130.10) | 3 | 1 | 12.5 | 0 | 100 | 0 | CL (2.59%)<br>ZN (0.86%)<br>PT (0.86%)<br>AU (0.86%) | NDP (0.86%)<br>NAP (0.86%) | *P* (4.31%)<br>U2G (2.59%)<br>ADT (2.59%)<br>U5P (2.59%)<br>AMP (2.59%)<br>POP (1.72%)<br>ATR (1.72%)<br>SFB (1.72%)<br>U5F (1.72%)<br>UMF (1.72%)<br>Other (27.52%) |
| **H-H-NQ** | 2 | 5 | 1.3 | bis(5'-adenosyl)-triphosphatase (3.6.1.29) (3.30.428.10)<br>UDP-sugar diphosphatase (3.6.1.45) (3.60.21.10)<br>aryldialkylphosphatase (3.1.8.1) (2.120.10.30)<br>beta-lactamase (3.5.2.6) (3.60.15.10)<br>Transferred to 2.7.1.191 and 2.7.1.192 and 2.7.1.193 and 2.7.1.194 and 2.7.1.195 and 2.7.1.196 and 2.7.1.197 and 2.7.1.198 and 2.7.1.199 and 2.7.1.200 and 2.7.1.201 and 2.7.1.202 and 2.7.1.203 and 2.7.1.204 and 2.7.1.205 and 2.7.1.206 and 2.7.1.207 and 2.7.1.208 (2.7.1.69) (1.20.58.80) | 24.2 | 0 | 12.4 | 80 | 20 | 0 | ZN (37.02%)<br>CD (1.1%)<br>CL (0.55%) | | RTD (10.5%)<br>SO4 (2.21%)<br>ZZ7 (2.21%)<br>AMP (1.1%)<br>PO4 (1.1%)<br>CO3 (1.1%)<br>BCT (1.1%)<br>QT2 (1.1%)<br>X8Z (1.1%)<br>S3C (1.1%)<br>Other (28.05%) |

| | | | | | | | | | | | | | |
|---|---|---|---|---|---|---|---|---|---|---|---|---|---|
| **H-H-R** | 2 | 7 | 1.4 | urease (3.5.1.5) (3.20.20.140)<br>phosphomonoesterase (3.1.3.2) (1.20.144.10)<br>NadB (1.4.3.16) (3.50.50.60)<br>SOD (1.15.1.1) (2.60.40.200)<br>Unknown (3.1.-.-) (3.90.540.10)<br>L-ascorbate peroxidase (1.11.1.11) (1.10.420.10)<br>UDP-N-acetyl-D-glucosamine 2-epimerase (5.1.3.14) (3.40.50.2000) | 15.3 | 13 | 27.9 | 57.1 | 42.9 | 0 | CU (15.64%)<br>CU1 (13.13%)<br>ZN (4.47%)<br>NI (3.35%)<br>O (0.56%)<br>MN (0.28%)<br>NA (0.28%)<br>F (0.28%) | HEM (28.49%)<br>ZNH (1.12%)<br>HEB (0.84%)<br>FMI (0.56%)<br>PP9 (0.28%)<br>DDH (0.28%) | TOX (3.91%)<br>SO4 (2.79%)<br>DOD (1.96%)<br>OXY (1.68%)<br>HEM (1.4%)<br>2PA (1.12%)<br>HAE (0.84%)<br>PO4 (0.84%)<br>CO3 (0.56%)<br>MOO (0.56%)<br>Other (5.88%) |
| **H-H-STY** | 1 | 11 | 1.4 | NO-forming) (1.7.2.1) (2.60.40.420)<br>assemblin (3.4.21.97) (3.20.16.10)<br>galactose oxidase (1.1.3.9) (2.130.10.80)<br>astacin (3.4.24.21) (3.40.390.10)<br>OYE (1.6.99.1) (3.20.20.70)<br>nicotinamidase (3.5.1.19) (3.40.50.850)<br>1 (1.13.11.1) (2.60.130.10)<br>redoxyendonuclease (3.1.21.2) (3.20.20.150)<br>tyrosine---tRNA ligase (6.1.1.1) (3.40.50.620)<br>ATP-sulfurylase (2.7.7.4) (3.40.50.620)<br>EIIGlc (2.7.1.199) (2.70.70.10) | 40.9 | 3.4 | 32.7 | 54.5 | 36.4 | 9.1 | FE (11.34%)<br>ZN (7.14%)<br>CU (6.72%)<br>HG (0.42%)<br>CO (0.42%)<br>NI (0.42%) | FMN (13.03%)<br>FNR (0.42%) | FMN (5.04%)<br>SO4 (5.04%)<br>*P* (3.36%)<br>ACT (2.94%)<br>CAQ (2.1%)<br>FNR (1.68%)<br>COU (1.68%)<br>0FP (1.26%)<br>HBA (1.26%)<br>DHB (1.26%)<br>Other (26.46%) |
| **H-H-Z** | 1 | 2 | 1.4 | urease (3.5.1.5) (3.20.20.140)<br>aryldialkylphosphatase (3.1.8.1) (3.20.20.140) | 95.5 | 0 | 24 | 100 | 0 | 0 | NI (35.14%)<br>ZN (9.46%)<br>CO (3.15%)<br>F (2.25%)<br>CD (2.25%)<br>MN (1.8%) | | KCX (23.87%)<br>FMT (3.15%)<br>OH (2.7%)<br>PO4 (2.25%)<br>2PA (1.8%)<br>HAE (1.35%)<br>BME (0.9%)<br>BO3 (0.45%)<br>SO4 (0.45%)<br>DJM (0.45%)<br>Other (4.05%) |
| **H-K-NQ** | 1 | 2 | 1.8 | nucleoside-diphosphate kinase (2.7.4.6) (3.30.70.141)<br>SPT (2.3.1.50) (3.40.640.10) | 6.5 | 18.5 | 28 | 0 | 100 | 0 | MG (4.65%) | PLP (8.14%) | ADP (15.12%)<br>CDP (5.81%)<br>UDP (4.65%)<br>TYD (4.65%)<br>GDP (4.65%)<br>PLP (4.65%)<br>DGI (2.33%)<br>AMP (2.33%)<br>PCG (2.33%)<br>3AN (2.33%)<br>Other (22.04%) |
| **H-K-STY** | 1 | 10 | 1.3 | gliostatins (2.4.2.4) (3.40.1030.10)<br>TDP-4-keto-L-rhamnose-3 (5.1.3.13) (2.60.120.10)<br>tRNA-intron lyase (4.6.1.16) (3.40.1350.10)<br>dehydrogenase (1.1.2.3) (3.20.20.70)<br>uricase (1.7.3.3) (3.10.270.10)<br>DD-peptidase (3.4.16.4) (3.40.710.10)<br>allyl-alcohol dehydrogenase (1.1.1.54) (3.20.20.100)<br>6-phosphogluconic dehydrogenase (1.1.1.44) (3.40.50.720)<br>cerebroside-sulfatase (3.1.6.8) (3.40.720.10)<br>UDP-sulfoquinovose synthase (3.13.1.1) (3.40.50.720) | 1.6 | 16.8 | 28.9 | 0 | 100 | 0 | CL (3.31%)<br>K (0.25%) | NAP (16.28%)<br>FMN (7.63%)<br>NDP (2.54%)<br>FNS (0.51%)<br>NAD (0.51%)<br>FNR (0.25%) | AZA (11.7%)<br>DOD (3.31%)<br>NAP (2.54%)<br>OXY (2.29%)<br>URC (2.04%)<br>ZST (1.78%)<br>LDT (1.78%)<br>FID (1.53%)<br>6PG (1.53%)<br>AZI (1.27%)<br>Other (32.51%) |
| **H-K-STY-STY** | 1 | 2 | 2.5 | dehydrogenase (1.1.2.3) (3.20.20.70)<br>UDP-sulfoquinovose synthase (3.13.1.1) (3.40.50.720) | 0 | 43.5 | 50 | 0 | 100 | 0 | | FMN (73.17%)<br>FNS (4.88%)<br>FNR (2.44%) | FMN (9.76%)<br>UPG (4.88%) |
| **H-K-X** | 1 | 2 | 1.5 | adenylosuccinate synthase (6.3.4.4) (3.40.440.10)<br>RNase (4.6.1.18) (3.10.130.10) | 38 | 0 | 58 | 50 | 50 | 0 | MG (11.98%)<br>CL (5.39%)<br>NA (0.6%)<br>IR (0.6%) | | GDP (11.98%)<br>DOD (7.19%)<br>IMP (4.79%)<br>PO4 (4.19%)<br>IMO (3.59%)<br>CGP (2.99%)<br>NO3 (2.4%)<br>AMP (1.8%)<br>SO4 (1.8%)<br>HDA (1.8%)<br>Other (23.4%) |

| | | | | | | | | | | | | | |
|---|---|---|---|---|---|---|---|---|---|---|---|---|---|
| **H-NQ-NQ-X** | 1 | 4 | 0.3 | papain (3.4.22.2) (3.90.70.10)<br>bleomycin hydrolase (3.4.22.40) (3.90.70.10)<br>Unknown (3.4.22.-) (3.90.70.10)<br>cathepsin K (3.4.22.38) (3.90.70.10) | 0 | 0 | 0.2 | 0 | 100 | 0 | | | 074 (13.64%)<br>D1R (9.09%)<br>2VC (9.09%)<br>MYP (9.09%)<br>9U8 (9.09%)<br>VS4 (9.09%)<br>*P* (4.55%)<br>TCK (4.55%)<br>0IW (4.55%)<br>BCQ (4.55%)<br>Other (18.2%) |
| **H-NQ-R** | 1 | 4 | 1.3 | OTC (2.1.3.3) (3.40.50.1370)<br>peroxidase (1.11.1.7) (1.10.520.10)<br>aspartate-semialdehyde dehydrogenase (1.2.1.11) (3.30.360.10)<br>N-FDH (1.17.1.9) (3.40.50.720) | 0 | 0 | 16.5 | 0 | 50 | 50 | | | CP (21.95%)<br>PAO (12.2%)<br>PO4 (9.76%)<br>NVA (7.32%)<br>FER (7.32%)<br>BHO (7.32%)<br>CYS (7.32%)<br>SO4 (4.88%)<br>ACT (4.88%)<br>CIR (2.44%)<br>Other (9.76%) |
| **H-NQ-STY** | 1 | 7 | 1.6 | pentalenene synthase (4.2.3.7) (1.10.600.10)<br>alginase (4.2.2.3) (1.50.10.100)<br>phosphohexomutase (5.3.1.8) (2.60.120.10)<br>5 (2.1.2.2) (3.40.50.170)<br>PPIase (5.2.1.8) (3.10.50.40)<br>hyaluronate lyase (4.2.2.1) (1.50.10.100)<br>meso-2 (4.2.1.28) (3.20.20.350) | 14.3 | 0 | 22.4 | 42.9 | 28.6 | 28.6 | K (16.36%)<br>CA (5.45%) | | ASG (12.73%)<br>PGO (12.73%)<br>GCT (5.45%)<br>BDP (5.45%)<br>NAG (5.45%)<br>GCU (1.82%)<br>GAR (1.82%)<br>PVC (1.82%)<br>IDR (1.82%)<br>MAN (1.82%)<br>Other (14.56%) |
| **H-R-R** | 1 | 2 | 1.2 | aspartate---tRNA ligase (6.1.1.12) (3.30.930.10)<br>phosphomonoesterase (3.1.3.2) (3.40.50.1240) | 0 | 0 | 31 | 0 | 50 | 50 | | | VO4 (12.5%)<br>PO4 (12.5%)<br>TLA (12.5%)<br>TRS (12.5%)<br>2BF (12.5%)<br>MLA (12.5%) |
| **H-R-STY** | 1 | 4 | 1.3 | swivelase (5.6.2.1) (1.20.120.380)<br>PEPPM (5.4.2.9) (3.20.20.60)<br>carbamylaspartotranskinase (2.1.3.2) (3.40.50.1370)<br>xanthan lyase (4.2.2.12) (1.50.10.100) | 0 | 0 | 31.2 | 0 | 50 | 50 | | | ASG (9.38%)<br>PCT (7.81%)<br>CP (6.25%)<br>ASP (4.69%)<br>GCT (4.69%)<br>*P* (3.12%)<br>OXL (3.12%)<br>PO4 (3.12%)<br>PAL (3.12%)<br>MLI (3.12%)<br>Other (28.08%) |
| **H-R-Z** | 1 | 2 | 1.8 | FDHH (1.17.98.4) (3.40.228.10)<br>arylsulfatase (3.1.6.1) (3.40.720.10) | 30 | 50 | 17.5 | 100 | 0 | 0 | 6MO (13.64%) | MGD (22.73%) | DDZ (36.36%)<br>FGP (9.09%)<br>SO4 (9.09%)<br>NO2 (4.55%)<br>ALS (4.55%) |
| **H-STY-STY** | 1 | 10 | 1.9 | methylmalonyl-CoA mutase (5.4.99.2) (3.20.20.240)<br>OYE (1.6.99.1) (3.20.20.70)<br>cytochrome-b5 reductase (1.6.2.2) (2.40.30.10)<br>adenosylhomocysteinase (3.3.1.1) (3.40.50.1480)<br>8-lipoxygenase (1.13.11.40) (2.40.180.10)<br>2' (3.1.4.37) (3.90.1140.10)<br>scytalone dehydratase (4.2.1.94) (3.10.450.50)<br>UDP-sulfoquinovose synthase (3.13.1.1) (3.40.50.720)<br>caspase-3 (3.4.22.56) (3.40.50.1460)<br>[histone H3]-lysine4 N-methyltransferase (2.1.1.364)<br>(2.170.270.10) | 0.4 | 35 | 16.6 | 10 | 80 | 10 | CL (0.94%) | FMN (10.8%)<br>B12 (2.82%)<br>HEM (0.94%)<br>MLC (0.47%)<br>MCA (0.47%)<br>SCA (0.47%)<br>3CP (0.47%)<br>2CP (0.47%)<br>FNR (0.47%)<br>FAD (0.47%)<br>Other (0%) | *P* (51.64%)<br>FMN (3.76%)<br>HBA (3.29%)<br>07V (1.88%)<br>0WV (1.88%)<br>5AD (0.94%)<br>TNF (0.94%)<br>NCA (0.94%)<br>3RN (0.94%)<br>BFS (0.94%)<br>Other (8.93%) |
| **H-STY-X** | 1 | 5 | 1.5 | PUNPI (2.4.2.1) (3.40.50.1580)<br>AANAT (2.3.1.87) (3.40.630.30)<br>FabD (2.3.1.39) (3.40.366.10)<br>Unknown (3.1.1.-) (3.40.50.1110)<br>2' (3.1.4.37) (3.90.1140.10) | 0 | 0 | 12.4 | 0 | 100 | 0 | | | SO4 (6.9%)<br>PO4 (5.17%)<br>DXX (5.17%)<br>IMH (3.45%)<br>DIH (3.45%)<br>ACT (3.45%)<br>DSJ (1.72%)<br>R1P (1.72%) |

| | | | | | | | | | | | | | |
|---|---|---|---|---|---|---|---|---|---|---|---|---|---|
| | | | | | | | | | | | | | DTS (1.72%)<br>CTN (1.72%)<br>Other (6.88%) |
| **H-STY-X-X** | 1 | 3 | 1.4 | destabilase (3.5.1.44) (3.40.50.180)<br>Unknown (3.4.21.-) (3.30.750.44)<br>PHP (3.9.1.3) (3.50.20.20) | 0 | 0 | 33.3 | 0 | 100 | 0 | | | PO4 (75.0%)<br>CHM (7.14%)<br>*P* (7.14%)<br>DKT (3.57%)<br>FMT (3.57%)<br>SO4 (3.57%) |
| **H-X-X** | 1 | 5 | 1.6 | pyruvate (2.7.9.1) (3.30.1490.20)<br>caspase-3 (3.4.22.56) (3.40.50.1460)<br>rhodanese (2.8.1.1) (3.40.250.10)<br>caspase-1 (3.4.22.36) (3.40.50.1460)<br>DBP2 (3.6.4.13) (2.40.10.10) | 0.4 | 0 | 13.6 | 0 | 80 | 20 | CL (0.44%)<br>ZN (0.44%) | | *P* (24.89%)<br>0QE (23.14%)<br>ASJ (3.93%)<br>DTT (2.18%)<br>PJE (2.18%)<br>010 (2.18%)<br>ECC (2.18%)<br>1U8 (1.75%)<br>0JU (1.75%)<br>HSV (1.31%)<br>Other (33.35%) |
| **H-X-X-X** | 1 | 2 | 0.3 | caspase-3 (3.4.22.56) (3.40.50.1460)<br>caspase-3 (3.4.22.56) (3.40.50.1460) | 0 | 0 | 28.5 | 0 | 100 | 0 | | | *P* (64.63%)<br>0QE (8.54%)<br>MX4 (2.44%)<br>PZN (2.44%)<br>Y2Y (2.44%)<br>158 (2.44%)<br>161 (2.44%)<br>NA3 (1.22%)<br>XVE (1.22%)<br>CNE (1.22%)<br>Other (8.54%) |
| **K-NQ-R** | 1 | 2 | 1.5 | 3-amino-5-hydroxybenzoate synthase (4.2.1.144) (3.40.640.10)<br>2 (4.1.1.64) (3.90.1150.10) | 0 | 34.5 | 22 | 0 | 100 | 0 | | PLP (15.0%)<br>PMP (5.0%) | PXG (5.0%)<br>5PA (5.0%)<br>EPC (5.0%)<br>HCP (5.0%)<br>ELP (5.0%)<br>MPM (5.0%)<br>DCS (5.0%)<br>LCS (5.0%)<br>NMA (5.0%) |
| **K-NQ-STY** | 2 | 4 | 1.1 | DHODH (1.3.5.2) (3.20.20.70)<br>cyclohydrolase (3.5.4.9) (3.40.50.10860)<br>L-DEX (3.8.1.2) (3.40.50.1000)<br>beta-hydroxyoctanoyl-ACP-dehydrase (4.2.1.59) (3.40.50.720) | 0 | 6.8 | 47.5 | 0 | 75 | 25 | | FMN (18.18%)<br>FNR (1.52%) | ORO (66.67%)<br>DOR (4.55%)<br>FMT (2.27%)<br>L34 (1.52%)<br>KUN (0.76%)<br>KUK (0.76%)<br>9L9 (0.76%)<br>MTX (0.76%)<br>2OP (0.76%)<br>NDP (0.76%)<br>Other (0%) |
| **K-R-R** | 1 | 5 | 1.9 | pyruvate (2.7.9.1) (3.30.1490.20)<br>NADPH-sulfite reductase (1.8.1.2) (3.30.413.10)<br>swivelase (5.6.2.1) (3.90.15.10)<br>myokinase (2.7.4.3) (3.40.50.300)<br>pectolyase (4.2.2.10) (2.160.20.10) | 4.2 | 20 | 61.4 | 0 | 40 | 60 | CL (3.4%) | SRM (16.33%) | AP5 (35.37%)<br>ADP (12.24%)<br>PO4 (4.76%)<br>AMP (4.08%)<br>SO4 (2.72%)<br>SO3 (2.72%)<br>C5P (2.72%)<br>NO2 (1.36%)<br>*P* (1.36%)<br>AF3 (1.36%)<br>Other (8.84%) |
| **K-R-STY** | 1 | 7 | 1.5 | PEPK (4.1.1.49) (3.90.228.20)<br>AlaDH (1.4.1.1) (3.40.50.720)<br>pyruvate kinase (2.7.1.40) (3.20.20.60)<br>phosphohexomutase (5.3.1.8) (2.60.120.10)<br>trans (4.2.3.6) (1.10.600.10)<br>AvDH1 (3.6.4.12) (3.40.50.300)<br>duplicase (2.7.7.7) (2.30.40.20) | 1.9 | 0 | 51.3 | 14.3 | 0 | 85.7 | MN (2.99%) | | OXL (22.39%)<br>PYR (13.43%)<br>POP (8.96%)<br>THJ (5.97%)<br>ATP (5.97%)<br>ADP (5.97%)<br>GOL (4.48%)<br>CO2 (2.99%)<br>OXD (2.99%) |

| | | | | | | | | | | | | | |
|---|---|---|---|---|---|---|---|---|---|---|---|---|---|
| | | | | | | | | | | | | | FLC (2.99%)<br>Other (20.9%) |
| **K-STY-STY** | 2 | 11 | 1.4 | galactowaldenase (5.1.3.2) (3.40.50.720)<br>chalcone isomerase (5.5.1.6) (3.50.70.10)<br>dTDP-glucose 4 (4.2.1.46) (3.40.50.720)<br>Unknown (1.1.1.30) (3.40.50.720)<br>4-hydroxy-tetrahydrodipicolinate synthase (4.3.3.7) (3.20.20.70)<br>17beta (1.1.1.62) (3.40.50.720)<br>(R)-oxynitrilase (4.1.2.10) (3.50.50.60)<br>Glu-AdT (6.3.5.7) (3.90.1300.10)<br>ATP:polynucleotidylexotransferase (2.7.7.19) (1.10.1410.10)<br>D-cysteine desulfhydrase (4.4.1.15) (3.40.50.1100)<br>ADP-glyceromanno-heptose 6-epimerase (5.1.3.20) (3.40.50.720) | 0 | 32.7 | 14.8 | 0 | 72.7 | 27.3 | | NAD (36.65%)<br>NAP (6.76%)<br>NDP (2.85%)<br>NAI (1.42%)<br>NAE (0.36%)<br>NDC (0.36%)<br>NAQ (0.36%)<br>PMP (0.36%) | NAD (8.19%)<br>NDP (6.41%)<br>NAP (3.91%)<br>DAU (1.78%)<br>3AT (1.78%)<br>NAI (1.42%)<br>GDD (1.07%)<br>EMO (1.07%)<br>GLN (1.07%)<br>TDX (0.71%)<br>Other (18.97%) |
| **K-STY-X** | 2 | 6 | 1.5 | beta-lactamase (3.5.2.6) (3.40.710.10)<br>glutaminase (3.5.1.2) (1.10.1500.10)<br>Unknown (3.1.1.-) (3.40.710.10)<br>SPC (3.4.21.89) (2.10.109.10)<br>ingensin (3.4.25.1) (3.60.20.10)<br>HslUV (3.4.25.2) (3.60.20.10) | 0.3 | 0 | 18.5 | 0 | 100 | 0 | NA (0.57%)<br>K (0.57%) | | DOD (5.11%)<br>SO4 (4.55%)<br>BO2 (4.55%)<br>NXL (3.41%)<br>GLU (3.41%)<br>04C (3.41%)<br>PO4 (2.84%)<br>*P* (2.84%)<br>EPE (2.27%)<br>3BV (2.27%)<br>Other (59.81%) |
| **K-X-X** | 1 | 3 | 1.3 | phosphooxymethylpyrimidine kinase (2.7.4.7) (3.40.1190.20)<br>repressor LexA (3.4.21.88) (2.10.109.10)<br>isopentenyl phosphate kinase (2.7.4.26) (3.40.1160.10) | 0 | 0 | 40.7 | 0 | 33.3 | 66.7 | | | ACP (9.09%)<br>ADP (9.09%)<br>ATP (9.09%)<br>IP8 (9.09%) |
| **NQ-R-STY** | 1 | 2 | 1.4 | NAD+-malic enzyme (1.1.1.38) (3.40.50.10380)<br>uricase (1.7.3.3) (3.10.270.10) | 1 | 17.5 | 79.5 | 0 | 100 | 0 | CL (1.71%) | NAD (5.13%)<br>NAP (0.85%) | AZA (39.32%)<br>URC (6.84%)<br>AZI (5.13%)<br>OXY (4.27%)<br>OXL (3.42%)<br>TTN (3.42%)<br>MUA (3.42%)<br>MAK (2.56%)<br>XDS (2.56%)<br>IUP (2.56%)<br>Other (17.9%) |
| **NQ-STY-STY** | 1 | 3 | 1.4 | DHODH (1.3.5.2) (3.20.20.70)<br>DD-peptidase (3.4.16.4) (3.40.710.10)<br>aspartase (4.3.1.1) (1.20.200.10) | 0 | 0 | 43 | 0 | 66.7 | 33.3 | | | ORO (52.83%)<br>HE0 (1.89%)<br>2PB (1.89%)<br>BSA (1.89%)<br>REY (1.89%)<br>0MU (1.89%)<br>PNM (1.89%)<br>RE1 (1.89%)<br>CP5 (1.89%)<br>HEL (1.89%)<br>Other (11.34%) |
| **NQ-STY-X** | 1 | 3 | 1.7 | modification methylase (2.1.1.72) (3.40.50.150)<br>DD-peptidase (3.4.16.4) (1.10.3810.10)<br>2-hydroxy-6-oxohepta-2 (3.7.1.9) (3.40.50.1820) | 20.7 | 25 | 12.3 | 33.3 | 33.3 | 33.3 | MG (10.0%)<br>CL (10.0%) | SAM (10.0%)<br>SAH (10.0%)<br>SFG (10.0%) | M0E (20.0%)<br>*P* (10.0%)<br>LHI (10.0%) |
| **NQ-X-X** | 1 | 2 | 1.4 | Dtd2 (3.1.1.96) (3.50.80.10)<br>glycylpeptide N-tetradecanoyltransferase (2.3.1.97) (3.40.630.30) | 0 | 43.5 | 54 | 0 | 50 | 50 | | MYA (41.12%)<br>NHW (16.82%)<br>NHM (2.8%)<br>YNC (0.93%)<br>COA (0.93%) | *P* (22.43%)<br>MYA (5.61%)<br>D3Y (1.87%)<br>NHW (1.87%)<br>DSN (0.93%)<br>A3G (0.93%)<br>GOL (0.93%)<br>A62 (0.93%)<br>ENF (0.93%) |
| **R-R-R** | 2 | 4 | 1.2 | myokinase (2.7.4.3) (3.40.50.300)<br>UDP-sugar diphosphatase (3.6.1.45) (3.90.780.10)<br>arginine kinase (2.7.3.3) (3.30.590.10)<br>SerRS (6.1.1.11) (3.30.930.10) | 1 | 0 | 68.2 | 0 | 0 | 100 | MG (0.67%) | | AP5 (34.9%)<br>ADP (28.19%)<br>AMP (7.38%)<br>SSA (4.7%)<br>C5P (2.68%) |

| | | | | | | | | | | | | | |
|---|---|---|---|---|---|---|---|---|---|---|---|---|---|
| | | | | | | | | | | | | | ATP (2.68%)<br>ANP (1.34%)<br>AF3 (1.34%)<br>ALF (1.34%)<br>WO4 (1.34%)<br>Other (4.69%) |
| **R-STY-STY** | 1 | 2 | 1.8 | PEPK (4.1.1.49) (3.90.228.20)<br>proton-translocating NAD(P)+ transhydrogenase (7.1.1.1) (3.40.50.1220) | 10 | 50 | 36.5 | 50 | 0 | 50 | MG (8.57%) | NAP (31.43%)<br>NDP (17.14%)<br>TXP (2.86%)<br>TAP (2.86%) | ATP (25.71%)<br>ADP (5.71%)<br>AF3 (5.71%) |
| **R-X-X** | 1 | 2 | 1.8 | formate---tetrahydrofolate ligase (6.3.4.3) (3.40.50.300)<br>deoxyuridine-triphosphatase (3.6.1.23) (2.70.40.10) | 0 | 0 | 37.5 | 0 | 0 | 100 | | | SO4 (85.71%)<br>EDO (14.29%) |
| **STY-STY-STY** | 1 | 3 | 1.9 | N4-(beta-N-acetylglucosaminyl)-L-asparaginase (3.5.1.26) (3.60.20.30)<br>dehydrogenase (1.1.2.3) (3.20.20.70)<br>ubiquinol oxidase (H+-transporting) (7.1.1.3) (1.20.210.10) | 0 | 29 | 11 | 0 | 66.7 | 33.3 | | FMN (68.18%)<br>FNS (4.55%)<br>FNR (2.27%) | FMN (9.09%)<br>GLY (6.82%)<br>GOL (2.27%)<br>ASP (2.27%)<br>ASN (2.27%)<br>NAG (2.27%) |
| **STY-STY-X** | 1 | 2 | 1.1 | [histone H3]-trimethyl-L-lysine36 demethylase (1.14.11.69) (2.60.120.650)<br>endothiapepsin (3.4.23.22) (2.40.70.10) | 0 | 0 | 19.5 | 0 | 0 | 100 | | | M3L (28.0%)<br>EDO (14.0%)<br>MLZ (4.0%)<br>MLY (4.0%)<br>MMK (2.0%)<br>5V1 (2.0%)<br>5U8 (2.0%)<br>5YQ (2.0%)<br>2MR (2.0%)<br>FQ5 (2.0%)<br>Other (22.0%) |
| **STY-W-X** | 1 | 2 | 1.1 | transglutaminase (2.3.2.13) (3.90.260.10)<br>CocE (3.1.1.84) (3.40.50.1820) | 0 | 0 | 11.5 | 0 | 50 | 50 | | | DBC (66.67%)<br>BEZ (22.22%)<br>PBC (11.11%) |
| **STY-X-X** | 2 | 11 | 1.3 | Unknown (4.1.1.-) (3.90.226.10)<br>beta-lactamase (3.5.2.6) (3.40.710.10)<br>Transferred to 3.3.2.9 and 3.3.2.10 (3.3.2.3) (3.40.50.1820)<br>glutaminase (3.5.1.2) (1.10.1500.10)<br>chymotrypsin (3.4.21.1) (2.40.10.10)<br>semacylase (3.5.1.11) (3.60.20.10)<br>EAS (4.2.3.61) (1.10.600.10)<br>PEPPM (5.4.2.9) (3.20.20.60)<br>D-aminopeptidase (3.4.11.19) (3.60.70.12)<br>Glu-AdT (6.3.5.7) (3.90.1300.10)<br>arsenite-transporting ATPase (7.3.2.7) (3.40.50.300) | 6.3 | 0 | 22.9 | 0 | 63.6 | 36.4 | CL (2.58%)<br>MG (2.58%)<br>ZN (0.65%)<br>F (0.65%) | | GLU (3.87%)<br>GLN (2.58%)<br>EDO (2.58%)<br>ADP (2.58%)<br>ONL (1.94%)<br>PAC (1.94%)<br>GRO (1.94%)<br>ASN (1.94%)<br>MPD (1.29%)<br>0G6 (1.29%)<br>Other (59.71%) |
| **STY-X-X-X** | 1 | 2 | 0.2 | acetylcholinesterase (3.1.1.7) (3.40.50.1820)<br>acetylcholinesterase (3.1.1.7) (3.40.50.1820) | 0 | 0 | 7.5 | 0 | 100 | 0 | | | VX (8.06%)<br>DPF (6.45%)<br>CO3 (4.84%)<br>ELT (4.84%)<br>PGE (3.23%)<br>DEP (3.23%)<br>UNX (3.23%)<br>03S (3.23%)<br>NWA (3.23%)<br>L2Y (3.23%)<br>Other (48.3%) |
| **X-X-X** | 1 | 5 | 1.2 | trypsin (3.4.21.4) (2.40.10.10)<br>Unknown (2.3.1.-) (3.30.1600.10)<br>lysine 2 (5.4.3.2) (3.20.20.70)<br>phosphooxymethylpyrimidine kinase (2.7.4.7) (3.40.1190.20)<br>Transferred to 5.6.1.3 (3.6.4.4) (3.40.850.10) | 0 | 20 | 9.6 | 0 | 0 | 80 | | PLP (1.63%) | *P* (9.76%)<br>ADP (1.63%)<br>K7J (0.81%)<br>K7I (0.81%)<br>7P0 (0.81%)<br>0G6 (0.81%)<br>SRB (0.81%)<br>907 (0.81%)<br>BBL (0.81%)<br>5JM (0.81%)<br>Other (49.41%) |

| Module | Number of 3D clusters | Number of templates | Mean RMSD | Enzymes (Name (EC) (CATH)) | Ligand scores (%) | | | Role class propensities (%) | | | Observed ligands | | |
|---|---|---|---|---|---|---|---|---|---|---|---|---|---|
| | | | | | Metals | Co-factors | Substrates | Metal Ligand | Reactant | Spectator | Metals | Co-factors | Substrates |
| **A-X-X** | 1 | 2 | 1.9 | 2-hydroxy-6-oxohepta-2 (3.7.1.9) (3.40.50.1820)<br>omptin (3.4.23.49) (2.40.128.90) | 0 | 0 | 25 | 0 | 100 | 0 | | | HPK (13.04%)<br>MLI (8.7%)<br>C1E (8.7%)<br>BUA (4.35%)<br>PPI (4.35%)<br>KEM (4.35%)<br>PCS (4.35%)<br>C0E (4.35%)<br>BEZ (4.35%)<br>IVA (4.35%)<br>Other (39.15%) |
| **C-C-C** | 1 | 6 | 1.1 | Gfa (4.4.1.22) (3.90.1590.10)<br>NADPH-sulfite reductase (1.8.1.2) (3.30.413.10)<br>7-carboxy-7-deazaguanine synthase (4.3.99.3) (3.20.20.70)<br>biotin synthase (2.8.1.6) (3.20.20.70)<br>RlmN (2.1.1.192) (3.20.20.70)<br>biotin synthase (2.8.1.6) (3.20.20.70) | 5.5 | 91 | 36.8 | 100 | 0 | 0 | ZN (2.13%) | SF4 (53.19%)<br>SAM (19.15%)<br>GSH (4.26%)<br>FES (4.26%) | SF4 (17.02%) |
| **C-C-C-C** | 2 | 4 | 1.5 | Unknown (2.1.1.n11) (3.40.10.10)<br>ADH3 (1.1.1.284) (3.90.180.10)<br>NADPH-sulfite reductase (1.8.1.2) (3.30.413.10)<br>ferredoxin:thioredoxin reductase (1.8.7.2) (3.90.460.10) | 50 | 19.8 | 30.2 | 100 | 0 | 0 | ZN (47.92%) | SF4 (39.58%) | SF4 (12.5%) |
| **C-C-C-H** | 1 | 2 | 1.3 | MetRS (6.1.1.10) (2.170.220.10)<br>Q-insertase (ambiguous) (2.4.2.29) (3.20.20.105) | 99.5 | 0 | 0 | 100 | 0 | 0 | ZN (100.0%) | | |
| **C-C-DE** | 1 | 5 | 1.6 | diaminopimelate epimerase (5.1.1.7) (3.10.310.10)<br>thioredoxin-disulfide reductase (1.8.1.9) (3.50.50.60)<br>ADH3 (1.1.1.284) (3.90.180.10)<br>protein disulfide-isomerase (5.3.4.1) (3.40.30.10)<br>SerRS (6.1.1.11) (3.30.930.10) | 40 | 12.8 | 12 | 40 | 60 | 0 | ZN (27.91%) | FAD (37.21%)<br>NAD (4.65%)<br>FDA (2.33%) | FAD (16.28%)<br>SER (4.65%)<br>ATP (2.33%) |
| **C-C-DE-H** | 1 | 2 | 1.9 | cytidine deaminase (3.5.4.5) (3.40.140.10)<br>anhydrase (4.2.1.1) (3.40.1050.10) | 93 | 0 | 35.5 | 100 | 0 | 0 | ZN (88.0%)<br>CO (2.0%) | | DHZ (2.0%)<br>URI (2.0%)<br>ZEB (2.0%)<br>CTD (2.0%)<br>U (2.0%) |
| **C-C-H** | 1 | 3 | 1.4 | glutathione-disulfide reductase (1.8.1.7) (3.50.50.60)<br>ADH (1.1.1.1) (3.90.180.10)<br>ferredoxin:thioredoxin reductase (1.8.7.2) (3.90.460.10) | 33.3 | 44.7 | 46 | 66.7 | 33.3 | 0 | ZN (31.16%)<br>CD (0.59%)<br>AU (0.3%)<br>CU (0.3%) | FAD (19.29%)<br>NAD (12.17%)<br>NAJ (3.26%)<br>GSH (0.89%)<br>NAP (0.89%)<br>NAI (0.59%)<br>NDP (0.59%)<br>GDS (0.3%)<br>CND (0.3%)<br>8ID (0.3%)<br>Other (0%) | FAD (4.15%)<br>ETF (2.97%)<br>PFB (2.37%)<br>NAD (1.19%)<br>FU2 (0.89%)<br>NAJ (0.89%)<br>NAI (0.89%)<br>ELI (0.59%)<br>PYZ (0.59%)<br>ACT (0.59%)<br>Other (9.56%) |
| **C-C-X** | 1 | 3 | 1.8 | beta-ketothiolase (2.3.1.16) (3.40.47.10)<br>ArsC (ambiguous) (1.20.4.4) (3.40.50.2300)<br>prostaglandin-E synthase (5.3.99.3) (3.40.30.10) | 0 | 4.7 | 0.7 | 0 | 100 | 0 | | COA (33.33%)<br>ACO (33.33%) | PN5 (8.33%)<br>ACO (8.33%)<br>SCY (8.33%) |
| **C-DE-DE** | 2 | 4 | 1.2 | 3 (4.1.99.12) (3.90.870.10)<br>glucosylceramidase (3.2.1.45) (3.20.20.80)<br>MsrA (1.8.4.11) (3.30.1060.10)<br>thiamine pyridinylase (2.5.1.2) (3.40.190.10) | 0 | 0 | 32.8 | 0 | 100 | 0 | | | 5RP (7.41%)<br>3RK (7.41%)<br>IFM (7.41%)<br>HKW (7.41%)<br>RJR (7.41%)<br>NBV (3.7%)<br>3RI (3.7%)<br>NND (3.7%)<br>AMF (3.7%)<br>LGS (3.7%)<br>Other (29.6%) |

| | | | | | | | | | | | | | |
|---|---|---|---|---|---|---|---|---|---|---|---|---|---|
| **C-DE-H** | 2 | 9 | 1.3 | NO-forming) (1.7.2.1) (2.60.40.420)<br>methylated-DNA---[protein]-cysteine S-methyltransferase (2.1.1.63) (1.10.10.10)<br>arylamine N-acetyltransferase (2.3.1.5) (2.40.128.150)<br>conjugase (3.4.19.9) (3.40.50.880)<br>Unknown (3.4.22.-) (3.40.395.10)<br>carboxymethylenebutenolidase (3.1.1.45) (3.40.50.1820)<br>selenocysteine lyase (4.4.1.16) (3.40.640.10)<br>glutamate racemase (5.1.1.3) (3.40.50.1860)<br>cytosine deaminase (3.5.4.1) (3.40.140.10) | 11.2 | 0 | 4.1 | 22.2 | 66.7 | 11.1 | ZN (55.56%)<br>NA (3.7%) | | DOD (11.11%)<br>FMT (7.41%)<br>DUC (7.41%)<br>HPY (7.41%)<br>ACT (3.7%)<br>GOL (3.7%) |
| **C-DE-H-NQ** | 1 | 2 | 2.5 | adenosylhomocysteinase (3.3.1.1) (3.40.50.720)<br>TRS (6.1.1.3) (3.30.930.10) | 36.5 | 50 | 20.5 | 50 | 50 | 0 | ZN (34.88%)<br>K (2.33%) | NAD (16.28%)<br>NAI (2.33%) | 2CR (9.3%)<br>TSB (9.3%)<br>SSA (6.98%)<br>1B2 (4.65%)<br>409 (2.33%)<br>THR (2.33%)<br>SO4 (2.33%)<br>1B3 (2.33%)<br>BC9 (2.33%)<br>X16 (2.33%)<br>Other (0%) |
| **C-DE-R** | 1 | 2 | 1.4 | MoCu-CODH (1.2.5.3) (3.30.365.10)<br>anhydrase (4.2.1.1) (3.40.1050.10) | 22.5 | 50 | 5 | 100 | 0 | 0 | ZN (35.48%)<br>CU (3.23%)<br>CO (3.23%) | MCN (22.58%)<br>CUB (6.45%) | BCT (12.9%)<br>CUM (6.45%)<br>CUN (6.45%)<br>OMO (3.23%) |
| **C-DE-STY** | 1 | 7 | 1.8 | glutamate racemase (5.1.1.3) (3.40.50.1860)<br>arginine kinase (2.7.3.3) (3.30.590.10)<br>ferredoxin-NADP+-oxidoreductase (1.18.1.2) (3.40.50.80)<br>7-carboxy-7-deazaguanine synthase (4.3.99.3) (3.20.20.70)<br>phosphopentosisomerase (5.3.1.6) (3.40.1400.10)<br>YkvM (1.7.1.13) (3.30.1130.10)<br>vinylacetyl-CoA Delta-isomerase (5.3.3.3) (2.40.110.10) | 0 | 18.3 | 33.3 | 14.3 | 85.7 | 0 | | FAD (32.61%)<br>SAM (13.04%)<br>SF4 (13.04%) | ARG (8.7%)<br>DAR (6.52%)<br>ORN (2.17%)<br>3S5 (2.17%)<br>IOM (2.17%)<br>ILO (2.17%)<br>R5P (2.17%)<br>RB5 (2.17%)<br>AOS (2.17%)<br>R52 (2.17%)<br>Other (6.51%) |
| **C-H-H** | 1 | 4 | 1.3 | GTP cyclohydrolase (3.5.4.16) (3.30.1130.10)<br>PPIase (5.2.1.8) (3.10.50.40)<br>neelaredoxin (1.15.1.2) (2.60.40.730)<br>mannonate dehydratase (4.2.1.8) (3.20.20.150) | 54.5 | 0 | 13 | 75 | 25 | 0 | ZN (33.33%)<br>FE (23.08%)<br>MN (12.82%)<br>CA (5.13%) | | GTP (7.69%)<br>8GT (5.13%)<br>CS2 (5.13%)<br>8DG (2.56%)<br>IPA (2.56%)<br>QBQ (2.56%) |
| **C-H-H-STY** | 1 | 2 | 2.2 | PPIase (5.2.1.8) (3.10.50.40)<br>N-acetylmuramoyl-L-alanine amidase (3.5.1.28) (3.40.80.10) | 50 | 0 | 0 | 0 | 100 | 0 | ZN (100.0%) | | |
| **C-H-NQ** | 2 | 6 | 1.5 | DOCS (2.3.1.74) (3.40.47.10)<br>F1-ATPase (7.1.2.2) (2.170.16.10)<br>cathepsin L (3.4.22.15) (3.90.70.10)<br>beta-lactamase (3.5.2.6) (3.60.15.10)<br>GTP cyclohydrolase (3.5.4.16) (3.30.1130.10)<br>cathepsin K (3.4.22.38) (3.90.70.10) | 10.8 | 0 | 12.3 | 33.3 | 66.7 | 0 | ZN (35.39%)<br>CD (0.91%)<br>CL (0.68%)<br>AU (0.23%) | | MRI (12.79%)<br>RTD (4.34%)<br>*P* (1.6%)<br>E64 (1.37%)<br>ZZ7 (1.14%)<br>OH (0.91%)<br>QUE (0.91%)<br>MYC (0.91%)<br>COA (0.68%)<br>0IW (0.68%)<br>Other (29.66%) |
| **C-H-STY** | 2 | 7 | 1.7 | protein-tyrosine-phosphatase (3.1.3.48) (3.90.190.10)<br>ADH3 (1.1.1.284) (3.90.180.10)<br>MDR (1.1.1.105) (3.90.180.10)<br>ADH (1.1.1.1) (3.90.180.10)<br>3 (4.1.99.12) (3.90.870.10)<br>vinylacetyl-CoA Delta-isomerase (5.3.3.3) (1.20.140.10)<br>destabilase (3.5.1.44) (3.30.1330.200) | 14.3 | 17.9 | 5 | 42.9 | 57.1 | 0 | ZN (6.48%) | NAD (42.59%)<br>NAJ (10.19%)<br>NAP (3.7%)<br>NAI (1.85%)<br>NDP (1.85%)<br>CND (0.93%)<br>8ID (0.93%)<br>PAD (0.93%) | NAD (3.7%)<br>CXF (2.78%)<br>NAJ (2.78%)<br>NAI (2.78%)<br>MRD (1.85%)<br>ETF (1.85%)<br>COD (1.85%)<br>SF4 (1.85%)<br>PVS (0.93%)<br>PSY (0.93%)<br>Other (4.65%) |

| | | | | | | | | | | | | | |
|---|---|---|---|---|---|---|---|---|---|---|---|---|---|
| **C-NQ-X** | 1 | 3 | 1.6 | peptide deformylase (3.5.1.88) (3.90.45.10)<br>amidophosphoribosyltransferase (2.4.2.14) (3.60.20.10)<br>ferredoxin-glutamate synthase (1.4.7.1) (3.60.20.10) | 33.3 | 0 | 24.3 | 33.3 | 66.7 | 0 | ZN (32.33%)<br>NI (17.24%)<br>CD (5.6%)<br>CO (4.74%)<br>FE (1.72%)<br>FE2 (0.86%)<br>CL (0.43%) | | BB2 (19.4%)<br>FMT (3.02%)<br>ONL (1.72%)<br>LHY (1.29%)<br>K3U (1.29%)<br>K1U (1.29%)<br>K2U (1.29%)<br>SB7 (0.86%)<br>MLN (0.86%)<br>PO4 (0.86%)<br>Other (3.87%) |
| **C-NQ-X-X-X** | 1 | 2 | 1.6 | hexosephosphate aminotransferase (2.6.1.16) (3.60.20.10)<br>GOGAT (1.4.1.13) (3.60.20.10) | 0 | 0 | 19.5 | 0 | 100 | 0 | | | ONL (100.0%) |
| **C-R-STY** | 1 | 3 | 1.7 | sulfite oxidase (1.8.3.1) (3.90.420.10)<br>McyF (5.1.1.13) (3.40.50.1860)<br>protein-tyrosine-phosphatase (3.1.3.48) (3.90.190.10) | 33.3 | 0 | 42.7 | 33.3 | 66.7 | 0 | MO (5.41%) | | SO4 (5.41%)<br>073 (5.41%)<br>PO4 (5.41%)<br>UA1 (5.41%)<br>234 (2.7%)<br>OTA (2.7%)<br>IX1 (2.7%)<br>F20 (2.7%)<br>F32 (2.7%)<br>DBD (2.7%)<br>Other (45.9%) |
| **C-STY-STY** | 1 | 5 | 1.3 | pyruvate (2.7.9.1) (3.20.20.60)<br>(S)-oxynitrilase (4.1.2.47) (3.40.50.1820)<br>GDP-4-keto-6-deoxy-D-mannose-3 (1.1.1.271) (3.40.50.720)<br>riboflavin synthase (2.5.1.9) (2.40.30.20)<br>ribonucleoside-triphosphate reductase (formate) (1.1.98.6)<br>(3.20.70.20) | 0 | 2.8 | 34.6 | 0 | 100 | 0 | | NAP (8.7%) | ACN (13.04%)<br>SCN (8.7%)<br>GOL (8.7%)<br>GFB (8.7%)<br>CNH (4.35%)<br>MNN (4.35%)<br>FAC (4.35%)<br>ACT (4.35%)<br>CLX (4.35%)<br>ICN (4.35%)<br>Other (26.1%) |
| **C-STY-X** | 1 | 2 | 1.2 | N-carbamoylsarcosine amidase (3.5.1.59) (3.40.50.850)<br>adenosylmethionine decarboxylase (4.1.1.50) (3.60.90.10) | 0 | 0 | 0 | 0 | 100 | 0 | | | SO4 (100.0%) |
| **C-X-X** | 1 | 3 | 1.7 | D-camphor-exo-hydroxylase (1.14.15.1) (1.10.630.10)<br>beta-hydroxyoctanoyl-ACP-dehydrase (4.2.1.59) (3.40.47.10)<br>rhodanese (2.8.1.1) (3.40.250.10) | 0 | 27 | 2 | 33.3 | 66.7 | 0 | | HEM (53.26%)<br>7HE (1.09%)<br>MNR (1.09%) | HEM (11.96%)<br>CER (5.43%)<br>OCA (2.17%)<br>6NA (2.17%)<br>MRJ (2.17%)<br>PNS (2.17%)<br>TLM (2.17%)<br>HEC (1.09%)<br>CMO (1.09%)<br>1XG (1.09%)<br>Other (7.63%) |
| **C-X-X-X-X** | 1 | 2 | 1.1 | ArsC (ambiguous) (1.20.4.4) (3.40.50.2300)<br>rhodanese (2.8.1.1) (3.40.250.10) | 0 | 0 | 12.5 | 0 | 100 | 0 | | | SO4 (50.0%)<br>ACT (50.0%) |
| **DE-DE-DE** | 4 | 18 | 1.4 | calcium pump (7.2.2.10) (3.40.50.1000)<br>neopullulanase (3.2.1.135) (3.20.20.80)<br>4-alpha-glucanotransferase (2.4.1.25) (3.20.20.80)<br>exodeoxyribonuclease (lambda-induced) (3.1.11.3) (3.40.50.1010)<br>glucoamylase (3.2.1.3) (1.50.10.10)<br>fumarylacetoacetase (3.7.1.2) (3.90.850.10)<br>xylose isomerase (5.3.1.5) (3.20.20.150)<br>enolase (4.2.1.11) (3.20.20.120)<br>2-chloro-2 (5.5.1.7) (3.20.20.120)<br>duplicase (2.7.7.7) (3.30.460.10)<br>transphosphoribosidase (2.4.2.8) (3.40.50.2020)<br>Unknown (2.4.2.-) (3.40.50.2020)<br>protoporphyrin ferrochelatase (4.99.1.1) (3.40.50.1400)<br>glycopeptidase (3.5.1.52) (2.60.120.230)<br>glucoamylase (3.2.1.3) (1.50.10.10)<br>isoamylase (3.2.1.68) (3.20.20.80)<br>exodeoxyribonuclease (lambda-induced) (3.1.11.3) (3.40.50.1010)<br>UDP-N-acetyl-D-glucosamine 2-epimerase (5.1.3.14)<br>(3.40.50.2000) | 24.4 | 0 | 26.8 | 50 | 50 | 0 | MG (26.19%)<br>MN (10.48%)<br>CA (1.75%)<br>CO (0.58%)<br>ZN (0.23%)<br>D8U (0.12%)<br>CL (0.12%) | | *P* (9.31%)<br>GLC (3.96%)<br>MF4 (2.44%)<br>ALF (1.51%)<br>XYL (1.51%)<br>BGC (0.93%)<br>EDO (0.93%)<br>DUP (0.93%)<br>8OG (0.81%)<br>ACP (0.7%)<br>Other (27.7%) |
| **DE-DE-DE-DE** | 1 | 12 | 2.2 | NAD+-malic enzyme (1.1.1.38) (3.40.50.10380)<br>xylose isomerase (5.3.1.5) (3.20.20.150)<br>Xaa-Pro aminopeptidase (3.4.11.9) (3.90.230.10)<br>3'(2') (3.1.3.7) (3.40.190.80)<br>alpha-1 (4.2.2.2) (2.160.20.10) | 53.5 | 2.1 | 38.2 | 83.3 | 8.3 | 8.3 | MN (22.11%)<br>MG (20.35%)<br>CA (4.27%)<br>ZN (4.02%)<br>TL (2.01%) | NAD (1.01%)<br>NAP (0.25%) | PO4 (6.28%)<br>XYL (3.27%)<br>GLC (2.76%)<br>EDO (1.51%)<br>PRO (1.51%) |

| | | | | | | | | | | | | | |
|---|---|---|---|---|---|---|---|---|---|---|---|---|---|
| | | | | 3'(2') (3.1.3.7) (3.40.190.80)<br>fructose-bisphosphatase (3.1.3.11) (3.30.540.10)<br>swivelase (5.6.2.1) (3.40.50.140)<br>exodeoxyribonuclease (lambda-induced) (3.1.11.3) (3.40.50.1010)<br>lysozyme (3.2.1.17) (3.20.20.80)<br>inorganic diphosphatase (3.6.1.1) (3.90.80.10)<br>non-chaperonin molecular chaperone ATPase (3.6.4.10)<br>(3.30.30.30) | | | | | | | CO (1.76%)<br>NA (0.75%)<br>K (0.25%) | | AMP (1.51%)<br>XLS (1.26%)<br>OXL (1.01%)<br>TTN (1.01%)<br>GOL (1.01%)<br>Other (11.51%) |
| **DE-DE-DE-DE-H** | 2 | 7 | 1.7 | neopullulanase (3.2.1.135) (3.20.20.80)<br>L-fuconate dehydratase (4.2.1.68) (3.20.20.120)<br>GS (6.3.1.2) (3.30.590.10)<br>arginase (3.5.3.1) (3.40.800.10)<br>mandelate racemase (5.1.2.2) (3.20.20.120)<br>exodeoxyribonuclease III (3.1.11.2) (3.60.10.10)<br>cis (5.5.1.1) (3.20.20.120) | 39.4 | 0 | 24.3 | 85.7 | 14.3 | 0 | MN (48.1%)<br>MG (11.96%)<br>CO (2.72%)<br>NI (1.09%)<br>CA (0.82%)<br>ZN (0.54%) | | ABH (2.99%)<br>P3S (2.17%)<br>GLC (1.9%)<br>S2C (1.9%)<br>ORN (1.36%)<br>NNH (1.09%)<br>X7A (1.09%)<br>NOJ (0.82%)<br>AC1 (0.82%)<br>HAR (0.82%)<br>Other (17.83%) |
| **DE-DE-DE-DE-H-H** | 2 | 7 | 1.9 | bacterial leucyl aminopeptidase (3.4.11.10) (3.40.630.10)<br>glutamate carboxypeptidase (3.4.17.11) (3.40.630.10)<br>arginase (3.5.3.1) (3.40.800.10)<br>L-rhamnose isomerase (5.3.1.14) (3.20.20.150)<br>catalase (1.11.1.6) (1.20.1260.10)<br>methane hydroxylase (1.14.13.25) (1.10.620.20)<br>redoxyendonuclease (3.1.21.2) (3.20.20.150) | 71.9 | 0 | 32.3 | 100 | 0 | 0 | MN (31.67%)<br>FE (23.58%)<br>ZN (11.02%)<br>CO (2.41%)<br>FE2 (1.55%)<br>MN3 (1.38%)<br>NI (0.69%)<br>O (0.34%) | | ABH (1.89%)<br>S2C (1.2%)<br>OH (1.2%)<br>ORN (0.86%)<br>LEU (0.69%)<br>BES (0.69%)<br>NNH (0.69%)<br>X7A (0.69%)<br>TRS (0.52%)<br>PLU (0.52%)<br>Other (12.43%) |
| **DE-DE-DE-DE-K** | 1 | 2 | 2.3 | cis (5.5.1.1) (3.20.20.120)<br>OSBS (4.2.1.113) (3.20.20.120) | 73.5 | 0 | 29.5 | 100 | 0 | 0 | MG (39.74%)<br>MN (11.54%)<br>CL (1.28%) | | MUC (7.69%)<br>DGL (6.41%)<br>TYR (3.85%)<br>ACY (2.56%)<br>AME (2.56%)<br>DLY (1.28%)<br>SUG (1.28%)<br>VAL (1.28%)<br>ALA (1.28%)<br>SMG (1.28%)<br>Other (7.68%) |
| **DE-DE-DE-DE-X** | 1 | 2 | 0.7 | fructose-bisphosphatase (3.1.3.11) (3.30.540.10)<br>D-galactose 1-phosphate phosphatase (3.1.3.94) (3.30.540.10) | 81 | 2.5 | 69 | 100 | 0 | 0 | MG (26.4%)<br>CA (13.71%)<br>MN (10.15%)<br>ZN (7.11%)<br>TL (5.08%)<br>GD (2.03%)<br>CL (0.51%) | NAP (1.52%) | PO4 (16.75%)<br>IPD (4.57%)<br>F6P (3.05%)<br>SO4 (2.03%)<br>PI (1.02%)<br>PO3 (1.02%)<br>FBP (1.02%)<br>FDP (0.51%)<br>INS (0.51%)<br>LIP (0.51%)<br>Other (1.02%) |
| **DE-DE-DE-H** | 2 | 12 | 1.7 | stearoyl-[acyl-carrier-protein] 9-desaturase (1.14.19.2)<br>(1.10.620.20)<br>L-fuconate dehydratase (4.2.1.68) (3.20.20.120)<br>caffeate O-methyltransferase (2.1.1.68) (3.40.50.150)<br>creatininase (3.5.2.10) (3.40.50.10310)<br>diisopropyl-fluorophosphatase (3.1.8.2) (2.120.10.30)<br>mandelate racemase (5.1.2.2) (3.20.20.120)<br>exo-cellobiohydrolase (3.2.1.91) (2.70.100.10)<br>methane hydroxylase (1.14.13.25) (1.10.620.20)<br>3 (4.1.99.12) (3.90.870.10)<br>duplicase (2.7.7.7) (3.30.420.10)<br>Unknown (3.1.-.-) (3.30.420.10)<br>cis (5.5.1.1) (3.20.20.120) | 42.3 | 0 | 18.4 | 83.3 | 16.7 | 0 | FE (35.46%)<br>MG (13.52%)<br>MN (4.59%)<br>FE2 (2.55%)<br>ZN (1.28%)<br>CA (0.77%)<br>CO (0.51%)<br>NA (0.51%)<br>CL (0.26%)<br>ER3 (0.26%)<br>Other (0%) | | BGC (4.34%)<br>FMT (0.77%)<br>*P* (0.77%)<br>MSE (0.51%)<br>MCR (0.51%)<br>AZI (0.51%)<br>PER (0.51%)<br>PCR (0.51%)<br>SO4 (0.51%)<br>TMP (0.51%)<br>Other (14.0%) |
| **DE-DE-DE-H-H** | 1 | 5 | 1.6 | DNase (3.1.21.1) (3.60.10.10)<br>exo-1 (3.2.1.114) (3.20.110.10)<br>tripeptidase (3.4.11.4) (1.10.390.10)<br>arginase (3.5.3.1) (3.40.800.10)<br>methane hydroxylase (1.14.13.25) (1.10.620.20) | 79 | 0 | 18.4 | 100 | 0 | 0 | ZN (22.43%)<br>FE (21.34%)<br>MN (14.8%)<br>FE2 (1.4%)<br>CO (1.09%)<br>NI (0.31%)<br>MG (0.16%) | | ABH (1.71%)<br>BES (1.4%)<br>S2C (1.09%)<br>LYS (0.78%)<br>ORN (0.78%)<br>NNH (0.62%)<br>X7A (0.62%)<br>HAR (0.47%)<br>MSN (0.31%) |

| | | | | | | | | | | | | | |
|---|---|---|---|---|---|---|---|---|---|---|---|---|---|
| | | | | | | | | | | | | | SWA (0.31%)<br>Other (24.2%) |
| **DE-DE-DE-H-H-H** | 1 | 2 | 1.4 | Unknown (1.-.-.-) (3.60.15.10)<br>creatininase (3.5.2.10) (3.40.50.10310) | 61 | 0 | 5 | 100 | 0 | 0 | ZN (29.55%)<br>MN (15.91%)<br>FE (9.09%)<br>O (2.27%)<br>CL (2.27%) | | FEO (15.91%)<br>OXY (9.09%)<br>CRN (2.27%) |
| **DE-DE-DE-H-H-H-STY** | 1 | 2 | 0.4 | phosphomonoesterase (3.1.3.1) (3.40.720.10)<br>2 (5.4.2.12) (3.40.720.10) | 31 | 0 | 92.5 | 100 | 0 | 0 | ZN (29.63%)<br>CO (7.41%) | | PO4 (33.33%)<br>2PG (7.41%)<br>3PG (7.41%)<br>PAE (3.7%)<br>MMQ (3.7%)<br>AF3 (3.7%)<br>WO4 (3.7%) |
| **DE-DE-DE-H-K** | 1 | 2 | 0.8 | Unknown (4.6.1.-) (3.20.20.190)<br>glycerophosphodiester phosphodiesterase (3.1.4.46)<br>(3.20.20.190) | 61.5 | 0 | 7.5 | 100 | 0 | 0 | MG (33.33%)<br>CA (11.11%)<br>FE (3.7%) | | GOL (11.11%)<br>IPD (3.7%) |
| **DE-DE-DE-H-NQ** | 1 | 2 | 1.9 | beta-methylaspartase (4.3.1.2) (3.20.20.120)<br>N-sulfoglucosamine sulfohydrolase (3.10.1.1) (3.40.720.10) | 87.5 | 0 | 6 | 100 | 0 | 0 | MG (46.15%)<br>CA (23.08%) | | FGP (23.08%)<br>2AS (7.69%) |
| **DE-DE-DE-H-R** | 1 | 4 | 2.5 | konchizaimu (2.4.1.19) (3.20.20.80)<br>MICL (4.1.3.30) (3.20.20.60)<br>Xaa-Pro aminopeptidase (3.4.11.9) (3.90.230.10)<br>endo-polygalacturonase (3.2.1.15) (2.160.20.10) | 0 | 0 | 13.8 | 50 | 50 | 0 | | | GLC (16.67%)<br>PRO (10.0%)<br>ACI (6.67%)<br>*P* (6.67%)<br>BGC (5.0%)<br>NOJ (5.0%)<br>ADH (5.0%)<br>M44 (3.33%)<br>GTR (3.33%)<br>OPG (1.67%)<br>Other (10.02%) |
| **DE-DE-DE-K** | 3 | 13 | 1.7 | enolase (4.2.1.11) (3.20.20.120)<br>OSBS (4.2.1.113) (3.20.20.120)<br>Unknown (3.6.1.74) (3.20.100.10)<br>OSBS (4.2.1.113) (3.20.20.120)<br>Unknown (5.5.1.-) (3.20.20.120)<br>NADP+-ICDH (1.1.1.42) (3.40.718.10)<br>L-fuconate dehydratase (4.2.1.68) (3.20.20.120)<br>mandelate racemase (5.1.2.2) (3.20.20.120)<br>swivelase (5.6.2.1) (3.40.50.140)<br>2-chloro-2 (5.5.1.7) (3.20.20.120)<br>beta-phosphoglucomutase (5.4.2.6) (3.40.50.1000)<br>licheninase (3.2.1.73) (3.20.20.80)<br>endo-1 (3.2.1.39) (3.20.20.80) | 44.2 | 0.2 | 23.4 | 84.6 | 15.4 | 0 | MG (40.82%)<br>MN (5.1%)<br>CA (3.32%)<br>CL (0.51%) | NAD (0.77%) | IPM (4.59%)<br>MUC (2.55%)<br>ICT (2.55%)<br>DGL (2.04%)<br>ALF (2.04%)<br>MGF (2.04%)<br>TYR (1.79%)<br>APG (1.28%)<br>AKG (1.02%)<br>LGT (1.02%)<br>Other (20.28%) |
| **DE-DE-DE-K-K** | 1 | 2 | 0.8 | Hyp-B 2-epimerase (5.1.1.22) (3.20.20.120)<br>Unknown (5.5.1.-) (3.20.20.120) | 60.5 | 0 | 27.5 | 100 | 0 | 0 | MG (48.31%)<br>MN (2.25%)<br>CL (1.12%) | | 0XW (5.62%)<br>TYR (4.49%)<br>GLU (3.37%)<br>MUC (3.37%)<br>PBE (2.25%)<br>5CR (2.25%)<br>DGL (2.25%)<br>NPQ (2.25%)<br>4OP (1.12%)<br>NSK (1.12%)<br>Other (12.32%) |
| **DE-DE-DE-NQ-Z** | 1 | 2 | 0.3 | cerebroside-sulfatase (3.1.6.8) (3.40.720.10)<br>arylsulfatase (3.1.6.1) (3.40.720.10) | 96 | 0 | 23 | 100 | 0 | 0 | CA (35.71%)<br>MG (2.38%) | | DDZ (19.05%)<br>FGP (9.52%)<br>SO4 (9.52%)<br>ALS (4.76%)<br>SV7 (4.76%)<br>FGL (2.38%)<br>62Y (2.38%) |
| **DE-DE-DE-STY** | 3 | 13 | 2.3 | (+)-bornylpyrophosphate cyclase (5.5.1.8) (1.10.600.10)<br>levansucrase (2.4.1.10) (2.115.10.20)<br>IAG-NH (3.2.2.1) (3.90.245.10)<br>cellulase (3.2.1.4) (1.50.10.10)<br>duplicase (2.7.7.7) (3.30.70.270)<br>beta-glucuronidase (3.2.1.31) (2.60.120.260)<br>exo-cellobiohydrolase (3.2.1.91) (3.20.20.40) | 16.5 | 0 | 25.6 | 30.8 | 69.2 | 0 | CA (4.35%)<br>MG (1.45%)<br>IOD (0.48%) | | AMI (5.8%)<br>IPT (4.83%)<br>NAG (4.83%)<br>149 (4.35%)<br>*P* (3.38%)<br>VR0 (2.9%)<br>NGT (2.9%) |

| | | | | | | | | | | | | | |
|---|---|---|---|---|---|---|---|---|---|---|---|---|---|
| | | | | chitinase (3.2.1.14) (3.20.20.80)<br>biliverdin reductase (1.3.1.24) (3.30.360.10)<br>chitinase (3.2.1.14) (3.20.20.80)<br>trans-deoxyribosylase (2.4.2.6) (3.40.50.450)<br>sedolisin (3.4.21.100) (3.40.50.200)<br>inorganic diphosphatase (3.6.1.1) (3.90.80.10) | | | | | | | | | A1L (2.42%)<br>GLC (1.93%)<br>3AG (1.45%)<br>Other (45.85%) |
| **DE-DE-DE-X** | 1 | 3 | 1.8 | hybridase (3.1.26.4) (3.30.420.10)<br>thermonuclease (3.1.31.1) (2.40.50.90)<br>ADPR-PPase (3.6.1.13) (3.90.79.10) | 57.7 | 0 | 54 | 100 | 0 | 0 | CA (48.0%)<br>MN (1.6%)<br>MG (0.8%)<br>CO (0.8%) | | THP (47.2%)<br>PO4 (0.8%)<br>ADV (0.8%) |
| **DE-DE-H** | 3 | 25 | 1.3 | glycine amidinotransferase (2.1.4.1) (3.75.10.10)<br>haloalkane dehalogenase (3.8.1.5) (3.40.50.1820)<br>exo-cellobiohydrolase (3.2.1.91) (3.20.20.80)<br>glyoxylate reductase (1.1.1.26) (3.40.50.720)<br>Unknown (3.4.24.-) (3.30.830.10)<br>Transferred to 3.3.2.9 and 3.3.2.10 (3.3.2.3) (3.40.50.1820)<br>PRMT1 (gene name) (2.1.1.319) (3.40.50.150)<br>YkvM (1.7.1.13) (3.30.1130.10)<br>3'-phosphoadenylyl-sulfate:[heparan sulfate]-glucosamine 3-sulfotransferase (2.8.2.30) (3.40.50.300)<br>chloroperoxidase (1.11.1.10) (1.10.489.10)<br>phosphatidase (3.1.1.4) (1.20.90.10)<br>glucosamine-6-phosphate deaminase (3.5.99.6) (3.40.50.1360)<br>L-fucose isomerase (5.3.1.25) (3.20.14.10)<br>carboxypeptidase T (3.4.17.18) (3.40.630.10)<br>Unknown (1.-.-.-) (3.60.15.10)<br>galactarate dehydratase (D-threo-forming) (4.2.1.158) (3.20.20.120)<br>Unknown (2.4.1.-) (3.90.550.10)<br>UDP-sugar diphosphatase (3.6.1.45) (3.60.21.10)<br>arylsulfatase (3.1.6.1) (3.40.720.10)<br>Unknown (3.1.5.-) (1.10.3210.10)<br>protoporphyrin ferrochelatase (4.99.1.1) (3.40.50.1400)<br>aldolase (4.1.2.13) (3.20.20.70)<br>enolase (4.2.1.11) (3.20.20.120)<br>creatinase (3.5.3.3) (3.90.230.10)<br>PdxJ (2.6.99.2) (3.20.20.70) | 25.5 | 5.5 | 25.1 | 48 | 52 | 0 | ZN (30.61%)<br>MN (5.54%)<br>CA (3.5%)<br>FE (0.58%)<br>MG (0.58%)<br>CL (0.29%)<br>HG (0.29%)<br>O (0.29%) | HEM (4.37%)<br>PXP (0.87%) | *P* (4.66%)<br>DDZ (2.33%)<br>FEO (1.75%)<br>FOC (1.46%)<br>CXA (1.17%)<br>OXY (1.17%)<br>SO4 (1.17%)<br>IPA (0.87%)<br>3K0 (0.87%)<br>GOL (0.58%)<br>Other (30.16%) |
| **DE-DE-H-H** | 1 | 16 | 2 | redoxyendonuclease (3.1.21.2) (3.20.20.150)<br>thermolysin (3.4.24.27) (1.10.390.10)<br>isopentenyl-diphosphate Delta-isomerase (5.3.3.2) (3.90.79.10)<br>deuterolysin (3.4.24.39) (3.40.390.10)<br>neprilysin (3.4.24.11) (3.40.390.10)<br>bontoxilysin (3.4.24.69) (3.90.1240.10)<br>peptidyl-Lys metalloendopeptidase (3.4.24.20) (3.40.390.10)<br>dGTPase (3.1.5.1) (1.10.3210.10)<br>redoxyendonuclease (3.1.21.2) (3.20.20.150)<br>alpha-toxin (3.1.4.3) (1.10.575.10)<br>DNase (3.1.21.1) (3.60.10.10)<br>NDO (1.14.12.12) (3.90.380.10)<br>nicotinamidase (3.5.1.19) (3.40.50.850)<br>Xaa-Pro aminopeptidase (3.4.11.9) (3.90.230.10)<br>Unknown (3.4.24.-) (3.30.830.10)<br>2 (5.4.2.12) (3.40.720.10) | 69.3 | 0 | 32.4 | 100 | 0 | 0 | ZN (48.85%)<br>MN (2.52%)<br>CD (1.61%)<br>CU (0.46%)<br>FE2 (0.46%)<br>FE (0.46%)<br>CU1 (0.23%)<br>CL (0.23%)<br>NA (0.23%)<br>RU (0.23%)<br>Other (0.23%) | | *P* (4.82%)<br>VAL (3.21%)<br>RDF (1.38%)<br>LEU (0.92%)<br>GOL (0.92%)<br>BNL (0.92%)<br>OXY (0.92%)<br>MRD (0.69%)<br>NX6 (0.69%)<br>ASP (0.69%)<br>Other (23.92%) |
| **DE-DE-H-H-H** | 1 | 4 | 2.2 | alpha-toxin (3.1.4.3) (1.10.575.10)<br>hydroxyacylglutathione hydrolase (3.1.2.6) (3.60.15.10)<br>neprilysin (3.4.24.11) (3.40.390.10)<br>adenosine deaminase (3.5.4.4) (3.20.20.140) | 94 | 2.5 | 30.8 | 100 | 0 | 0 | ZN (60.65%)<br>FE (12.26%)<br>CD (2.58%)<br>CO (0.65%)<br>NI (0.65%) | GBP (0.65%) | RDF (3.23%)<br>PRH (1.94%)<br>DCF (1.29%)<br>ADE (1.29%)<br>MCF (1.29%)<br>LAC (0.65%)<br>TLA (0.65%)<br>CAC (0.65%)<br>ACY (0.65%)<br>OIR (0.65%)<br>Other (10.4%) |
| **DE-DE-H-H-STY** | 1 | 2 | 1.3 | neurolysin (3.4.24.16) (3.40.390.10)<br>lethal toxin (3.4.24.83) (3.40.390.10) | 93.5 | 0 | 13 | 50 | 50 | 0 | ZN (53.49%) | | *P* (2.33%)<br>2ZL (2.33%)<br>GOL (2.33%)<br>56R (2.33%)<br>GM6 (2.33%)<br>407 (2.33%)<br>30H (2.33%)<br>56Q (2.33%)<br>0LX (2.33%) |

| | | | | | | | | | | | | | |
|---|---|---|---|---|---|---|---|---|---|---|---|---|---|
| | | | | | | | | | | | | | 30P (2.33%)<br>Other (23.3%) |
| **DE-DE-H-NQ** | 2 | 13 | 2 | 1-phosphatidylinositol-4 (3.1.4.11) (3.20.20.190)<br>phosphomonoesterase (3.1.3.2) (3.60.21.10)<br>exodeoxyribonuclease III (3.1.11.2) (3.60.10.10)<br>Unknown (4.6.1.-) (3.20.20.190)<br>1-phosphatidylinositol-4 (3.1.4.11) (3.20.20.190)<br>Unknown (3.1.5.-) (1.10.3210.10)<br>histidinol dehydrogenase (1.1.1.23) (3.40.50.1980)<br>beta-methylaspartase (4.3.1.2) (3.20.20.120)<br>UDP-sugar diphosphatase (3.6.1.45) (3.60.21.10)<br>cerebroside-sulfatase (3.1.6.8) (3.40.720.10)<br>UDP-sugar diphosphatase (3.6.1.45) (3.60.21.10)<br>Unknown (3.1.-.-) (3.40.1800.10)<br>exo-cellobiohydrolase (3.2.1.91) (3.20.20.80) | 53.1 | 0 | 26.2 | 84.6 | 15.4 | 0 | CA (15.32%)<br>MG (11.91%)<br>ZN (8.51%)<br>FE (5.96%)<br>MN (2.55%)<br>SM (1.7%)<br>LA (0.85%)<br>PB (0.85%)<br>BA (0.43%)<br>F (0.43%)<br>Other (0%) | | XYP (8.94%)<br>3DR (5.96%)<br>*P* (4.68%)<br>I3P (2.55%)<br>DV3 (2.55%)<br>PO4 (2.13%)<br>SO4 (1.28%)<br>XDN (1.28%)<br>XIF (1.28%)<br>IP2 (0.85%)<br>Other (15.81%) |
| **DE-DE-H-R** | 1 | 4 | 2 | fumarylacetoacetase (3.7.1.2) (3.90.850.10)<br>3-phosphoshikimate 1-carboxyvinyltransferase (2.5.1.19) (3.65.10.10)<br>beta-galactosidase (3.2.1.23) (3.20.20.80)<br>neprilysin (3.4.24.11) (3.40.390.10) | 20.2 | 0 | 24.8 | 75 | 25 | 0 | ZN (13.68%) | | FMT (7.37%)<br>GPJ (5.26%)<br>BGC (5.26%)<br>G2F (5.26%)<br>RDF (5.26%)<br>BGP (4.21%)<br>GIM (3.16%)<br>GOX (3.16%)<br>GPF (2.11%)<br>BG6 (2.11%)<br>Other (33.61%) |
| **DE-DE-H-STY** | 1 | 6 | 2.1 | DNase (3.1.21.1) (3.60.10.10)<br>phosphomonoesterase (3.1.3.1) (3.40.720.10)<br>stearoyl-[acyl-carrier-protein] 9-desaturase (1.14.19.2) (1.10.620.20)<br>thermolysin (3.4.24.27) (1.10.390.10)<br>2 (5.4.2.12) (3.40.720.10)<br>SPT (2.3.1.50) (3.40.640.10) | 46.8 | 6.2 | 35.2 | 83.3 | 16.7 | 0 | ZN (19.82%)<br>FE (7.21%)<br>FE2 (5.41%)<br>MN (3.6%)<br>CO (1.8%) | PLP (6.31%) | LYS (10.81%)<br>PO4 (8.11%)<br>PLP (3.6%)<br>*P* (2.7%)<br>2PG (1.8%)<br>3PG (1.8%)<br>PAE (0.9%)<br>MMQ (0.9%)<br>AF3 (0.9%)<br>WO4 (0.9%)<br>Other (22.5%) |
| **DE-DE-H-X** | 1 | 2 | 2 | Unknown (3.2.-.-) (3.90.245.10)<br>phosphatidase (3.1.1.4) (1.20.90.10) | 16.5 | 0 | 20 | 100 | 0 | 0 | CA (20.0%) | | PIR (20.0%)<br>DNB (20.0%)<br>NOS (10.0%)<br>BDR (10.0%)<br>RIB (10.0%) |
| **DE-DE-K** | 2 | 12 | 1.4 | QAPRTase (2.4.2.19) (3.20.20.70)<br>methionine adenosyltransferase (2.5.1.6) (3.30.300.10)<br>(2E (2.5.1.10) (1.10.600.10)<br>lysine 2 (5.4.3.2) (3.20.20.70)<br>type II site-specific deoxyribonuclease (3.1.21.4) (3.40.91.10)<br>phosphoserine phosphatase (3.1.3.3) (3.40.50.1000)<br>isomeroreductase (1.1.1.86) (1.10.1040.10)<br>type II site-specific deoxyribonuclease (3.1.21.4) (3.40.210.10)<br>2-chloro-2 (5.5.1.7) (3.20.20.120)<br>nicotinamidase (3.5.1.19) (3.40.50.850)<br>OMP-DC (4.1.1.23) (3.20.20.70)<br>NADPH-hydroxymethylglutaryl-CoA reductase (1.1.1.34) (3.90.770.10) | 29 | 8.3 | 56.8 | 66.7 | 33.3 | 0 | MG (42.21%)<br>MN (4.02%)<br>CL (2.01%)<br>CA (1.01%)<br>K (0.5%)<br>NA (0.5%) | NAP (1.01%)<br>HMG (0.5%) | NCN (3.52%)<br>*P* (3.02%)<br>MSE (2.01%)<br>PO4 (2.01%)<br>HIO (2.01%)<br>MUC (2.01%)<br>TYR (2.01%)<br>40E (1.51%)<br>DGL (1.51%)<br>U5P (1.51%)<br>Other (23.59%) |
| **DE-DE-K-K-NQ** | 1 | 2 | 0.7 | mandelate racemase (5.1.2.2) (3.20.20.120)<br>glucarate dehydratase (4.2.1.40) (3.20.20.120) | 51 | 0 | 23 | 100 | 0 | 0 | MG (44.16%)<br>NA (2.6%)<br>CA (1.3%)<br>MN (1.3%) | | GLR (5.19%)<br>GKR (3.9%)<br>APG (2.6%)<br>DXL (2.6%)<br>LGT (2.6%)<br>XYH (2.6%)<br>ACT (1.3%)<br>BHO (1.3%)<br>0YR (1.3%)<br>3PY (1.3%)<br>Other (15.6%) |

| | | | | | | | | | | | | | |
|---|---|---|---|---|---|---|---|---|---|---|---|---|---|
| **DE-DE-K-NQ** | 1 | 4 | 1.6 | cerebroside-sulfatase (3.1.6.8) (3.40.720.10)<br>beta-methylaspartase (4.3.1.2) (3.20.20.120)<br>catechol O-methyltransferase (2.1.1.6) (3.40.50.150)<br>D-N-carbamoylase (3.5.1.77) (3.60.110.10) | 47.5 | 3.8 | 5 | 75 | 25 | 0 | MG (37.72%)<br>NA (3.51%)<br>CA (2.63%)<br>K (1.75%) | SAM (6.14%)<br>SAH (0.88%)<br>SFG (0.88%) | DNC (2.63%)<br>619 (2.63%)<br>FGP (1.75%)<br>FGL (0.88%)<br>ALS (0.88%)<br>2AS (0.88%)<br>72W (0.88%)<br>611 (0.88%)<br>76T (0.88%)<br>7JX (0.88%)<br>Other (33.44%) |
| **DE-DE-K-STY** | 1 | 2 | 1.6 | NADP+-ICDH (1.1.1.42) (3.40.718.10)<br>AvDH1 (3.6.4.12) (3.40.50.300) | 14.5 | 0 | 16.5 | 100 | 0 | 0 | MG (23.44%)<br>CA (10.94%)<br>ZN (3.12%)<br>MN (3.12%) | | ICT (17.19%)<br>IPM (17.19%)<br>SO4 (6.25%)<br>AKG (3.12%)<br>ICA (3.12%)<br>OXL (1.56%)<br>XYM (1.56%)<br>48Y (1.56%)<br>CIT (1.56%)<br>GOL (1.56%)<br>Other (1.56%) |
| **DE-DE-K-STY-STY-X** | 1 | 2 | 0.8 | SRR (5.1.1.18) (3.40.50.1100)<br>SRR (5.1.1.18) (3.40.50.1100) | 0 | 0 | 0 | 100 | 0 | 0 | | | |
| **DE-DE-NQ** | 1 | 12 | 1.7 | adenosylhomocysteinase (3.3.1.1) (3.40.50.1480)<br>exodeoxyribonuclease III (3.1.11.2) (3.60.10.10)<br>glucarate dehydratase (4.2.1.40) (3.20.20.120)<br>exodeoxyribonuclease III (3.1.11.2) (3.60.10.10)<br>kanamycin kinase (2.7.1.95) (3.90.1200.10)<br>SerRS (6.1.1.11) (3.30.930.10)<br>methanol dehydrogenase (1.1.2.7) (2.140.10.10)<br>biotin carboxylase (6.3.4.14) (3.30.470.20)<br>catechol O-methyltransferase (2.1.1.6) (3.40.50.150)<br>lysozyme (3.2.1.17) (1.10.530.10)<br>anhydrase (4.2.1.1) (2.160.10.10)<br>squalene---hopene cyclase (5.4.99.17) (1.50.10.20) | 28 | 5.8 | 21.9 | 66.7 | 33.3 | 0 | MG (28.94%)<br>CA (8.06%)<br>NA (1.1%)<br>CD (1.1%)<br>PR (0.73%)<br>RU (0.73%)<br>CE (0.37%)<br>EU (0.37%)<br>CO (0.37%)<br>MN (0.37%)<br>Other (0%) | PQQ (6.23%)<br>SAM (1.83%)<br>SAH (0.37%) | PQQ (3.66%)<br>KAN (2.93%)<br>GLR (1.47%)<br>ADP (1.47%)<br>GKR (1.1%)<br>ATP (1.1%)<br>DNC (1.1%)<br>619 (1.1%)<br>SO4 (1.1%)<br>BCT (1.1%)<br>Other (20.69%) |
| **DE-DE-NQ-NQ** | 1 | 2 | 1.9 | diisopropyl-fluorophosphatase (3.1.8.2) (2.120.10.30)<br>aryldialkylphosphatase (3.1.8.1) (2.120.10.30) | 100 | 0 | 0 | 100 | 0 | 0 | CA (69.57%) | | DOD (21.74%)<br>DI9 (4.35%) |
| **DE-DE-NQ-R** | 1 | 4 | 1.1 | glutathione synthase (6.3.2.3) (3.30.470.20)<br>ABL (2.7.10.2) (1.10.510.10)<br>endo-1 (3.2.1.78) (3.20.20.80)<br>D-alanine---D-alanine ligase (6.3.2.4) (3.30.470.20) | 48 | 0 | 49 | 75 | 25 | 0 | MG (37.78%) | | ADP (8.89%)<br>*P* (6.67%)<br>ANP (6.67%)<br>ACP (4.44%)<br>ATP (4.44%)<br>DAL (4.44%)<br>SO4 (2.22%)<br>N42 (2.22%)<br>2P5 (2.22%)<br>1Q4 (2.22%)<br>Other (17.76%) |
| **DE-DE-R** | 1 | 17 | 1.8 | TruB (5.4.99.25) (3.30.2350.10)<br>limonene-1 (3.3.2.8) (3.10.450.50)<br>mannonate dehydratase (4.2.1.8) (3.20.20.120)<br>xanthine oxidase (1.17.3.2) (3.30.365.10)<br>biotin carboxylase (6.3.4.14) (3.30.470.20)<br>pectinesterase (3.1.1.11) (2.160.20.10)<br>xanthine oxidase (1.17.3.2) (3.30.365.10)<br>Unknown (3.6.1.-) (3.90.79.10)<br>PEPPM (5.4.2.9) (3.20.20.60)<br>squalene---hopene cyclase (5.4.99.17) (1.50.10.20)<br>endo-1 (3.2.1.89) (3.20.20.80)<br>cholesterol oxidase (1.1.3.6) (3.40.462.10)<br>alpha-1 (4.2.2.2) (2.160.20.10)<br>mannonate dehydratase (4.2.1.8) (3.20.20.120)<br>thymidine kinase (2.7.1.21) (3.40.50.300)<br>5 (6.3.3.2) (3.40.50.10420)<br>OSBS (4.2.1.113) (3.20.20.120) | 21.2 | 0 | 27.2 | 47.1 | 47.1 | 5.9 | MG (21.6%)<br>ZN (0.8%)<br>NA (0.8%) | | URC (9.6%)<br>SAL (6.4%)<br>3ZS (3.2%)<br>CO3 (3.2%)<br>GCO (3.2%)<br>UNX (3.2%)<br>3ZQ (2.4%)<br>ADA (2.4%)<br>HPN (1.6%)<br>HYH (1.6%)<br>Other (28.0%) |

| | | | | | | | | | | | | | |
|---|---|---|---|---|---|---|---|---|---|---|---|---|---|
| **DE-DE-R-STY** | 1 | 3 | 2.2 | swivelase (5.6.2.1) (3.40.50.140)<br>exo-cellobiohydrolase (3.2.1.91) (3.20.20.40)<br>galactarate dehydratase (D-threo-forming) (4.2.1.158) (3.20.20.120) | 37.7 | 0 | 0 | 66.7 | 33.3 | 0 | MG (55.56%)<br>CA (11.11%) | | SSG (11.11%)<br>RCB (11.11%)<br>XYS (11.11%) |
| **DE-DE-STY** | 3 | 15 | 1.3 | dethiobiotin synthase (6.3.3.3) (3.40.50.300)<br>isopentenyl-diphosphate Delta-isomerase (5.3.3.2) (3.90.79.10)<br>EAS (4.2.3.61) (1.10.600.10)<br>6-phosphofructokinase (2.7.1.11) (3.40.50.450)<br>hyaluronoglucosaminidase (3.2.1.35) (3.20.20.70)<br>exodeoxyribonuclease III (3.1.11.2) (3.60.10.10)<br>endothiapepsin (3.4.23.22) (2.40.70.10)<br>steroid Delta-isomerase (5.3.3.1) (3.10.450.50)<br>anhydrosialidase (4.2.2.15) (2.120.10.10)<br>exo-alpha-sialidase (3.2.1.18) (2.120.10.10)<br>P450(eryF) (1.14.15.35) (1.10.630.10)<br>exo-alpha-sialidase (3.2.1.18) (2.120.10.10)<br>formyltetrahydrofolate dehydrogenase (1.5.1.6) (3.40.605.10)<br>tripeptidase (3.4.11.4) (1.10.390.10)<br>chitosanase (3.2.1.132) (1.20.141.10) | 10.1 | 5.1 | 28.9 | 33.3 | 66.7 | 0 | MG (3.89%)<br>MN (1.8%) | NDP (0.9%)<br>NAD (0.6%)<br>HEM (0.3%)<br>NAP (0.3%) | DAN (5.99%)<br>G39 (5.09%)<br>SIA (2.99%)<br>ZMR (2.69%)<br>BES (2.69%)<br>ADP (1.5%)<br>DPO (1.5%)<br>EQP (1.5%)<br>FSI (1.2%)<br>BCZ (1.2%)<br>Other (54.9%) |
| **DE-DE-STY-STY** | 1 | 4 | 1.8 | phosphomonoesterase (3.1.3.1) (3.40.720.10)<br>dTDP-glucose 4 (4.2.1.46) (3.40.50.720)<br>peroxidase (1.11.1.7) (1.10.640.10)<br>endo-1 (3.2.1.8) (2.60.120.180) | 50.5 | 9 | 16 | 50 | 50 | 0 | CA (59.21%)<br>MG (7.24%)<br>ZN (2.63%)<br>CO (0.66%) | NAD (3.29%) | DOD (8.55%)<br>XYP (2.63%)<br>X2F (1.97%)<br>TDX (1.32%)<br>GDD (1.32%)<br>DAU (1.32%)<br>DFX (0.66%)<br>C5X (0.66%)<br>07E (0.66%)<br>C3X (0.66%)<br>Other (0.66%) |
| **DE-DE-X** | 1 | 10 | 1.2 | methionine adenosyltransferase (2.5.1.6) (3.30.300.10)<br>Unknown (3.2.-.-) (3.90.245.10)<br>mGDH (1.1.5.2) (2.120.10.30)<br>phosphonatase (3.11.1.1) (3.40.50.1000)<br>fructose-bisphosphatase (3.1.3.11) (3.30.540.10)<br>acetolactate synthase (2.2.1.6) (3.40.50.970)<br>phosphoserine phosphatase (3.1.3.3) (3.40.50.1000)<br>Unknown (3.1.3.-) (3.40.50.1000)<br>type II site-specific deoxyribonuclease (3.1.21.4) (3.40.600.20)<br>aspartate---ammonia ligase (6.3.1.1) (3.30.930.10) | 86.7 | 14.3 | 45.4 | 100 | 0 | 0 | MG (34.02%)<br>CA (9.28%)<br>TL (7.73%)<br>ZN (4.64%)<br>MN (2.06%)<br>CL (1.55%)<br>K (1.03%)<br>NA (0.52%) | SAM (0.52%)<br>EN0 (0.52%)<br>5GY (0.52%)<br>TPP (0.52%) | PO4 (13.4%)<br>HE3 (1.55%)<br>2O2 (1.55%)<br>PIR (1.03%)<br>DNB (1.03%)<br>PI (1.03%)<br>PO3 (1.03%)<br>ALF (1.03%)<br>APO (1.03%)<br>PPK (0.52%)<br>Other (7.8%) |
| **DE-DE-X-X** | 1 | 2 | 2.4 | vesicle-fusing ATPase (3.6.4.6) (3.40.50.300)<br>nicotinamidase (3.5.1.19) (3.40.50.850) | 4 | 0 | 42.5 | 100 | 0 | 0 | MG (13.33%) | | ATP (40.0%)<br>ADP (13.33%)<br>NIO (13.33%)<br>ANP (6.67%)<br>VGL (6.67%)<br>CAF (6.67%) |
| **DE-F-K-STY** | 1 | 2 | 1.6 | kynureninase (3.7.1.3) (3.40.640.10)<br>L-lysine 6-transaminase (2.6.1.36) (3.40.640.10) | 0 | 83 | 1 | 0 | 100 | 0 | | PLP (71.11%)<br>PMP (2.22%) | RW2 (6.67%)<br>PLP (4.44%)<br>PFM (2.22%)<br>P00 (2.22%)<br>PXG (2.22%)<br>IF1 (2.22%)<br>PSZ (2.22%)<br>POI (2.22%)<br>PPE (2.22%) |
| **DE-F-VLI** | 1 | 3 | 1.9 | DHFR (1.5.1.3) (3.40.430.10)<br>pyruvate oxidase (1.2.3.3) (3.40.50.970)<br>histone acetyltransferase (2.3.1.48) (3.40.630.30) | 0 | 0 | 17 | 0 | 33.3 | 66.7 | | | TRR (40.14%)<br>MTX (16.9%)<br>TMQ (15.49%)<br>BDM (13.38%)<br>MMV (2.11%)<br>QKJ (2.11%)<br>FOL (1.41%)<br>TOP (1.41%)<br>WPD (1.41%)<br>WFA (0.7%)<br>Other (4.9%) |
| **DE-H-H** | 3 | 21 | 1.5 | beta-lactamase (3.5.2.6) (3.60.15.10)<br>alpha-toxin (3.1.4.3) (1.10.575.10)<br>phosphomonoesterase (3.1.3.2) (3.60.21.10)<br>L-fuculose-phosphate aldolase (4.1.2.17) (3.40.225.10)<br>luciferase (1.14.14.3) (3.20.20.30)<br>phenylalaninase (1.14.16.1) (1.10.800.10)<br>isopenicillin-N synthase (1.21.3.1) (2.60.120.330) | 61 | 1 | 31 | 71.4 | 28.6 | 0 | ZN (41.22%)<br>FE (6.62%)<br>FE2 (3.89%)<br>MN (1.47%)<br>CO (1.16%)<br>CD (0.95%)<br>NI (0.84%) | HBI (0.74%)<br>H4B (0.42%)<br>H2B (0.11%) | MRI (5.89%)<br>RTD (2.1%)<br>OGA (1.79%)<br>OH (0.84%)<br>PO4 (0.84%)<br>AKG (0.63%)<br>ZZ7 (0.53%) |

| | | | | | | | | | | | | |
|---|---|---|---|---|---|---|---|---|---|---|---|---|
| | | | | biotin carboxylase (6.3.4.14) (3.20.20.70)<br>mutarotase (5.1.3.3) (2.70.98.10)<br>[histone H3]-trimethyl-L-lysine36 demethylase (1.14.11.69) (2.60.120.650)<br>single-stranded-nucleate endonuclease (3.1.30.1) (1.10.575.10)<br>single-stranded-nucleate endonuclease (3.1.30.1) (1.10.575.10)<br>rhamnulose-1-phosphate aldolase (4.1.2.19) (3.40.225.10)<br>leucocyanidin (1.14.20.4) (2.60.120.330)<br>BphI (4.1.3.43) (3.20.20.70)<br>G6PDH (1.1.1.363) (3.30.360.10)<br>4a-hydroxytetrahydrobiopterin dehydratase (4.2.1.96) (3.30.1360.20)<br>methylglyoxal synthase (4.2.3.3) (3.40.50.1380)<br>phospholipase D (3.1.4.4) (3.30.870.10)<br>NDO (1.14.12.12) (3.90.380.10)<br>methylglyoxalase (4.4.1.5) (3.10.180.10) | | | | | | | CL (0.21%)<br>AU (0.21%)<br>MG (0.21%)<br>Other (0.11%) | | ACV (0.53%)<br>QUE (0.42%)<br>MYC (0.42%)<br>Other (21.48%) |
| **DE-H-H-H** | 3 | 21 | 1.7 | ToyB (4.1.2.50) (3.30.479.10)<br>anhydrase (4.2.1.1) (3.10.200.10)<br>glycerol dehydrogenase (1.1.1.6) (1.20.1090.10)<br>astacin (3.4.24.21) (3.40.390.10)<br>metalloelastase (3.4.24.65) (3.40.390.10)<br>glycoprotein gp63 (3.4.24.36) (3.10.170.20)<br>metalloelastase (3.4.24.65) (3.40.390.10)<br>rhamnulose-1-phosphate aldolase (4.1.2.19) (3.40.225.10)<br>oxalate oxidase (1.2.3.4) (2.60.120.10)<br>urease (3.5.1.5) (3.20.20.140)<br>beta-lactamase (3.5.2.6) (3.20.20.140)<br>Unknown (3.1.5.-) (1.10.3210.10)<br>2 (1.13.11.2) (3.10.180.10)<br>hydroxyacylglutathione hydrolase (3.1.2.6) (3.60.15.10)<br>homogentisicase (1.13.11.5) (2.60.120.10)<br>DD-carboxypeptidase (3.4.17.14) (3.30.1380.10)<br>protocatechuate 4 (1.13.11.8) (3.40.830.10)<br>beta-lactamase (3.5.2.6) (3.60.15.10)<br>SOD (1.15.1.1) (2.60.40.200)<br>adenosine deaminase (3.5.4.4) (3.20.20.140)<br>quercetinase (1.13.11.24) (2.60.120.10) | 79.7 | 2.2 | 20.9 | 90.5 | 4.8 | 0 | ZN (55.76%)<br>FE (3.15%)<br>CO (1.54%)<br>FE2 (1.35%)<br>CU (0.71%)<br>MN (0.71%)<br>NI (0.26%)<br>CD (0.26%)<br>MG (0.06%)<br>HG (0.06%)<br>Other (0.12%) | BIO (0.06%)<br>NAD (0.06%)<br>GBP (0.06%) | CGS (2.06%)<br>HSI (1.8%)<br>0DS (1.67%)<br>HAV (1.29%)<br>RTD (1.29%)<br>*P* (0.9%)<br>SO4 (0.77%)<br>R47 (0.45%)<br>NGH (0.39%)<br>GDE (0.39%)<br>Other (21.81%) |
| **DE-H-H-H-H-Z** | 1 | 2 | 0.6 | urease (3.5.1.5) (3.20.20.140)<br>Unknown (3.4.19.-) (3.20.20.140) | 88 | 0 | 11 | 100 | 0 | 0 | NI (41.27%)<br>ZN (11.64%)<br>F (2.65%)<br>MN (1.06%) | | KCX (25.93%)<br>OH (3.17%)<br>PO4 (2.65%)<br>2PA (2.12%)<br>HAE (1.59%)<br>BME (1.06%)<br>BO3 (0.53%)<br>DJM (0.53%)<br>URE (0.53%)<br>9XN (0.53%)<br>Other (4.24%) |
| **DE-H-H-H-NQ** | 1 | 2 | 1.9 | aldolase (4.1.2.13) (3.20.20.70)<br>oxalate oxidase (1.2.3.4) (2.60.120.10) | 100 | 0 | 100 | 100 | 0 | 0 | ZN (37.5%)<br>MN (12.5%) | | PGH (37.5%)<br>GLV (12.5%) |
| **DE-H-H-H-NQ-R** | 1 | 2 | 0.2 | calcineurin (3.1.3.16) (3.60.21.10)<br>calcineurin (3.1.3.16) (3.60.21.10) | 88 | 0 | 37 | 100 | 0 | 0 | MN (45.04%)<br>ZN (12.98%)<br>FE (9.16%)<br>MG (2.29%)<br>NI (2.29%)<br>CL (0.76%) | | PO4 (12.98%)<br>ENL (2.29%)<br>NHC (2.29%)<br>MES (1.53%)<br>SO4 (1.53%)<br>MLI (1.53%)<br>AGS (0.76%)<br>8D4 (0.76%)<br>NHE (0.76%)<br>4TF (0.76%)<br>Other (2.28%) |
| **DE-H-H-H-STY** | 1 | 2 | 1.9 | 2 (1.13.11.2) (3.10.180.10)<br>thermolysin (3.4.24.27) (3.10.170.10) | 93.5 | 0 | 31 | 100 | 0 | 0 | ZN (47.89%)<br>FE2 (6.1%)<br>FE (4.23%)<br>MN (3.29%)<br>CD (1.41%)<br>CO (0.94%)<br>CU (0.47%)<br>CU1 (0.47%)<br>CL (0.47%) | | VAL (1.88%)<br>DHY (1.41%)<br>LEU (1.41%)<br>NX6 (1.41%)<br>ASP (1.41%)<br>ACN (0.94%)<br>M3P (0.94%)<br>FEL (0.94%)<br>MCT (0.94%) |

| | | | | | | | | | | | | | |
|---|---|---|---|---|---|---|---|---|---|---|---|---|---|
| | | | | | | | | | | | RU (0.47%)<br>Other (0%) | | *P* (0.94%)<br>Other (19.27%) |
| **DE-H-H-NQ** | 1 | 2 | 2.3 | cytosine deaminase (3.5.4.1) (3.20.20.140)<br>aryldialkylphosphatase (3.1.8.1) (2.120.10.30) | 10 | 0 | 30 | 100 | 0 | 0 | MG (25.0%) | | FPY (25.0%)<br>O7U (16.67%)<br>1LD (8.33%)<br>IGA (8.33%)<br>HPY (8.33%)<br>17E (8.33%) |
| **DE-H-H-R** | 1 | 8 | 1.9 | phosphatidylinositol diacylglycerol-lyase (4.6.1.13) (3.20.20.190)<br>taurine dioxygenase (1.14.11.17) (3.60.130.10)<br>carboxypeptidase T (3.4.17.18) (3.40.630.10)<br>single-stranded-nucleate endonuclease (3.1.30.1) (1.10.575.10)<br>phosphomonoesterase (3.1.3.2) (3.40.50.1240)<br>oxalate decarboxylase (4.1.1.2) (2.60.120.10)<br>VanX (3.4.13.22) (3.30.1380.10)<br>bisphosphoglyceromutase (5.4.2.4) (3.40.50.1240) | 55.2 | 0.2 | 24.1 | 62.5 | 37.5 | 0 | ZN (33.65%)<br>FE2 (3.77%)<br>MN (2.83%)<br>FE (2.52%)<br>CO (2.2%)<br>MG (0.31%)<br>HG (0.31%) | DG2 (0.31%) | AKG (5.97%)<br>*P* (3.77%)<br>INS (2.52%)<br>3PG (2.2%)<br>VVO (1.26%)<br>CXA (1.26%)<br>SO4 (1.26%)<br>3K0 (0.94%)<br>VO4 (0.94%)<br>SIN (0.63%)<br>Other (23.96%) |
| **DE-H-H-STY** | 2 | 4 | 1.5 | Unknown (1.-.-.-) (3.60.15.10)<br>beta-hydroxyoctanoyl-ACP-dehydrase (4.2.1.59) (3.10.129.110)<br>[histone H3]-trimethyl-L-lysine36 demethylase (1.14.11.69) (2.60.120.650)<br>BphI (4.1.3.43) (3.20.20.70) | 30.8 | 0 | 42.8 | 75 | 25 | 0 | FE (3.64%)<br>MN (2.73%)<br>MG (1.82%)<br>ZN (0.91%) | | OGA (15.45%)<br>FEO (6.36%)<br>OXY (4.55%)<br>AKG (2.73%)<br>M3L (2.73%)<br>PYR (2.73%)<br>OXL (2.73%)<br>2HG (1.82%)<br>MMK (1.82%)<br>PD2 (1.82%)<br>Other (41.86%) |
| **DE-H-K** | 2 | 19 | 1.5 | methionine adenosyltransferase (2.5.1.6) (3.30.300.10)<br>PEPK (4.1.1.49) (3.90.228.20)<br>UlaD (4.1.1.85) (3.20.20.70)<br>Unknown (2.4.1.-) (3.90.550.10)<br>allyl-alcohol dehydrogenase (1.1.1.54) (3.20.20.100)<br>isochorismatase (3.3.2.1) (3.40.50.850)<br>phosphoenolpyruvate carboxykinase (GTP) (4.1.1.32) (3.40.449.10)<br>phosphomonoesterase (3.1.3.1) (3.40.720.10)<br>mandelate racemase (5.1.2.2) (3.20.20.120)<br>methylmalonyl-CoA mutase (5.4.99.2) (3.40.50.280)<br>SpeA (4.1.1.19) (3.20.20.10)<br>DHQase (4.2.1.10) (3.20.20.70)<br>(S)-oxynitrilase (4.1.2.47) (3.40.50.1820)<br>xylose isomerase (5.3.1.5) (3.20.20.150)<br>L-rhamnose isomerase (5.3.1.14) (3.20.20.150)<br>kasA (gene name) (2.3.1.293) (3.40.47.10)<br>SpeC (4.1.1.17) (3.40.640.10)<br>gliostatins (2.4.2.4) (3.40.1030.10)<br>neelaredoxin (1.15.1.2) (2.60.40.730) | 36.9 | 11.4 | 30.2 | 47.4 | 47.4 | 0 | MN (21.67%)<br>MG (11.74%)<br>ZN (3.16%)<br>CA (2.26%)<br>CO (1.81%)<br>FE (1.35%)<br>CL (0.23%)<br>NA (0.23%) | PLP (3.84%)<br>NAP (0.23%) | GTP (4.29%)<br>XYL (2.93%)<br>PO4 (2.71%)<br>GLC (2.48%)<br>ATP (2.03%)<br>OXL (2.03%)<br>PEP (1.81%)<br>SPV (1.81%)<br>EDO (1.35%)<br>PGA (1.13%)<br>Other (22.96%) |
| **DE-H-K-NQ** | 1 | 2 | 1.8 | triose-phosphate isomerase (5.3.1.1) (3.20.20.70)<br>phosphohexomutase (5.3.1.8) (2.60.120.10) | 50 | 0 | 11.5 | 50 | 50 | 0 | ZN (6.67%) | | PGA (26.67%)<br>PO4 (20.0%)<br>EDO (13.33%)<br>TPS (6.67%)<br>13P (6.67%)<br>PGH (6.67%)<br>129 (6.67%) |
| **DE-H-NQ** | 1 | 14 | 1.7 | steroid Delta-isomerase (5.3.3.1) (3.50.50.60)<br>6-phosphogluconic dehydrogenase (1.1.1.44) (1.10.1040.10)<br>Unknown (3.1.5.-) (1.10.3210.10)<br>barley nuclease (3.1.30.2) (3.40.570.10)<br>glucarate dehydratase (4.2.1.40) (3.20.20.120)<br>cytosine deaminase (3.5.4.1) (3.20.20.140)<br>peroxidase (1.11.1.7) (1.10.640.10)<br>peroxidase (1.11.1.7) (1.10.640.10)<br>oxalate oxidase (1.2.3.4) (2.60.120.10)<br>DHQase (4.2.1.10) (3.40.50.9100)<br>Unknown (2.3.1.-) (3.40.50.1220)<br>alpha-galactosyltransferase (2.4.1.87) (3.90.550.10)<br>glycine amidinotransferase (2.1.4.1) (3.75.10.10)<br>aminoacyl-tRNA hydrolase (3.1.1.29) (3.40.50.1470) | 8.6 | 18.8 | 27.1 | 42.9 | 57.1 | 0 | MG (2.69%)<br>K (2.39%)<br>IOD (0.6%)<br>NA (0.6%) | HEM (51.34%)<br>FAD (2.39%)<br>FDA (0.3%)<br>FAE (0.3%)<br>SFD (0.3%)<br>NDP (0.3%) | OSM (3.28%)<br>PEO (2.99%)<br>CYN (2.39%)<br>6PG (1.79%)<br>SHA (1.79%)<br>HEC (1.19%)<br>3TR (1.19%)<br>PZA (1.19%)<br>MMZ (1.19%)<br>OXY (0.9%)<br>Other (16.2%) |

| | | | | | | | | | | | | | |
|---|---|---|---|---|---|---|---|---|---|---|---|---|---|
| **DE-H-NQ-STY-X** | 1 | 4 | 1.1 | alkylacetyl-GPC:acetylhydrolase (3.1.1.47) (3.40.50.1110)<br>CA (3.5.1.93) (3.60.20.10)<br>RGAE (3.1.1.86) (3.40.50.1110)<br>palmitoyl-CoA hydrolase (3.1.2.2) (3.40.50.1110) | 0 | 0 | 0 | 0 | 100 | 0 | | | GD7 (100.0%) |
| **DE-H-NQ-X** | 1 | 2 | 1 | adenain (3.4.22.39) (3.40.395.10)<br>nucleoside-triphosphatase (3.6.1.15) (3.90.70.10) | 0 | 0 | 0 | 0 | 100 | 0 | | | E69 (100.0%) |
| **DE-H-R** | 2 | 17 | 1.5 | FRD (1.3.5.4) (3.90.700.10)<br>glycerophosphodiester phosphodiesterase (3.1.4.46) (3.20.20.190)<br>Xaa-Pro aminopeptidase (3.4.11.9) (3.90.230.10)<br>GS (6.3.1.2) (3.30.590.10)<br>Unknown (3.1.-.-) (3.90.540.10)<br>fructose-2 (3.1.3.46) (3.40.50.1240)<br>chloramphenicol O-acetyltransferase (2.3.1.28) (3.30.559.10)<br>mGDH (1.1.5.2) (2.120.10.30)<br>glycerophosphodiester phosphodiesterase (3.1.4.46) (3.20.20.190)<br>bromoperoxidase (1.11.1.18) (1.10.606.10)<br>bromoperoxidase (1.11.1.18) (1.10.606.10)<br>citrogenase (2.3.3.16) (1.10.230.10)<br>isocitrase (4.1.3.1) (3.20.20.60)<br>L(+)-nLDH (1.1.1.27) (3.90.110.10)<br>bontoxilysin (3.4.24.69) (3.90.1240.10)<br>barnase (4.6.1.24) (3.10.450.30)<br>PdxJ (2.6.99.2) (3.20.20.70) | 10.9 | 3.2 | 30.1 | 47.1 | 52.9 | 0 | MG (5.15%)<br>ZN (5.15%)<br>CA (1.47%)<br>MN (0.37%)<br>FE (0.37%) | NAI (2.21%)<br>NAD (2.21%)<br>CMC (1.47%)<br>CMX (0.74%)<br>ACO (0.74%)<br>HAX (0.37%)<br>SDX (0.37%)<br>FCX (0.37%)<br>COA (0.37%)<br>AMX (0.37%)<br>Other (2.59%) | *P* (8.09%)<br>OAA (5.51%)<br>P3S (2.94%)<br>PO4 (2.94%)<br>CIT (2.94%)<br>PRO (2.21%)<br>GOL (2.21%)<br>OXM (1.84%)<br>FUM (1.47%)<br>ADP (1.47%)<br>Other (32.13%) |
| **DE-H-R-R** | 1 | 4 | 1.4 | barnase (4.6.1.24) (3.10.450.30)<br>carboxy-cis (5.5.1.5) (2.130.10.10)<br>endo-alpha-sialidase (3.2.1.129) (2.120.10.10)<br>GS (6.3.1.2) (3.30.590.10) | 22.5 | 0 | 23 | 25 | 75 | 0 | MG (17.78%)<br>MN (2.22%) | | P3S (17.78%)<br>2GP (8.89%)<br>ADP (8.89%)<br>PO4 (8.89%)<br>3GP (6.67%)<br>SGP (4.44%)<br>*P* (2.22%)<br>ANP (2.22%) |
| **DE-H-R-STY** | 1 | 2 | 1.3 | Xaa-Pro aminopeptidase (3.4.11.9) (3.90.230.10)<br>hyaluronate lyase (4.2.2.1) (1.50.10.100) | 0 | 0 | 11.5 | 50 | 50 | 0 | | | PRO (26.09%)<br>*P* (17.39%)<br>M44 (8.7%)<br>GCT (8.7%)<br>IPA (4.35%)<br>MRD (4.35%)<br>PVC (4.35%)<br>ASG (4.35%)<br>IDR (4.35%) |
| **DE-H-STY** | 3 | 25 | 1.4 | 2 (1.3.1.34) (3.20.20.70)<br>trimethylamine dehydrogenase (1.5.8.2) (3.20.20.70)<br>5 (1.5.1.20) (3.20.20.220)<br>nucleoside-diphosphate kinase (2.7.4.6) (3.30.70.141)<br>destabilase (3.5.1.44) (3.40.50.180)<br>bromoperoxidase (1.11.1.18) (1.10.606.10)<br>3-mercaptopyruvate sulfurtransferase (2.8.1.2) (3.40.250.10)<br>acetylxylan esterase (3.1.1.72) (3.40.50.1820)<br>3 (4.1.99.12) (3.90.870.10)<br>Unknown (2.3.1.-) (3.40.50.1820)<br>endoprolylpeptidase (3.4.21.26) (3.40.50.1820)<br>AlaDH (1.4.1.1) (3.40.50.720)<br>BphI (4.1.3.43) (3.20.20.70)<br>peptidyl-Lys metalloendopeptidase (3.4.24.20) (3.40.390.10)<br>citrogenase (2.3.3.16) (1.10.580.10)<br>phosphatidase (3.1.1.4) (1.20.90.10)<br>scytalone dehydratase (4.2.1.94) (3.10.450.50)<br>D-arginine dehydrogenase (1.4.99.6) (3.50.50.60)<br>barnase (4.6.1.24) (3.10.450.30)<br>4HmO (1.1.3.46) (3.20.20.70)<br>ribulose-phosphate 3-epimerase (5.1.3.1) (3.20.20.70)<br>endo-1 (3.2.1.78) (3.20.20.80)<br>galactarate dehydratase (D-threo-forming) (4.2.1.158) (3.30.390.10)<br>nucleoside-triphosphatase (3.6.1.15) (2.40.10.10)<br>MetH (2.1.1.13) (3.40.50.280) | 11.2 | 9.2 | 14.6 | 28 | 60 | 12 | MG (7.6%)<br>ZN (2.28%)<br>MN (1.14%)<br>CA (1.14%)<br>FE2 (1.14%)<br>CD (0.38%)<br>NA (0.38%) | FMN (12.55%)<br>FNS (0.76%)<br>MDE (0.38%)<br>SDX (0.38%)<br>CIC (0.38%)<br>FNR (0.38%) | 2GP (11.41%)<br>FMN (10.65%)<br>OAA (4.94%)<br>3GP (3.8%)<br>CIT (2.66%)<br>SGP (1.9%)<br>LMR (1.52%)<br>MPD (1.52%)<br>2AM (1.52%)<br>PYR (1.14%)<br>Other (22.42%) |

| | | | | | | | | | | | | | |
|---|---|---|---|---|---|---|---|---|---|---|---|---|---|
| **DE-H-STY-STY-X** | 1 | 2 | 1.4 | Xaa-Pro-dipeptidyl-aminopeptidase (3.4.14.5) (3.40.50.1820)<br>CocE (3.1.1.84) (3.40.50.1820) | 0 | 0 | 1 | 0 | 100 | 0 | | | *P* (100.0%) |
| **DE-H-STY-X** | 2 | 5 | 0.8 | acetylcholinesterase (3.1.1.7) (3.40.50.1820)<br>narbonolide synthase (2.3.1.240) (3.40.50.1820)<br>DBP2 (3.6.4.13) (2.40.10.120)<br>chymotrypsin (3.4.21.1) (2.40.10.10)<br>coagulation factor XIa (3.4.21.27) (2.40.10.10) | 0 | 0 | 3 | 0 | 100 | 0 | | | 2ZF (21.21%)<br>*P* (15.15%)<br>2R9 (4.04%)<br>SV6 (4.04%)<br>1BV (3.03%)<br>0G6 (3.03%)<br>ELT (2.02%)<br>SUE (2.02%)<br>HTB (1.01%)<br>57H (1.01%)<br>Other (43.43%) |
| **DE-H-STY-X-X** | 1 | 11 | 1.3 | deamidase (3.4.16.5) (3.40.50.11320)<br>butyrinase (3.1.1.3) (3.40.50.1820)<br>prolyl aminopeptidase (3.4.11.5) (3.40.50.1820)<br>butyrinase (3.1.1.3) (3.40.50.1820)<br>palmitoyl[protein] hydrolase (3.1.2.22) (3.40.50.1820)<br>Unknown (3.1.2.-) (3.40.50.1820)<br>dipeptidase E (3.4.13.21) (3.40.50.880)<br>cutinase (3.1.1.74) (3.40.50.1820)<br>acetylxylan esterase (3.1.1.72) (3.40.50.1820)<br>ClpP (3.4.21.92) (3.90.226.10)<br>cholinesterase (3.1.1.8) (3.40.50.1820) | 0 | 0 | 0.7 | 0 | 100 | 0 | | | MUP (14.29%)<br>M3D (14.29%)<br>PLC (14.29%)<br>EMM (14.29%)<br>ELT (14.29%)<br>AT3 (14.29%)<br>NTJ (14.29%) |
| **DE-H-W** | 1 | 4 | 1.6 | stearoyl-[acyl-carrier-protein] 9-desaturase (1.14.19.2) (1.10.620.20)<br>L-ascorbate peroxidase (1.11.1.11) (1.10.420.10)<br>ribonuclease T2 (4.6.1.19) (3.90.730.10)<br>FGD1 (1.1.98.2) (3.20.20.30) | 0 | 10.2 | 0 | 50 | 50 | 0 | | HEM (72.6%)<br>ZNH (6.85%)<br>FMI (2.74%)<br>HEB (2.74%)<br>PP9 (1.37%)<br>DDH (1.37%) | DOD (5.48%)<br>522 (1.37%)<br>HEC (1.37%)<br>F42 (1.37%) |
| **DE-H-X** | 1 | 11 | 1.4 | 4-chlorobenzoyl-CoA dehalogenase (3.8.1.7) (3.90.226.10)<br>methylglyoxal synthase (4.2.3.3) (3.40.50.1380)<br>RNase (4.6.1.18) (3.10.130.10)<br>Unknown (4.6.1.-) (3.20.20.190)<br>acyl-[acyl-carrier-protein]---UDP-N-acetylglucosamine O-acyltransferase (2.3.1.129) (2.160.10.10)<br>transglutaminase (2.3.2.13) (3.90.260.10)<br>5-oxoprolyl-peptidase (3.4.19.3) (3.40.630.20)<br>C_overbar_42_ (3.4.21.43) (2.40.10.10)<br>peptidase K (3.4.21.64) (3.40.50.200)<br>CocE (3.1.1.84) (3.40.50.1820)<br>Unknown (3.1.-.-) (3.40.960.10) | 0.2 | 0 | 16.2 | 18.2 | 81.8 | 0 | CL (2.17%) | | *P* (19.57%)<br>DOD (6.52%)<br>PO4 (4.35%)<br>CGP (3.26%)<br>BCA (2.17%)<br>C3P (2.17%)<br>U3P (2.17%)<br>IMP (2.17%)<br>C2P (2.17%)<br>AMP (2.17%)<br>Other (28.3%) |
| **DE-H-X-X** | 2 | 6 | 1.3 | trypsin (3.4.21.4) (2.40.10.10)<br>nucleoside-triphosphatase (3.6.1.15) (2.40.10.10)<br>coagulation factor XIa (3.4.21.27) (2.40.10.10)<br>Lys-gingipain (3.4.22.47) (3.40.50.1460)<br>AtECH2 (4.2.1.119) (3.10.129.10)<br>beta-hydroxyoctanoyl-ACP-dehydrase (4.2.1.59) (3.10.129.110) | 2.3 | 0 | 4.2 | 0 | 100 | 0 | CL (1.06%) | | *P* (24.47%)<br>AG7 (8.51%)<br>0G6 (2.13%)<br>15U (2.13%)<br>DRX (2.13%)<br>5GI (2.13%)<br>G85 (2.13%)<br>G83 (2.13%)<br>K7J (1.06%)<br>BBL (1.06%)<br>Other (49.82%) |
| **DE-H-X-X-X** | 2 | 4 | 0.3 | alpha-lytic endopeptidase (3.4.21.12) (2.40.10.10)<br>alpha-lytic endopeptidase (3.4.21.12) (2.40.10.10)<br>trans-2 (5.3.3.14) (3.10.129.10)<br>beta-hydroxyoctanoyl-ACP-dehydrase (4.2.1.59) (3.10.129.10) | 13.2 | 0.8 | 5.8 | 0 | 100 | 0 | CL (31.37%) | PNS (1.96%) | *P* (13.73%)<br>B2F (7.84%)<br>AES (7.84%)<br>0EG (3.92%)<br>2A1 (3.92%)<br>B2V (3.92%)<br>GOL (1.96%)<br>C9H (1.96%)<br>3MQ (1.96%)<br>7SB (1.96%)<br>Other (9.8%) |
| **DE-K-K** | 2 | 11 | 1.3 | glutathione synthase (6.3.2.3) (3.30.470.20)<br>glutathione synthase (6.3.2.3) (3.30.470.20)<br>aldolase (4.1.2.13) (3.20.20.70)<br>glucocycloaldolase (5.5.1.4) (3.40.50.720)<br>phosphodeoxyriboaldolase (4.1.2.4) (3.20.20.70)<br>acetylglutamate kinase (2.7.2.8) (3.40.1160.10)<br>isopentenyl phosphate kinase (2.7.4.26) (3.40.1160.10) | 5.5 | 5.9 | 50.9 | 27.3 | 45.5 | 27.3 | MG (10.44%)<br>ZN (0.55%) | NAD (3.3%)<br>NAI (2.75%) | CAP (24.73%)<br>ADP (8.24%)<br>PO4 (4.4%)<br>RUB (3.3%)<br>SO4 (2.75%)<br>ACY (1.65%)<br>NLG (1.65%) |

| | | | | | | | | | | | | | |
|---|---|---|---|---|---|---|---|---|---|---|---|---|---|
| | | | | indole-3-glycerol-phosphate synthase (4.1.1.48) (3.20.20.70)<br>beta-lactamase (3.5.2.6) (3.40.710.10)<br>D-ribulose-1 (4.1.1.39) (3.20.20.110)<br>3'-phosphoadenylyl-sulfate:[heparan sulfate]-glucosamine 3-sulfotransferase (2.8.2.30) (3.40.50.300) | | | | | | | | | MSE (1.65%)<br>137 (1.65%)<br>13P (1.1%)<br>Other (24.2%) |
| **DE-K-NQ** | 2 | 7 | 1.3 | PEPPM (5.4.2.9) (3.20.20.60)<br>3-phosphoshikimate 1-carboxyvinyltransferase (2.5.1.19) (3.65.10.10)<br>PDHK (2.7.11.2) (3.30.565.10)<br>Dp38 (2.7.11.24) (1.10.510.10)<br>Unknown (5.5.1.-) (3.20.20.120)<br>cyclohydrolase (3.5.4.9) (3.40.50.10860)<br>selenocysteine lyase (4.4.1.16) (3.40.640.10) | 16 | 14.3 | 15 | 57.1 | 28.6 | 14.3 | MG (25.96%)<br>MN (2.88%) | PLP (7.69%) | ANP (9.62%)<br>FMT (5.77%)<br>GPJ (4.81%)<br>ATP (3.85%)<br>*P* (2.88%)<br>L34 (2.88%)<br>OXL (1.92%)<br>PO4 (1.92%)<br>GPF (1.92%)<br>ADP (1.92%)<br>Other (22.08%) |
| **DE-K-R** | 2 | 5 | 1.4 | 3-dehydroquinate-forming) (4.2.3.4) (1.20.1090.10)<br>pentalenene synthase (4.2.3.7) (1.10.600.10)<br>3'-deoxynucleotidase (3.1.3.34) (3.40.50.300)<br>endo-polygalacturonase (3.2.1.15) (2.160.20.10)<br>exodeoxyribonuclease (lambda-induced) (3.1.11.3) (1.10.150.20) | 0 | 0 | 18.6 | 40 | 40 | 20 | | | CRB (45.45%)<br>GTR (18.18%)<br>GLY (9.09%)<br>GLU (9.09%)<br>GTK (9.09%) |
| **DE-K-STY** | 2 | 13 | 1.7 | glutathione-disulfide reductase (1.8.1.7) (3.50.50.60)<br>NAD+-malic enzyme (1.1.1.38) (3.40.50.10380)<br>transaldolase (2.2.1.2) (3.20.20.70)<br>ingensin (3.4.25.1) (3.60.20.10)<br>glutamin-(asparagin-)ase (3.5.1.38) (3.40.50.1170)<br>NADP+-ICDH (1.1.1.42) (3.40.718.10)<br>dethiobiotin synthase (6.3.3.3) (3.40.50.300)<br>diphosphate---fructose-6-phosphate 1-phosphotransferase (2.7.1.90) (3.40.50.450)<br>AvDH1 (3.6.4.12) (3.40.50.300)<br>UDP-N-acetylmuramate---L-alanine ligase (6.3.2.8) (3.40.1190.10)<br>glycine hydroxymethyltransferase (2.1.2.1) (3.40.640.10)<br>glutamate-1-semialdehyde 2 (5.4.3.8) (3.40.640.10)<br>7 (2.6.1.62) (3.40.640.10) | 8.6 | 19.5 | 41.7 | 30.8 | 46.2 | 15.4 | MG (5.12%)<br>CA (1.97%)<br>MN (0.98%) | PLP (23.82%)<br>FAD (12.8%)<br>PMP (2.36%) | PLG (5.51%)<br>PLP (4.92%)<br>IPM (3.54%)<br>FAD (2.76%)<br>ASP (2.76%)<br>ICT (1.97%)<br>ADP (1.57%)<br>F6R (1.38%)<br>PMP (1.38%)<br>04C (1.18%)<br>Other (25.1%) |
| **DE-K-W** | 1 | 2 | 1 | PSAT (2.6.1.52) (3.40.640.10)<br>AAT (2.6.1.1) (3.40.640.10) | 0 | 74 | 0 | 0 | 100 | 0 | | PLP (60.18%)<br>PMP (7.08%)<br>PLR (0.88%)<br>NOP (0.88%)<br>MPL (0.88%) | PLP (5.31%)<br>PLA (5.31%)<br>PPD (1.77%)<br>PGU (1.77%)<br>IK2 (1.77%)<br>PMP (1.77%)<br>PP3 (1.77%)<br>PY6 (0.88%)<br>77E (0.88%)<br>PY5 (0.88%)<br>Other (3.52%) |
| **DE-K-X** | 1 | 3 | 1.8 | 2-chloro-2 (5.5.1.7) (3.20.20.120)<br>asparaginase (3.5.1.1) (3.40.50.1170)<br>exodeoxyribonuclease (lambda-induced) (3.1.11.3) (3.90.320.10) | 26.3 | 0 | 34 | 66.7 | 33.3 | 0 | CA (8.22%)<br>MG (4.11%)<br>CL (1.37%) | | ASP (9.59%)<br>DGL (8.22%)<br>TYR (5.48%)<br>NPQ (4.11%)<br>GLU (2.74%)<br>5CR (2.74%)<br>NLQ (2.74%)<br>MUC (2.74%)<br>AME (2.74%)<br>OSB (2.74%)<br>Other (15.07%) |
| **DE-NQ-NQ-STY** | 1 | 2 | 1.5 | lysozyme (3.2.1.17) (1.10.530.10)<br>meso-2 (4.2.1.28) (3.20.20.350) | 50 | 0 | 38 | 50 | 50 | 0 | K (40.91%)<br>CA (13.64%) | | PGO (31.82%)<br>*P* (4.55%)<br>GOL (4.55%)<br>PGR (4.55%) |
| **DE-NQ-R** | 1 | 6 | 1.6 | aspartate---ammonia ligase (6.3.1.1) (3.30.930.10)<br>adenylylcyclase (4.6.1.1) (1.10.400.10)<br>acetokinase (2.7.2.1) (3.30.420.40)<br>A-kinase (2.7.11.1) (3.30.565.10)<br>MICL (4.1.3.30) (3.20.20.60)<br>AvDH1 (3.6.4.12) (3.40.50.300) | 10 | 0 | 47.2 | 33.3 | 50 | 16.7 | MG (3.03%) | | GSP (26.26%)<br>GDP (26.26%)<br>ALF (26.26%)<br>ASN (4.04%)<br>ATP (4.04%)<br>ADP (3.03%)<br>AMP (2.02%)<br>B4P (2.02%)<br>SIN (1.01%)<br>APC (1.01%)<br>Other (1.01%) |

| | | | | | | | | | | | | | |
|---|---|---|---|---|---|---|---|---|---|---|---|---|---|
| **DE-NQ-STY** | 1 | 3 | 1.4 | L-DEX (3.8.1.2) (3.40.50.1000)<br>beta-glucosidase (3.2.1.21) (3.20.20.80)<br>beta-galactosidase (3.2.1.23) (3.20.20.80) | 0 | 0 | 27.3 | 0 | 100 | 0 | | | BGC (8.05%)<br>BG6 (5.75%)<br>G2F (5.75%)<br>BGP (4.6%)<br>GIM (3.45%)<br>GOX (3.45%)<br>GLC (2.3%)<br>S55 (2.3%)<br>GTL (2.3%)<br>IFM (2.3%)<br>Other (28.75%) |
| **DE-NQ-W** | 1 | 2 | 1.9 | photolyase (4.1.99.3) (1.25.40.80)<br>2-hydroxychromene-2-carboxylate isomerase (5.99.1.4) (3.40.30.10) | 0 | 41 | 3 | 0 | 50 | 50 | | FAD (66.67%) | FAD (14.29%)<br>TCP (4.76%)<br>*P* (4.76%)<br>TDR (4.76%)<br>GSH (4.76%) |
| **DE-NQ-X** | 2 | 10 | 1.4 | formyl-CoA transferase (2.8.3.16) (3.40.50.10540)<br>Delta3-Delta2-enoyl-CoA isomerase (5.3.3.8) (3.90.226.10)<br>glutathione synthase (6.3.2.3) (3.30.1490.50)<br>Delta3-Delta2-enoyl-CoA isomerase (5.3.3.8) (3.90.226.10)<br>indolepyruvate decarboxylase (4.1.1.74) (3.40.50.970)<br>CEAS (2.5.1.66) (3.40.50.970)<br>isoamylase (3.2.1.68) (3.20.20.80)<br>pantothenate synthetase (6.3.2.1) (3.40.50.620)<br>adenylosuccinate synthase (6.3.4.4) (3.40.440.10)<br>ErmC' (2.1.1.184) (3.40.50.150) | 41.1 | 37.4 | 18.7 | 60 | 30 | 10 | MG (23.29%)<br>CA (0.68%) | TPP (13.01%)<br>COA (2.74%)<br>CO8 (1.37%)<br>SAM (1.37%)<br>CAA (0.68%)<br>3HC (0.68%)<br>3H9 (0.68%)<br>GSH (0.68%)<br>TDL (0.68%)<br>TDP (0.68%)<br>Other (2.04%) | GDP (7.53%)<br>IMP (6.16%)<br>IMO (4.11%)<br>SO4 (3.42%)<br>HDA (3.42%)<br>PO4 (2.74%)<br>NO3 (2.74%)<br>AMP (2.05%)<br>ADP (1.37%)<br>ACT (1.37%)<br>Other (12.26%) |
| **DE-R-R** | 3 | 10 | 1.5 | thermonuclease (3.1.31.1) (2.40.50.90)<br>glutathione synthase (6.3.2.3) (3.30.470.20)<br>M.BsuRIa (2.1.1.37) (3.40.50.150)<br>duplicase (2.7.7.7) (2.30.40.20)<br>E1 (6.2.1.45) (3.50.50.80)<br>E1 (6.2.1.45) (3.50.50.80)<br>TruA (5.4.99.12) (3.30.70.660)<br>SerRS (6.1.1.11) (3.30.930.10)<br>7 (4.1.2.25) (3.30.70.560)<br>thymidine kinase (2.7.1.21) (3.40.50.300) | 12.3 | 0 | 56.7 | 50 | 10 | 40 | CA (2.4%)<br>MG (1.6%) | | THP (72.8%)<br>SO4 (7.2%)<br>*P* (4.8%)<br>PO4 (2.4%)<br>C49 (2.4%)<br>ATP (1.6%)<br>C36 (0.8%)<br>C37 (0.8%)<br>PYO (0.8%)<br>DTP (0.8%)<br>Other (1.6%) |
| **DE-R-STY** | 4 | 16 | 1.6 | PTR1 (1.5.1.33) (3.40.50.720)<br>hexokinase (2.7.1.1) (3.30.420.40)<br>hexokinase (2.7.1.1) (3.30.420.40)<br>trans (4.2.3.6) (1.10.600.10)<br>NAD+ glycohydrolase (3.2.2.5) (3.90.176.10)<br>D-camphor-exo-hydroxylase (1.14.15.1) (1.10.630.10)<br>uridine-5'-diphospho-N-acetyl-2-amino-2-deoxy-3-O-lactylglucose:NADP-oxidoreductase (1.3.1.98) (3.90.78.10)<br>3-hydroxybenzoate 4-monooxygenase (1.14.13.23) (3.30.9.10)<br>SerRS (6.1.1.11) (3.30.930.10)<br>gliostatins (2.4.2.4) (3.40.1030.10)<br>biliverdin reductase (1.3.1.24) (3.30.360.10)<br>exo-alpha-sialidase (3.2.1.18) (2.120.10.10)<br>Unknown (2.4.2.-) (3.90.210.10)<br>UGM (5.4.99.9) (3.40.50.720)<br>uroporphyrinogen decarboxylase (4.1.1.37) (3.20.20.210)<br>aspartate---tRNA ligase (6.1.1.12) (3.30.930.10) | 0.7 | 16.2 | 16.7 | 18.8 | 56.2 | 25 | K (0.3%) | NAP (15.41%)<br>FAD (2.72%)<br>NDP (2.42%)<br>HBI (0.6%)<br>H4B (0.3%)<br>TAD (0.3%) | NAP (9.97%)<br>G39 (4.23%)<br>ZMR (2.42%)<br>SSA (2.11%)<br>SIA (2.11%)<br>DAN (1.81%)<br>MTX (1.51%)<br>G6P (1.21%)<br>AMP (1.21%)<br>BCZ (1.21%)<br>Other (46.56%) |
| **DE-R-X** | 1 | 2 | 1.4 | phosphohexomutase (5.3.1.9) (3.40.50.10490)<br>Unknown (6.3.1.21) (3.30.470.20) | 0 | 0 | 9 | 50 | 50 | 0 | | | 6PG (16.67%)<br>F6P (16.67%)<br>PA5 (10.0%)<br>E4P (10.0%)<br>S6P (10.0%)<br>A5P (6.67%)<br>DER (6.67%)<br>RI2 (3.33%)<br>G6Q (3.33%)<br>G6P (3.33%)<br>Other (3.33%) |
| **DE-STY-STY** | 3 | 11 | 1.7 | phosphatidase (3.1.1.4) (1.20.90.10)<br>ArsC (ambiguous) (1.20.4.4) (3.40.50.2300)<br>ingensin (3.4.25.1) (3.60.20.10)<br>dTDP-glucose 4 (4.2.1.46) (3.40.50.720)<br>cytochrome P450nor (1.7.1.14) (1.10.630.10)<br>D-alanine---D-alanine ligase (6.3.2.4) (3.30.1490.20)<br>Transferred to 5.6.1.7 (3.6.4.9) (1.10.560.10)<br>PARP (2.4.2.30) (3.90.228.10) | 4.7 | 1.6 | 34.2 | 0 | 54.5 | 36.4 | K (0.77%)<br>NA (0.38%)<br>MG (0.38%) | NAD (1.54%) | *P* (28.08%)<br>ASP (8.08%)<br>ADP (4.23%)<br>DAL (2.69%)<br>CNQ (1.54%)<br>2YQ (1.54%)<br>09L (1.54%)<br>78P (1.54%) |

| | | | | | | | | | | | | | |
|---|---|---|---|---|---|---|---|---|---|---|---|---|---|
| | | | | indophenolase (7.1.1.9) (1.20.210.10)<br>glutamin-(asparagin-)ase (3.5.1.38) (3.40.50.1170)<br>asparaginase (3.5.1.1) (3.40.50.1170) | | | | | | | | | 3BV (1.15%)<br>POL (1.15%)<br>Other (43.09%) |
| **DE-STY-W** | 1 | 2 | 1.3 | thymidylate synthase (2.1.1.45) (3.30.572.10)<br>AlkA (3.2.2.21) (1.10.340.30) | 0 | 2 | 15 | 0 | 50 | 50 | | C2F (7.32%)<br>THG (2.44%) | UMP (41.46%)<br>UFP (19.51%)<br>UMC (4.88%)<br>NOH (4.88%)<br>NDU (2.44%)<br>CB3 (2.44%)<br>VLD (2.44%)<br>FGT (2.44%)<br>CF9 (2.44%) |
| **DE-STY-X** | 1 | 4 | 1.7 | DNA[cytosine-N4]methyltransferase (2.1.1.113) (3.40.50.150)<br>phosphatidase (3.1.1.4) (3.40.1090.10)<br>limonene-1 (3.3.2.8) (3.10.450.50)<br>cis-aconitase (4.2.1.3) (3.20.19.10) | 0 | 0 | 18.2 | 0 | 100 | 0 | | | 3ZS (18.52%)<br>3ZQ (11.11%)<br>HPN (7.41%)<br>HYH (7.41%)<br>6VV (3.7%)<br>VPR (3.7%)<br>40O (3.7%)<br>ATH (3.7%)<br>MIC (3.7%)<br>FLC (3.7%)<br>Other (18.5%) |
| **DE-STY-X-X** | 1 | 2 | 1.8 | 1 (4.1.3.36) (3.90.226.10)<br>arsenite-transporting ATPase (7.3.2.7) (3.40.50.300) | 12.5 | 37.5 | 50 | 0 | 0 | 100 | MG (12.5%) | 1HA (25.0%)<br>2NE (12.5%) | ADP (50.0%) |
| **DE-STY-Z** | 1 | 2 | 0.5 | L-cysteine desulfidase (4.4.1.28) (3.40.640.10)<br>homocysteine desulfhydrase (4.4.1.2) (3.40.640.10) | 0 | 19 | 0 | 0 | 100 | 0 | | PLP (18.8%) | LLP (35.04%)<br>PLP (6.84%)<br>4LM (6.84%)<br>3LM (5.13%)<br>2LM (3.42%)<br>5OW (3.42%)<br>HEN (1.71%)<br>PPJ (1.71%)<br>KOU (1.71%)<br>LPI (1.71%)<br>Other (13.68%) |
| **DE-X-X** | 2 | 9 | 1.4 | beta-lactamase (3.5.2.6) (3.40.710.10)<br>Delta3-Delta2-enoyl-CoA isomerase (5.3.3.8) (3.90.226.10)<br>Transferred to 4.1.2.61 (4.2.1.101) (3.90.226.10)<br>triose-phosphate isomerase (5.3.1.1) (3.20.20.70)<br>Transferred to 5.6.1.3 (3.6.4.4) (3.40.850.10)<br>Unknown (5.3.3.-) (3.90.226.10)<br>crotonase (4.2.1.17) (3.90.226.40)<br>isovaleryl-CoA dehydrogenase (1.3.8.4) (1.20.140.10)<br>ArsC (ambiguous) (1.20.4.4) (3.40.50.2300) | 0.1 | 5.7 | 26.9 | 0 | 66.7 | 22.2 | K (0.84%) | ACO (4.2%)<br>CO8 (1.68%) | NXL (5.04%)<br>DOD (5.04%)<br>PGA (5.04%)<br>SO4 (4.2%)<br>PO4 (3.36%)<br>CB4 (2.52%)<br>ISS (2.52%)<br>2UL (1.68%)<br>C8V (1.68%)<br>IM2 (1.68%)<br>Other (50.4%) |
| **DE-X-X-X** | 1 | 3 | 1.9 | phosphatidase (3.1.1.4) (1.20.90.10)<br>ribokinase (2.7.1.15) (3.40.1190.20)<br>ketohexokinase (2.7.1.3) (3.40.1190.20) | 16.7 | 0 | 21.7 | 33.3 | 66.7 | 0 | CA (28.18%)<br>NA (1.36%)<br>CL (0.91%)<br>CD (0.45%) | | ADN (14.55%)<br>ADP (12.73%)<br>ATP (5.0%)<br>ACP (4.55%)<br>ANP (3.18%)<br>KDG (1.82%)<br>FRU (1.36%)<br>ACY (0.91%)<br>U8D (0.91%)<br>GEL (0.91%)<br>Other (15.79%) |
| **F-NQ-VLI** | 1 | 2 | 0.9 | pyruvate oxidase (1.2.3.3) (3.40.50.970)<br>pyruvate:ubiquinone-8-oxidoreductase (1.2.5.1) (3.40.50.970) | 0 | 25 | 0 | 0 | 0 | 100 | | TDP (100.0%) | |
| **H-H-H** | 1 | 3 | 1 | HGAD (4.3.2.5) (2.60.120.310)<br>SOD (1.15.1.1) (2.60.40.200)<br>isopentenyl-diphosphate Delta-isomerase (5.3.3.2) (3.90.79.10) | 85.3 | 0 | 0.3 | 100 | 0 | 0 | CU (41.62%)<br>CU1 (27.17%)<br>ZN (10.98%)<br>MN (5.78%)<br>CD (0.58%) | | AZI (1.16%)<br>CO3 (0.58%)<br>PEO (0.58%) |

| | | | | | | | | | | | | | |
|---|---|---|---|---|---|---|---|---|---|---|---|---|---|
| **H-H-H-H** | 1 | 3 | 1.9 | phosphomonoesterase (3.1.3.2) (3.60.21.10)<br>laccase (1.10.3.2) (2.60.40.420)<br>laccase (1.10.3.2) (2.60.40.420) | 69 | 0 | 50 | 100 | 0 | 0 | CU (82.63%)<br>NA (0.28%)<br>F (0.28%) | | OXY (10.36%)<br>PER (2.8%)<br>SO4 (0.84%)<br>PO4 (0.28%)<br>0K7 (0.28%)<br>WO4 (0.28%)<br>AD9 (0.28%)<br>0LV (0.28%)<br>H1T (0.28%)<br>ANP (0.28%)<br>Other (0%) |
| **H-H-H-NQ** | 1 | 4 | 1.6 | 3-dehydroquinate-forming) (4.2.3.4) (1.20.1090.10)<br>aldolase (4.1.2.13) (3.20.20.70)<br>anhydrase (4.2.1.1) (2.160.10.10)<br>13-lipoxidase (1.13.11.12) (1.20.245.10) | 78.5 | 0 | 59.5 | 75 | 0 | 25 | FE2 (29.79%)<br>FE (24.47%)<br>ZN (13.83%)<br>CO (3.19%)<br>MN (3.19%) | | CRB (5.32%)<br>BCT (4.26%)<br>PGH (3.19%)<br>SO4 (3.19%)<br>PO4 (1.06%)<br>GLU (1.06%)<br>9OH (1.06%)<br>13S (1.06%)<br>13R (1.06%)<br>EGT (1.06%)<br>Other (1.06%) |
| **H-H-H-STY** | 1 | 3 | 1.7 | indophenolase (7.1.1.9) (1.20.210.10)<br>beta-lactamase (3.5.2.6) (3.60.15.10)<br>primary-amine oxidase (1.4.3.21) (2.70.98.20) | 92.3 | 31.3 | 41.7 | 100 | 0 | 0 | CU (38.23%)<br>ZN (8.03%)<br>CU1 (1.39%)<br>CO (1.39%)<br>NI (0.83%)<br>O (0.28%)<br>CD (0.28%)<br>CL (0.28%) | HEA (16.9%) | TYQ (6.65%)<br>AZI (3.6%)<br>TPQ (3.6%)<br>PER (3.32%)<br>CMO (2.49%)<br>PEO (1.66%)<br>OH (1.11%)<br>HEA (1.11%)<br>MX1 (1.11%)<br>NO (0.83%)<br>Other (5.59%) |
| **H-H-K** | 1 | 2 | 0.7 | RNase (4.6.1.18) (3.10.130.10)<br>Unknown (3.1.27.-) (3.10.130.10) | 3 | 1 | 12.5 | 0 | 100 | 0 | CL (2.59%)<br>ZN (0.86%)<br>PT (0.86%)<br>AU (0.86%) | NDP (0.86%)<br>NAP (0.86%) | *P* (4.31%)<br>U2G (2.59%)<br>ADT (2.59%)<br>U5P (2.59%)<br>AMP (2.59%)<br>POP (1.72%)<br>ATR (1.72%)<br>SFB (1.72%)<br>U5F (1.72%)<br>UMF (1.72%)<br>Other (27.52%) |
| **H-H-NQ** | 2 | 5 | 1.3 | bis(5'-adenosyl)-triphosphatase (3.6.1.29) (3.30.428.10)<br>UDP-sugar diphosphatase (3.6.1.45) (3.60.21.10)<br>aryldialkylphosphatase (3.1.8.1) (2.120.10.30)<br>beta-lactamase (3.5.2.6) (3.60.15.10)<br>Transferred to 2.7.1.191 and 2.7.1.192 and 2.7.1.193 and<br>2.7.1.194 and 2.7.1.195 and 2.7.1.196 and 2.7.1.197 and<br>2.7.1.198 and 2.7.1.199 and 2.7.1.200 and 2.7.1.201 and<br>2.7.1.202 and 2.7.1.203 and 2.7.1.204 and 2.7.1.205 and<br>2.7.1.206 and 2.7.1.207 and 2.7.1.208 (2.7.1.69) (1.20.58.80) | 24.2 | 0 | 12.4 | 80 | 20 | 0 | ZN (37.02%)<br>CD (1.1%)<br>CL (0.55%) | | RTD (10.5%)<br>SO4 (2.21%)<br>ZZ7 (2.21%)<br>AMP (1.1%)<br>PO4 (1.1%)<br>CO3 (1.1%)<br>BCT (1.1%)<br>QT2 (1.1%)<br>X8Z (1.1%)<br>S3C (1.1%)<br>Other (28.05%) |
| **H-H-R** | 2 | 7 | 1.4 | urease (3.5.1.5) (3.20.20.140)<br>phosphomonoesterase (3.1.3.2) (1.20.144.10)<br>NadB (1.4.3.16) (3.50.50.60)<br>SOD (1.15.1.1) (2.60.40.200)<br>Unknown (3.1.-.-) (3.90.540.10)<br>L-ascorbate peroxidase (1.11.1.11) (1.10.420.10)<br>UDP-N-acetyl-D-glucosamine 2-epimerase (5.1.3.14)<br>(3.40.50.2000) | 15.3 | 13 | 27.9 | 57.1 | 42.9 | 0 | CU (15.64%)<br>CU1 (13.13%)<br>ZN (4.47%)<br>NI (3.35%)<br>O (0.56%)<br>MN (0.28%)<br>NA (0.28%)<br>F (0.28%) | HEM (28.49%)<br>ZNH (1.12%)<br>HEB (0.84%)<br>FMI (0.56%)<br>PP9 (0.28%)<br>DDH (0.28%) | TOX (3.91%)<br>SO4 (2.79%)<br>DOD (1.96%)<br>OXY (1.68%)<br>HEM (1.4%)<br>2PA (1.12%)<br>HAE (0.84%)<br>PO4 (0.84%)<br>CO3 (0.56%)<br>MOO (0.56%)<br>Other (5.88%) |
| **H-H-STY** | 1 | 11 | 1.4 | NO-forming) (1.7.2.1) (2.60.40.420)<br>assemblin (3.4.21.97) (3.20.16.10)<br>galactose oxidase (1.1.3.9) (2.130.10.80)<br>astacin (3.4.24.21) (3.40.390.10)<br>OYE (1.6.99.1) (3.20.20.70)<br>nicotinamidase (3.5.1.19) (3.40.50.850)<br>1 (1.13.11.1) (2.60.130.10)<br>redoxyendonuclease (3.1.21.2) (3.20.20.150) | 40.9 | 3.4 | 32.7 | 54.5 | 36.4 | 9.1 | FE (11.34%)<br>ZN (7.14%)<br>CU (6.72%)<br>HG (0.42%)<br>CO (0.42%)<br>NI (0.42%) | FMN (13.03%)<br>FNR (0.42%) | FMN (5.04%)<br>SO4 (5.04%)<br>*P* (3.36%)<br>ACT (2.94%)<br>CAQ (2.1%)<br>FNR (1.68%)<br>COU (1.68%)<br>0FP (1.26%) |

| | | | | | | | | | | | | | |
|---|---|---|---|---|---|---|---|---|---|---|---|---|---|
| | | | | tyrosine---tRNA ligase (6.1.1.1) (3.40.50.620)<br>ATP-sulfurylase (2.7.7.4) (3.40.50.620)<br>EIIGlc (2.7.1.199) (2.70.70.10) | | | | | | | | | HBA (1.26%)<br>DHB (1.26%)<br>Other (26.46%) |
| **H-H-Z** | 1 | 2 | 1.4 | urease (3.5.1.5) (3.20.20.140)<br>aryldialkylphosphatase (3.1.8.1) (3.20.20.140) | 95.5 | 0 | 24 | 100 | 0 | 0 | NI (35.14%)<br>ZN (9.46%)<br>CO (3.15%)<br>F (2.25%)<br>CD (2.25%)<br>MN (1.8%) | | KCX (23.87%)<br>FMT (3.15%)<br>OH (2.7%)<br>PO4 (2.25%)<br>2PA (1.8%)<br>HAE (1.35%)<br>BME (0.9%)<br>BO3 (0.45%)<br>SO4 (0.45%)<br>DJM (0.45%)<br>Other (4.05%) |
| **H-K-NQ** | 1 | 2 | 1.8 | nucleoside-diphosphate kinase (2.7.4.6) (3.30.70.141)<br>SPT (2.3.1.50) (3.40.640.10) | 6.5 | 18.5 | 28 | 0 | 100 | 0 | MG (4.65%) | PLP (8.14%) | ADP (15.12%)<br>CDP (5.81%)<br>UDP (4.65%)<br>TYD (4.65%)<br>GDP (4.65%)<br>PLP (4.65%)<br>DGI (2.33%)<br>AMP (2.33%)<br>PCG (2.33%)<br>3AN (2.33%)<br>Other (22.04%) |
| **H-K-STY** | 1 | 10 | 1.3 | gliostatins (2.4.2.4) (3.40.1030.10)<br>TDP-4-keto-L-rhamnose-3 (5.1.3.13) (2.60.120.10)<br>tRNA-intron lyase (4.6.1.16) (3.40.1350.10)<br>dehydrogenase (1.1.2.3) (3.20.20.70)<br>uricase (1.7.3.3) (3.10.270.10)<br>DD-peptidase (3.4.16.4) (3.40.710.10)<br>allyl-alcohol dehydrogenase (1.1.1.54) (3.20.20.100)<br>6-phosphogluconic dehydrogenase (1.1.1.44) (3.40.50.720)<br>cerebroside-sulfatase (3.1.6.8) (3.40.720.10)<br>UDP-sulfoquinovose synthase (3.13.1.1) (3.40.50.720) | 1.6 | 16.8 | 28.9 | 0 | 100 | 0 | CL (3.31%)<br>K (0.25%) | NAP (16.28%)<br>FMN (7.63%)<br>NDP (2.54%)<br>FNS (0.51%)<br>NAD (0.51%)<br>FNR (0.25%) | AZA (11.7%)<br>DOD (3.31%)<br>NAP (2.54%)<br>OXY (2.29%)<br>URC (2.04%)<br>ZST (1.78%)<br>LDT (1.78%)<br>FID (1.53%)<br>6PG (1.53%)<br>AZI (1.27%)<br>Other (32.51%) |
| **H-K-STY-STY** | 1 | 2 | 2.5 | dehydrogenase (1.1.2.3) (3.20.20.70)<br>UDP-sulfoquinovose synthase (3.13.1.1) (3.40.50.720) | 0 | 43.5 | 50 | 0 | 100 | 0 | | FMN (73.17%)<br>FNS (4.88%)<br>FNR (2.44%) | FMN (9.76%)<br>UPG (4.88%) |
| **H-K-X** | 1 | 2 | 1.5 | adenylosuccinate synthase (6.3.4.4) (3.40.440.10)<br>RNase (4.6.1.18) (3.10.130.10) | 38 | 0 | 58 | 50 | 50 | 0 | MG (11.98%)<br>CL (5.39%)<br>NA (0.6%)<br>IR (0.6%) | | GDP (11.98%)<br>DOD (7.19%)<br>IMP (4.79%)<br>PO4 (4.19%)<br>IMO (3.59%)<br>CGP (2.99%)<br>NO3 (2.4%)<br>AMP (1.8%)<br>SO4 (1.8%)<br>HDA (1.8%)<br>Other (23.4%) |
| **H-NQ-NQ-X** | 1 | 4 | 0.3 | papain (3.4.22.2) (3.90.70.10)<br>bleomycin hydrolase (3.4.22.40) (3.90.70.10)<br>Unknown (3.4.22.-) (3.90.70.10)<br>cathepsin K (3.4.22.38) (3.90.70.10) | 0 | 0 | 0.2 | 0 | 100 | 0 | | | 074 (13.64%)<br>D1R (9.09%)<br>2VC (9.09%)<br>MYP (9.09%)<br>9U8 (9.09%)<br>VS4 (9.09%)<br>*P* (4.55%)<br>TCK (4.55%)<br>0IW (4.55%)<br>BCQ (4.55%)<br>Other (18.2%) |
| **H-NQ-R** | 1 | 4 | 1.3 | OTC (2.1.3.3) (3.40.50.1370)<br>peroxidase (1.11.1.7) (1.10.520.10)<br>aspartate-semialdehyde dehydrogenase (1.2.1.11) (3.30.360.10)<br>N-FDH (1.17.1.9) (3.40.50.720) | 0 | 0 | 16.5 | 0 | 50 | 50 | | | CP (21.95%)<br>PAO (12.2%)<br>PO4 (9.76%)<br>NVA (7.32%)<br>FER (7.32%)<br>BHO (7.32%)<br>CYS (7.32%)<br>SO4 (4.88%) |

| | | | | | | | | | | | | |
|---|---|---|---|---|---|---|---|---|---|---|---|---|
| | | | | | | | | | | | | ACT (4.88%)<br>CIR (2.44%)<br>Other (9.76%) |
| **H-NQ-STY** | 1 | 7 | 1.6 | pentalenene synthase (4.2.3.7) (1.10.600.10)<br>alginase (4.2.2.3) (1.50.10.100)<br>phosphohexomutase (5.3.1.8) (2.60.120.10)<br>5 (2.1.2.2) (3.40.50.170)<br>PPIase (5.2.1.8) (3.10.50.40)<br>hyaluronate lyase (4.2.2.1) (1.50.10.100)<br>meso-2 (4.2.1.28) (3.20.20.350) | 14.3 | 0 | 22.4 | 42.9 | 28.6 | 28.6 | K (16.36%)<br>CA (5.45%) | | ASG (12.73%)<br>PGO (12.73%)<br>GCT (5.45%)<br>BDP (5.45%)<br>NAG (5.45%)<br>GCU (1.82%)<br>GAR (1.82%)<br>PVC (1.82%)<br>IDR (1.82%)<br>MAN (1.82%)<br>Other (14.56%) |
| **H-R-R** | 1 | 2 | 1.2 | aspartate---tRNA ligase (6.1.1.12) (3.30.930.10)<br>phosphomonoesterase (3.1.3.2) (3.40.50.1240) | 0 | 0 | 31 | 0 | 50 | 50 | | | VO4 (12.5%)<br>PO4 (12.5%)<br>TLA (12.5%)<br>TRS (12.5%)<br>2BF (12.5%)<br>MLA (12.5%) |
| **H-R-STY** | 1 | 4 | 1.3 | swivelase (5.6.2.1) (1.20.120.380)<br>PEPPM (5.4.2.9) (3.20.20.60)<br>carbamylaspartotranskinase (2.1.3.2) (3.40.50.1370)<br>xanthan lyase (4.2.2.12) (1.50.10.100) | 0 | 0 | 31.2 | 0 | 50 | 50 | | | ASG (9.38%)<br>PCT (7.81%)<br>CP (6.25%)<br>ASP (4.69%)<br>GCT (4.69%)<br>*P* (3.12%)<br>OXL (3.12%)<br>PO4 (3.12%)<br>PAL (3.12%)<br>MLI (3.12%)<br>Other (28.08%) |
| **H-R-Z** | 1 | 2 | 1.8 | FDHH (1.17.98.4) (3.40.228.10)<br>arylsulfatase (3.1.6.1) (3.40.720.10) | 30 | 50 | 17.5 | 100 | 0 | 0 | 6MO (13.64%) | MGD (22.73%) | DDZ (36.36%)<br>FGP (9.09%)<br>SO4 (9.09%)<br>NO2 (4.55%)<br>ALS (4.55%) |
| **H-STY-STY** | 1 | 10 | 1.9 | methylmalonyl-CoA mutase (5.4.99.2) (3.20.20.240)<br>OYE (1.6.99.1) (3.20.20.70)<br>cytochrome-b5 reductase (1.6.2.2) (2.40.30.10)<br>adenosylhomocysteinase (3.3.1.1) (3.40.50.1480)<br>8-lipoxygenase (1.13.11.40) (2.40.180.10)<br>2' (3.1.4.37) (3.90.1140.10)<br>scytalone dehydratase (4.2.1.94) (3.10.450.50)<br>UDP-sulfoquinovose synthase (3.13.1.1) (3.40.50.720)<br>caspase-3 (3.4.22.56) (3.40.50.1460)<br>[histone H3]-lysine4 N-methyltransferase (2.1.1.364)<br>(2.170.270.10) | 0.4 | 35 | 16.6 | 10 | 80 | 10 | CL (0.94%) | FMN (10.8%)<br>B12 (2.82%)<br>HEM (0.94%)<br>MLC (0.47%)<br>MCA (0.47%)<br>SCA (0.47%)<br>3CP (0.47%)<br>2CP (0.47%)<br>FNR (0.47%)<br>FAD (0.47%)<br>Other (0%) | *P* (51.64%)<br>FMN (3.76%)<br>HBA (3.29%)<br>07V (1.88%)<br>0WV (1.88%)<br>5AD (0.94%)<br>TNF (0.94%)<br>NCA (0.94%)<br>3RN (0.94%)<br>BFS (0.94%)<br>Other (8.93%) |
| **H-STY-X** | 1 | 5 | 1.5 | PUNPI (2.4.2.1) (3.40.50.1580)<br>AANAT (2.3.1.87) (3.40.630.30)<br>FabD (2.3.1.39) (3.40.366.10)<br>Unknown (3.1.1.-) (3.40.50.1110)<br>2' (3.1.4.37) (3.90.1140.10) | 0 | 0 | 12.4 | 0 | 100 | 0 | | | SO4 (6.9%)<br>PO4 (5.17%)<br>DXX (5.17%)<br>IMH (3.45%)<br>DIH (3.45%)<br>ACT (3.45%)<br>DSJ (1.72%)<br>R1P (1.72%)<br>DTS (1.72%)<br>CTN (1.72%)<br>Other (6.88%) |
| **H-STY-X-X** | 1 | 3 | 1.4 | destabilase (3.5.1.44) (3.40.50.180)<br>Unknown (3.4.21.-) (3.30.750.44)<br>PHP (3.9.1.3) (3.50.20.20) | 0 | 0 | 33.3 | 0 | 100 | 0 | | | PO4 (75.0%)<br>CHM (7.14%)<br>*P* (7.14%)<br>DKT (3.57%)<br>FMT (3.57%)<br>SO4 (3.57%) |
| **H-X-X** | 1 | 5 | 1.6 | pyruvate (2.7.9.1) (3.30.1490.20)<br>caspase-3 (3.4.22.56) (3.40.50.1460)<br>rhodanese (2.8.1.1) (3.40.250.10)<br>caspase-1 (3.4.22.36) (3.40.50.1460)<br>DBP2 (3.6.4.13) (2.40.10.10) | 0.4 | 0 | 13.6 | 0 | 80 | 20 | CL (0.44%)<br>ZN (0.44%) | | *P* (24.89%)<br>0QE (23.14%)<br>ASJ (3.93%)<br>DTT (2.18%)<br>PJE (2.18%)<br>010 (2.18%)<br>ECC (2.18%) |

| | | | | | | | | | | | | | |
|---|---|---|---|---|---|---|---|---|---|---|---|---|---|
| | | | | | | | | | | | | | 1U8 (1.75%)<br>0JU (1.75%)<br>HSV (1.31%)<br>Other (33.35%) |
| **H-X-X-X** | 1 | 2 | 0.3 | caspase-3 (3.4.22.56) (3.40.50.1460)<br>caspase-3 (3.4.22.56) (3.40.50.1460) | 0 | 0 | 28.5 | 0 | 100 | 0 | | | *P* (64.63%)<br>0QE (8.54%)<br>MX4 (2.44%)<br>PZN (2.44%)<br>Y2Y (2.44%)<br>158 (2.44%)<br>161 (2.44%)<br>NA3 (1.22%)<br>XVE (1.22%)<br>CNE (1.22%)<br>Other (8.54%) |
| **K-NQ-R** | 1 | 2 | 1.5 | 3-amino-5-hydroxybenzoate synthase (4.2.1.144) (3.40.640.10)<br>2 (4.1.1.64) (3.90.1150.10) | 0 | 34.5 | 22 | 0 | 100 | 0 | | PLP (15.0%)<br>PMP (5.0%) | PXG (5.0%)<br>5PA (5.0%)<br>EPC (5.0%)<br>HCP (5.0%)<br>ELP (5.0%)<br>MPM (5.0%)<br>DCS (5.0%)<br>LCS (5.0%)<br>NMA (5.0%) |
| **K-NQ-STY** | 2 | 4 | 1.1 | DHODH (1.3.5.2) (3.20.20.70)<br>cyclohydrolase (3.5.4.9) (3.40.50.10860)<br>L-DEX (3.8.1.2) (3.40.50.1000)<br>beta-hydroxyoctanoyl-ACP-dehydrase (4.2.1.59) (3.40.50.720) | 0 | 6.8 | 47.5 | 0 | 75 | 25 | | FMN (18.18%)<br>FNR (1.52%) | ORO (66.67%)<br>DOR (4.55%)<br>FMT (2.27%)<br>L34 (1.52%)<br>KUN (0.76%)<br>KUK (0.76%)<br>9L9 (0.76%)<br>MTX (0.76%)<br>2OP (0.76%)<br>NDP (0.76%)<br>Other (0%) |
| **K-R-R** | 1 | 5 | 1.9 | pyruvate (2.7.9.1) (3.30.1490.20)<br>NADPH-sulfite reductase (1.8.1.2) (3.30.413.10)<br>swivelase (5.6.2.1) (3.90.15.10)<br>myokinase (2.7.4.3) (3.40.50.300)<br>pectolyase (4.2.2.10) (2.160.20.10) | 4.2 | 20 | 61.4 | 0 | 40 | 60 | CL (3.4%) | SRM (16.33%) | AP5 (35.37%)<br>ADP (12.24%)<br>PO4 (4.76%)<br>AMP (4.08%)<br>SO4 (2.72%)<br>SO3 (2.72%)<br>C5P (2.72%)<br>NO2 (1.36%)<br>*P* (1.36%)<br>AF3 (1.36%)<br>Other (8.84%) |
| **K-R-STY** | 1 | 7 | 1.5 | PEPK (4.1.1.49) (3.90.228.20)<br>AlaDH (1.4.1.1) (3.40.50.720)<br>pyruvate kinase (2.7.1.40) (3.20.20.60)<br>phosphohexomutase (5.3.1.8) (2.60.120.10)<br>trans (4.2.3.6) (1.10.600.10)<br>AvDH1 (3.6.4.12) (3.40.50.300)<br>duplicase (2.7.7.7) (2.30.40.20) | 1.9 | 0 | 51.3 | 14.3 | 0 | 85.7 | MN (2.99%) | | OXL (22.39%)<br>PYR (13.43%)<br>POP (8.96%)<br>THJ (5.97%)<br>ATP (5.97%)<br>ADP (5.97%)<br>GOL (4.48%)<br>CO2 (2.99%)<br>OXD (2.99%)<br>FLC (2.99%)<br>Other (20.9%) |
| **K-STY-STY** | 2 | 11 | 1.4 | galactowaldenase (5.1.3.2) (3.40.50.720)<br>chalcone isomerase (5.5.1.6) (3.50.70.10)<br>dTDP-glucose 4 (4.2.1.46) (3.40.50.720)<br>Unknown (1.1.1.30) (3.40.50.720)<br>4-hydroxy-tetrahydrodipicolinate synthase (4.3.3.7) (3.20.20.70)<br>17beta (1.1.1.62) (3.40.50.720)<br>(R)-oxynitrilase (4.1.2.10) (3.50.50.60)<br>Glu-AdT (6.3.5.7) (3.90.1300.10)<br>ATP:polynucleotidylexotransferase (2.7.7.19) (1.10.1410.10)<br>D-cysteine desulfhydrase (4.4.1.15) (3.40.50.1100)<br>ADP-glyceromanno-heptose 6-epimerase (5.1.3.20) (3.40.50.720) | 0 | 32.7 | 14.8 | 0 | 72.7 | 27.3 | | NAD (36.65%)<br>NAP (6.76%)<br>NDP (2.85%)<br>NAI (1.42%)<br>NAE (0.36%)<br>NDC (0.36%)<br>NAQ (0.36%)<br>PMP (0.36%) | NAD (8.19%)<br>NDP (6.41%)<br>NAP (3.91%)<br>DAU (1.78%)<br>3AT (1.78%)<br>NAI (1.42%)<br>GDD (1.07%)<br>EMO (1.07%)<br>GLN (1.07%)<br>TDX (0.71%)<br>Other (18.97%) |

| | | | | | | | | | | | | | |
|---|---|---|---|---|---|---|---|---|---|---|---|---|---|
| **K-STY-X** | 2 | 6 | 1.5 | beta-lactamase (3.5.2.6) (3.40.710.10)<br>glutaminase (3.5.1.2) (1.10.1500.10)<br>Unknown (3.1.1.-) (3.40.710.10)<br>SPC (3.4.21.89) (2.10.109.10)<br>ingensin (3.4.25.1) (3.60.20.10)<br>HslUV (3.4.25.2) (3.60.20.10) | 0.3 | 0 | 18.5 | 0 | 100 | 0 | NA (0.57%)<br>K (0.57%) | | DOD (5.11%)<br>SO4 (4.55%)<br>BO2 (4.55%)<br>NXL (3.41%)<br>GLU (3.41%)<br>04C (3.41%)<br>PO4 (2.84%)<br>*P* (2.84%)<br>EPE (2.27%)<br>3BV (2.27%)<br>Other (59.81%) |
| **K-X-X** | 1 | 3 | 1.3 | phosphooxymethylpyrimidine kinase (2.7.4.7) (3.40.1190.20)<br>repressor LexA (3.4.21.88) (2.10.109.10)<br>isopentenyl phosphate kinase (2.7.4.26) (3.40.1160.10) | 0 | 0 | 40.7 | 0 | 33.3 | 66.7 | | | ACP (9.09%)<br>ADP (9.09%)<br>ATP (9.09%)<br>IP8 (9.09%) |
| **NQ-R-STY** | 1 | 2 | 1.4 | NAD+-malic enzyme (1.1.1.38) (3.40.50.10380)<br>uricase (1.7.3.3) (3.10.270.10) | 1 | 17.5 | 79.5 | 0 | 100 | 0 | CL (1.71%) | NAD (5.13%)<br>NAP (0.85%) | AZA (39.32%)<br>URC (6.84%)<br>AZI (5.13%)<br>OXY (4.27%)<br>OXL (3.42%)<br>TTN (3.42%)<br>MUA (3.42%)<br>MAK (2.56%)<br>XDS (2.56%)<br>IUP (2.56%)<br>Other (17.9%) |
| **NQ-STY-STY** | 1 | 3 | 1.4 | DHODH (1.3.5.2) (3.20.20.70)<br>DD-peptidase (3.4.16.4) (3.40.710.10)<br>aspartase (4.3.1.1) (1.20.200.10) | 0 | 0 | 43 | 0 | 66.7 | 33.3 | | | ORO (52.83%)<br>HE0 (1.89%)<br>2PB (1.89%)<br>BSA (1.89%)<br>REY (1.89%)<br>0MU (1.89%)<br>PNM (1.89%)<br>RE1 (1.89%)<br>CP5 (1.89%)<br>HEL (1.89%)<br>Other (11.34%) |
| **NQ-STY-X** | 1 | 3 | 1.7 | modification methylase (2.1.1.72) (3.40.50.150)<br>DD-peptidase (3.4.16.4) (1.10.3810.10)<br>2-hydroxy-6-oxohepta-2 (3.7.1.9) (3.40.50.1820) | 20.7 | 25 | 12.3 | 33.3 | 33.3 | 33.3 | MG (10.0%)<br>CL (10.0%) | SAM (10.0%)<br>SAH (10.0%)<br>SFG (10.0%) | M0E (20.0%)<br>*P* (10.0%)<br>LHI (10.0%) |
| **NQ-X-X** | 1 | 2 | 1.4 | Dtd2 (3.1.1.96) (3.50.80.10)<br>glycylpeptide N-tetradecanoyltransferase (2.3.1.97) (3.40.630.30) | 0 | 43.5 | 54 | 0 | 50 | 50 | | MYA (41.12%)<br>NHW (16.82%)<br>NHM (2.8%)<br>YNC (0.93%)<br>COA (0.93%) | *P* (22.43%)<br>MYA (5.61%)<br>D3Y (1.87%)<br>NHW (1.87%)<br>DSN (0.93%)<br>A3G (0.93%)<br>GOL (0.93%)<br>A62 (0.93%)<br>ENF (0.93%) |
| **R-R-R** | 2 | 4 | 1.2 | myokinase (2.7.4.3) (3.40.50.300)<br>UDP-sugar diphosphatase (3.6.1.45) (3.90.780.10)<br>arginine kinase (2.7.3.3) (3.30.590.10)<br>SerRS (6.1.1.11) (3.30.930.10) | 1 | 0 | 68.2 | 0 | 0 | 100 | MG (0.67%) | | AP5 (34.9%)<br>ADP (28.19%)<br>AMP (7.38%)<br>SSA (4.7%)<br>C5P (2.68%)<br>ATP (2.68%)<br>ANP (1.34%)<br>AF3 (1.34%)<br>ALF (1.34%)<br>WO4 (1.34%)<br>Other (4.69%) |
| **R-STY-STY** | 1 | 2 | 1.8 | PEPK (4.1.1.49) (3.90.228.20)<br>proton-translocating NAD(P)+ transhydrogenase (7.1.1.1) (3.40.50.1220) | 10 | 50 | 36.5 | 50 | 0 | 50 | MG (8.57%) | NAP (31.43%)<br>NDP (17.14%)<br>TXP (2.86%)<br>TAP (2.86%) | ATP (25.71%)<br>ADP (5.71%)<br>AF3 (5.71%) |
| **R-X-X** | 1 | 2 | 1.8 | formate---tetrahydrofolate ligase (6.3.4.3) (3.40.50.300)<br>deoxyuridine-triphosphatase (3.6.1.23) (2.70.40.10) | 0 | 0 | 37.5 | 0 | 0 | 100 | | | SO4 (85.71%)<br>EDO (14.29%) |

| | | | | | | | | | | | | | |
|---|---|---|---|---|---|---|---|---|---|---|---|---|---|
| **STY-STY-STY** | 1 | 3 | 1.9 | N4-(beta-N-acetylglucosaminyl)-L-asparaginase (3.5.1.26) (3.60.20.30)<br>dehydrogenase (1.1.2.3) (3.20.20.70)<br>ubiquinol oxidase (H+-transporting) (7.1.1.3) (1.20.210.10) | 0 | 29 | 11 | 0 | 66.7 | 33.3 | | FMN (68.18%)<br>FNS (4.55%)<br>FNR (2.27%) | FMN (9.09%)<br>GLY (6.82%)<br>GOL (2.27%)<br>ASP (2.27%)<br>ASN (2.27%)<br>NAG (2.27%) |
| **STY-STY-X** | 1 | 2 | 1.1 | [histone H3]-trimethyl-L-lysine36 demethylase (1.14.11.69) (2.60.120.650)<br>endothiapepsin (3.4.23.22) (2.40.70.10) | 0 | 0 | 19.5 | 0 | 0 | 100 | | | M3L (28.0%)<br>EDO (14.0%)<br>MLZ (4.0%)<br>MLY (4.0%)<br>MMK (2.0%)<br>5V1 (2.0%)<br>5U8 (2.0%)<br>5YQ (2.0%)<br>2MR (2.0%)<br>FQ5 (2.0%)<br>Other (22.0%) |
| **STY-W-X** | 1 | 2 | 1.1 | transglutaminase (2.3.2.13) (3.90.260.10)<br>CocE (3.1.1.84) (3.40.50.1820) | 0 | 0 | 11.5 | 0 | 50 | 50 | | | DBC (66.67%)<br>BEZ (22.22%)<br>PBC (11.11%) |
| **STY-X-X** | 2 | 11 | 1.3 | Unknown (4.1.1.-) (3.90.226.10)<br>beta-lactamase (3.5.2.6) (3.40.710.10)<br>Transferred to 3.3.2.9 and 3.3.2.10 (3.3.2.3) (3.40.50.1820)<br>glutaminase (3.5.1.2) (1.10.1500.10)<br>chymotrypsin (3.4.21.1) (2.40.10.10)<br>semacylase (3.5.1.11) (3.60.20.10)<br>EAS (4.2.3.61) (1.10.600.10)<br>PEPPM (5.4.2.9) (3.20.20.60)<br>D-aminopeptidase (3.4.11.19) (3.60.70.12)<br>Glu-AdT (6.3.5.7) (3.90.1300.10)<br>arsenite-transporting ATPase (7.3.2.7) (3.40.50.300) | 6.3 | 0 | 22.9 | 0 | 63.6 | 36.4 | CL (2.58%)<br>MG (2.58%)<br>ZN (0.65%)<br>F (0.65%) | | GLU (3.87%)<br>GLN (2.58%)<br>EDO (2.58%)<br>ADP (2.58%)<br>ONL (1.94%)<br>PAC (1.94%)<br>GRO (1.94%)<br>ASN (1.94%)<br>MPD (1.29%)<br>0G6 (1.29%)<br>Other (59.71%) |
| **STY-X-X-X** | 1 | 2 | 0.2 | acetylcholinesterase (3.1.1.7) (3.40.50.1820)<br>acetylcholinesterase (3.1.1.7) (3.40.50.1820) | 0 | 0 | 7.5 | 0 | 100 | 0 | | | VX (8.06%)<br>DPF (6.45%)<br>CO3 (4.84%)<br>ELT (4.84%)<br>PGE (3.23%)<br>DEP (3.23%)<br>UNX (3.23%)<br>03S (3.23%)<br>NWA (3.23%)<br>L2Y (3.23%)<br>Other (48.3%) |
| **X-X-X** | 1 | 5 | 1.2 | trypsin (3.4.21.4) (2.40.10.10)<br>Unknown (2.3.1.-) (3.30.1600.10)<br>lysine 2 (5.4.3.2) (3.20.20.70)<br>phosphooxymethylpyrimidine kinase (2.7.4.7) (3.40.1190.20)<br>Transferred to 5.6.1.3 (3.6.4.4) (3.40.850.10) | 0 | 20 | 9.6 | 0 | 0 | 80 | | PLP (1.63%) | *P* (9.76%)<br>ADP (1.63%)<br>K7J (0.81%)<br>K7I (0.81%)<br>7P0 (0.81%)<br>0G6 (0.81%)<br>SRB (0.81%)<br>907 (0.81%)<br>BBL (0.81%)<br>5JM (0.81%)<br>Other (49.41%) |
